# Microbiome diversity and host immune functions may define the fate of sponge holobionts under future ocean conditions

**DOI:** 10.1101/2021.06.20.449181

**Authors:** Niño Posadas, Jake Ivan P. Baquiran, Michael Angelou L. Nada, Michelle Kelly, Cecilia Conaco

## Abstract

The sponge-associated microbial community contributes to the overall health and adaptive capacity of the sponge holobiont. This community is regulated by the environment, as well as the immune system of the host. However, little is known about the effect of environmental stress on the regulation of host immune functions and how this may, in turn, affect sponge-microbe interactions. In this study, we compared the microbiomes and immune repertoire of two sponge species, the demosponge, *Neopetrosia compacta* and the calcareous sponge, *Leucetta chagosensis*, under varying levels of acidification and warming stress. *Neopetrosia compacta* harbors a diverse bacterial assemblage and possesses a rich repertoire of scavenger receptors while *L. chagosensis* has a less diverse microbiome and an expanded range of pattern recognition receptors and proteins with immunological domains. Upon exposure to warming and acidification, the microbiome and host transcriptome of *N. compacta* remained stable, which correlated with high survival. In contrast, the bacterial community of *L. chagosensis* exhibited drastic restructuring and widespread downregulation of host immune-related pathways, which accompanied tissue necrosis and mortality. Differences in microbiome diversity and immunological repertoire of diverse sponge groups highlight the central role of host-microbe interactions in predicting the fate of sponges under future ocean conditions.

## Introduction

Since the industrial revolution, the ocean has taken up a substantial amount of CO_2_ that has led to an increased marine inorganic carbon concentration, reduced pH, and decreased calcium carbonate saturation state (1). This global change in ocean chemistry, exacerbated by sea surface warming, can affect many organismal processes, consequently disrupting the reef population dynamics and ecosystem functioning (2). A meta-analysis of climate change-associated studies of abundant benthic groups revealed that sponges are likely winners under future climate scenarios (3).

Sponges (Porifera) are a major component of the benthic ecosystem and are responsible for many ecological processes, such as nutrient cycling, ecosystem structuring, reef consolidation, and bio-erosion (4). Sponges are generally thought to possess exceptional ecological adaptability, stemming from a complex physiology and diverse associated microbiome, that allows them to thrive even in extreme and perturbed environments (5). However, while most siliceous demosponges exhibit resistance and may even benefit from acidified ocean conditions, calcareous sponges are vulnerable when exposed to lower pH levels (6).

Poriferans forge a close relationship with diverse groups of microorganisms to form a complex structured ecosystem referred to as the holobiont (7). The microbial complement, which can constitute up to 35% of sponge biomass (8), has key roles in nutrient assimilation and metabolism, vitamin synthesis, and defense (9). However, the species-specific bacterial assemblage in sponges can undergo restructuring under drastic environmental perturbations (5). For example, elevated temperatures disrupt symbiotic functions in the demosponge, *Rhopaloeides odorabile* Thompson, Murphy, Bergquist & Evans, 1987, leading to holobiont destabilization and dysbiosis (10). While this pattern of bacterial community dynamics usually precedes mass mortalities and is widely observed in holobiont systems under stress (11), alternative trajectories of microbiome plasticity promote rapid organismal adaptation (12, 13). For example, the metagenomic profile of the microbiome of *Coelocarteria singaporensis* (Carter, 1883) at CO_2_ seeps have enhanced capacity to utilize the abundant inorganic carbon and exhibit metabolic features that are necessary for efficient carbon fixation and nitrogen metabolism in acidified conditions compared to individuals at control sites (14). Differences in the response of the bacterial community in the context of the organismal stress response determine the impact of environmental perturbations on different marine organisms (5).

The bacterial complement is shaped by various ecological selective forces acting within a holobiont. Resource limitation coupled with antagonistic interactions, such as interspecific competition and immune functions, fine-tune microbial populations that maintain holobiont homeostasis (15). However, the balance of these control mechanisms may be disrupted in disturbed conditions allowing the proliferation of opportunistic microbial taxa. The host’s innate immune system, which is involved in sensing microbial cells and activating phagocytosis, cell death, or production of antimicrobial molecules, has been shown to respond to different stress signals in marine invertebrates (16–18). For example, elevated temperature induced the coordinated expression of pattern recognition receptors (PRRs), immune-related signaling cascades, and apoptosis regulators in the demosponge, *Haliclona* (*Reniera*) *tubifera* (George & Wilson, 1919) (19). Interestingly, the immune gene repertoire varies among sponges from different taxa (20, 21) or in species with different microbiome composition (18, 22). For example, *Stylissa carteri* (Dendy, 1889), a low microbial abundance (LMA) sponge, possesses an expanded family of proteins with scavenger receptor cysteine rich (SRCR)-like domains relative to the high microbial abundance (HMA) sponge, *Xestospongia testudinaria* (Lamarck, 1815) (22). Moreover, a survey of sponge transcriptomes revealed the absence of certain immune pathway components, such as myeloid differentiation primary response 8 (*MyD88*), in the calcarean, *Sycon ciliatum* (Fabricus, 1780) (20). Distinct combinations of immune molecules may influence microbiome control in the sponge holobiont. Thus, elucidating the links between immune system functions and bacterial community structuring may provide a better understanding of inter-species differences in the tolerance of sponges to environmental stressors (23).

Here, we characterized the microbiomes and repertoire of immune-related genes in *Neopetrosia compacta* (Ridley & Dendy, 1886) (class Demospongiae, order Haplolsclerida, family Petrosiidae) and *Leucetta chagosensis* Dendy, 1913 (class Calcarea, order Clathrinida, family Leucettidae). We examined the response of these genes and of the sponge-associated microbial communities to varying stress conditions. In addition, we compared the immune gene repertoire and microbial diversity of other sponge species to elucidate common trends. We hypothesized that changes in sponge microbiome structure will correlate with changes in the expression of certain immune response genes. Our findings highlight the importance of host-microbe interactions in predicting the fate of marine sponges in the face of a rapidly changing ocean.

## Material and Methods

### Sponge sampling and culture

Six specimens each of *N. compacta* and *L. chagosensis* were collected from the Bolinao-Anda Reef Complex in Pangasinan, northwestern Philippines (16.296° N, 120.014° E), in September 2018 with permission from the Philippines Department of Agriculture (Gratuitous Permit No. 0169-19). Sponge identities were confirmed by their morphology (24) and 28S rRNA gene analyses (Fig. S1). Donor sponges were cut into twelve (≈1 cm^3^) fragments using a sterile razor and allowed to heal *in situ* for 30 days. Healed fragments were brought to the Bolinao Marine Laboratory and allowed to acclimatize for seven days in aquaria receiving flow-through seawater under ambient conditions of pH 8.0 and 28°C.

### Stress response experiments

Stress response experiments were conducted in independently aerated 10L aquaria with flow-through seawater. Temperatures were regulated using 300W submersible heaters, levels of injected CO_2_ manipulated with mass flow controller, and the illumination followed a 12:12 light: dark photoperiod using daylight LED lamps. Conditions were designed to simulate the present day and predicted 2100 Representative Concentration Pathway (RCP) 6.0 and 8.5 scenarios (25). Treatment conditions included (i) pH 8.0, 28°C (Present Day), (ii) pH 7.6, 28°C (Acidification), (iii) pH 8.0, 32°C (Warming), (iv) pH 7.8, 30°C (RCP 6.0), and (v) pH 7.6, 32°C (RCP 8.5). Each treatment was represented by four independent replicate aquaria containing three fragments of each sponge species. Temperature and pH levels were changed gradually (temperature: +1°C/day, pH: - 0.5/day) until the desired conditions were reached (Fig. S2). Treatment conditions were maintained for up to three days when the experiment was terminated because tissue necrosis had begun to manifest in some fragments. Tissues of surviving sponges were washed with UV filtered seawater, flash-frozen in liquid nitrogen, and stored at −80°C. Light and temperature in the tanks were monitored using submersible loggers (HOBO pendant, Onset Computer Corp., Bourne, MA, USA), pH was measured using a SevenGo Duo Pro pH meter (Mettler Toledo, Columbus, OH, USA), and DO and salinity were measured using a multiparameter meter (Pro 2030, YSI Inc., Yellow Springs, OH, USA). Dissolved inorganic carbon and total alkalinity (TA) were quantified using a TA Analyzer (Kimoto Electric, ATT-05, Japan). Seawater carbonate chemistry parameters were calculated from pH, TA, temperature, and salinity data using the CO_2_SYS package (Table S1) (26).

### 16S rRNA gene sequencing and analysis

Total genomic DNA was extracted from sponge tissues (three biological replicates per species for each treatment) using DNeasy PowerSoil Pro Kit (Mo Bio, Carlsbad, CA, USA). DNA concentration was quantified using a NanoDrop spectrophotometer (Thermo Fisher Scientific). DNA extracts were sent to Macrogen, South Korea, for sequencing. Bacterial 16S rRNA V3-V4 hypervariable region was amplified from the extracted DNA using barcoded primers Bakt_341F (5’-CCTACGGGNGGCWGCAG-3’) and Bakt_805R (5’-GACTACHVGGGTATCTAATCC-3’) (27). Paired-end sequencing (300 bp) was performed on the Illumina MiSeq platform following the dual-index sequencing strategy.

Sequences were processed using QIIME2 version 2019.7(28). The DADA2 package (29) was used to remove chimeric sequences and singletons, and to correct amplicon errors. The denoised forward and reverse reads were assembled into single contigs. The taxonomic assignment of processed sequences was carried out using a Naïve Bayes classifier trained on SILVA version 132 (30). The classifier was set to include V3-V4 regions of 16S rRNA genes at 99% sequence similarity. Sequence reads from chloroplasts and mitochondria were removed from the final set of Amplicon Sequence Variants (ASVs). Raw sequence reads can be accessed in the NCBI Short Read Archive database under the BioProject ID PRJNA689294.

### Microbial community composition analysis

Rarefied ASV libraries were produced through random down-sampling to the identified smallest library size. Alpha diversity indices were computed using Phyloseq (31). Community distance matrices based on Bray-Curtis dissimilarity index were estimated using vegan (32) and visualized by non-metric multidimensional scaling. ADONIS and ANOSIM tests were performed to evaluate changes in the structure and composition of bacterial communities across treatments. Responsive ASVs were described through pairwise comparisons between Present Day samples versus samples subjected to (i) Acidification, (ii) Warming, (iii) RCP 6.0, and (iv) RCP 8.5. Differentially abundant ASVs (log fold change ≥ |2|, Benjamini-Hochberg-adjusted p-value ≤ 0.1) were identified across treatments using Phyloseq-DESeq (33) implemented in R.

### Functional prediction and analyses

Phylogenetic Investigation of Communities by Reconstruction of Unobserved States (PICRUSt2) (34), installed as QIIME2 plugin, and Tax4Fun2 (35) were used to predict the functional profiles of the bacterial communities. These tools use marker genes, such as 16S rRNA, to predict community gene counts based on ASV taxon affiliations. The weighted Nearest Sequenced Taxon Index (NSTI) cut-off score was set to 2.0 to increase accuracy of PICRUSt2 predictions. The NSTI score summarizes the relatedness of the ASVs in the sample to the closest available reference genome and serves as a basis for assessing the quality of prediction. Low NSTI values indicate higher similarity to the reference genomes and, thus, more accurate functional gene prediction (34). KEGG ortholog (KO) prediction and abundance estimation was performed and associated high-level functions were determined using the pathway_pipeline.py script with a KEGG pathways mapping file. In Tax4Fun, representative sequences were searched against the Ref100NR database to find the closest reference genome using NCBI blast+. Thereafter, KO counts and KEGG pathway profiles were predicted using the makeFunctionalPrediction command.

Changes in the predicted functional potential of the bacterial communities were described through pairwise comparisons between Present Day samples versus samples subjected to the other treatments. Differentially abundant KOs (Benjamini-Hochberg-adjusted p-value ≤ 0.05) were identified across treatments using Phyloseq-DeSeq (33) implemented in R. The combined set of differentially abundant KOs across treatments was searched against the KEGG database using the KEGG mapper-search pathway mapping tool. The relative abundance of the retrieved KEGG pathways was then calculated from the sum of the relative abundance of the associated KOs.

### Transcriptome sequencing, assembly, and annotation

Total RNA was extracted from *N. compacta* and *L. chagosensis* samples using TRIzol (Invitrogen, Waltham, MA, USA). Contaminating DNA was removed using TURBO DNA-free kit (Invitrogen). RNA concentration was determined using a NanoDrop spectrophotometer (Thermo Fisher Scientific, Waltham, MA, USA). The integrity of RNA extracts was evaluated using gel electrophoresis on 1% agarose in 1x TBE and the Agilent 2200 TapeStation System (Agilent Technologies, Santa Clara, CA, USA). Libraries were prepared from three samples per treatment, except for the *L. chagosensis* Warming and RCP 6.0 treatments, for which we were only able to obtain high quality RNA for two samples each. Barcoded libraries were prepared at Macrogen, South Korea, using the Truseq RNA library preparation kit (Illumina, San Diego, CA, USA). mRNA-enriched libraries were sequenced on the Illumina Novaseq 6000 platform to generate 100 bp paired-end reads.

Raw sequence reads were visualized with FastQC v0.11.8 (Babraham Bioinformatics) and trimmed using Trimmomatic v0.32 (36). Filtering included the removal of poor quality bases (quality score < 3) at the start and end of the reads, scanning the read with a 4-base sliding window, trimming if the average per-base quality is below 20, excluding reads below 36 bases long, and cutting 15 bases from the start of the reads.

*De novo* transcriptome assembly was carried out on Trinity (37). Transcripts with 90% sequence similarity were clustered and the longest representative contigs (>300bp) were retained. Reads were mapped back to the assembled transcriptomes and isoforms with zero isoform percentage (IsoPct) were removed to filter out putative misassembled transcripts. Isoforms with the highest combined IsoPct or longest length were retained for each transcript to generate a reference transcriptome for each species. The non-redundant transcriptomes of *N. compacta* and *L. chagosensis* are composed of 69 202 (N50= 1 150) and 92 629 (N50=1 475) transcripts, respectively. The quality of the assembled transcriptome is comparable to other Poriferan transcriptomes (20, 38) as assessed through Bowtie (39), Detonate (40), and Transrate (41) (Table S2). Highly expressed transcripts were also determined to have high contig length (Fig. S3). The assembly contains more than 90% of the metazoan core genes measured through BUSCO (42) with Metazoa *odb9* dataset (Table S2). Raw sequence reads were deposited in the NCBI Short Read Archive database under BioProject ID PRJNA689294. The reference transcriptomes used in this study has been deposited at DDBJ/EMBL/GenBank under the accession GIYW00000000 (*N. compacta*) and GIYV00000000 (*L. chagosensis*). The version described in this paper are the first versions, GIYW01000000 and GIYV01000000, respectively.

*N. compacta* and *L. chagosensis* peptides were predicted using the Transdecoder package in Trinity. Peptides were mapped against the UniProtKB/Swiss-Prot database (April 2020). To predict gene ontology (GO) annotations, the top Blastp hit for each sequence was used as input into Blast2GO (43), while protein domains were annotated by mapping the peptide sequences against Pfam 32.0 database (44) using HMMER v3.3 (45).

### Expression analysis

Reads were mapped to the assembled reference transcriptomes to estimate transcript abundance using RNA-Seq by Expectation Maximization (RSEM) (46) with bowtie alignment method (39). Differentially expressed transcripts were identified using the edgeR (47) package in R. Expected counts were converted to counts per million (CPM) and only genes with >10 CPM in at least two libraries were included in the analysis. Genes were considered differentially expressed if up or downregulation was greater than 4-fold relative to the controls with a Benjamini-Hochberg-adjusted p-value <1×10^−5^. Pairwise comparisons were conducted between control samples (Present Day) and samples subjected to the other treatments. Functional enrichment analysis for differentially expressed transcripts was done using the topGO package (48) in R. Only GO terms with a p-value <0.05 were considered significantly enriched. Protein-protein interactions for sponge homologs of genes involved in the human innate immune response were retrieved from the STRING v.11 database (49). Interaction networks were visualized using Cytoscape v.3.7.2 (50). Relative expression of sponge gene homologs in each treatment relative to the Present Day control was computed as the average sum of log2 transformed transcripts per million (TPM).

### Comparative analysis of bacterial communities and predicted metagenomes

Selected datasets from the Sponge Microbiome Project (51) were retrieved from the Qiita database under Study ID: 10 793 (March 2020). Sequences from healthy adult individuals viz. Demospongiae (n = 1441), Calcarea (n = 20), Homoscleromorpha (n = 41), and Hexactinellida (n = 2) were included in the analysis. Alpha diversity indices were computed with Phyloseq (31). The predicted functions of 128 sponge microbiomes with known LMA-HMA status viz. Demospongiae-LMA (n = 69), Demospongiae-HMA (n = 48), Calcarea-LMA (n = 4), Homoscleromorpha-LMA (n = 1), Homoscleromorpha-HMA (n = 5), Hexactinellida-LMA (n = 1), as described in a previous study (52), were shared by Miguel Lurgi (CNRS-Paul Sabatier University, France).

### Comparative analysis of sponge immunological repertoire

Predicted peptide sequences of representative demosponges (*Amphimedon queenslandica* Hooper & van Soest, 2006 (53), *H. tubifera* (38), *Petrosia* (*Petrosia*) *ficiformis* (Poiret, 1789) (20)), calcareans (*S. ciliatum, Leucosolenia complicata* (Montagu, 1814) (54)), and homoscleromorph (*Oscarella carmela* Muricy & Pearse, 2004) were annotated against the UniProtKB/Swiss-Prot (April 2020) and Pfam 32.0 (44) database. Sponge peptide sequences were downloaded from Compagen (55), except for *A. queenslandica*, which was retrieved from Ensembl Metazoa, and *P. ficiformis*, which was shared by Ana Riesgo (Natural History Museum, London).

NACHT domain-containing genes, with bona fide or tripartite NLR gene architecture (56), were identified from the sponge predicted peptides. Amino acid sequences corresponding to the NACHT domain (PF05729) were used for phylogenetic comparisons. Multiple sequence alignment was performed using Clustal Omega (57) and the aligned sequences were manually trimmed. The best-fit substitution model (LG+G+F) was identified based on Bayesian Information Criterion using prottest v3.4.2 (58). Bayesian inference analysis was performed in MrBayes v.3.2 (59) with two-independent MCMC runs and four chains per run. The analysis was sampled every 100 trees until the average standard deviation of split frequencies was <0.01. The first 25% of trees were discarded as burn-in.

### Statistical analyses and visualization

Community alpha diversity values and expression levels of PRRs were tested for normality using Shapiro-Wilk test and homogeneity of variance through Levene’s test. Statistical differences were calculated using Welch’s t-test or Wilcoxon test with p-values <0.05 considered statistically significant. KEGG functions and immunological domains that distinguished among sponge groups were identified using Linear Discriminant Analysis effect size (LDA-LEfSe) (60) based on relative abundance values. All visualizations were done using the ggplot2 package (61) in R.

## Results and Discussion

### Sponges exhibit differential survival under ocean warming and acidification

Fragments of the siliceous sponge, *N. compacta* (Fig. 1A), remained healthy under variable levels of pH and temperature stress, as well as to combinations of these two stressors (Fig. S4). In contrast, the calcareous sponge, *L. chagosensis* (Fig. 1B), showed visible tissue necrosis under the warming, RCP 6.0, and RCP 8.5 conditions, but not in the acidification only treatment (Fig. S4). While up to 97% of *N. compacta* fragments survived the most extreme condition at RCP 8.5, only 25% of the *L. chagosensis* fragments survived in the RCP 8.5 treatment after just two days of sustained exposure (Fig. 1C). These observations are comparable to the findings of other studies that reported the high survivorship of demosponges and the susceptibility of calcareous sponges subjected to these climate change-associated stressors (19).

**Fig.1.**
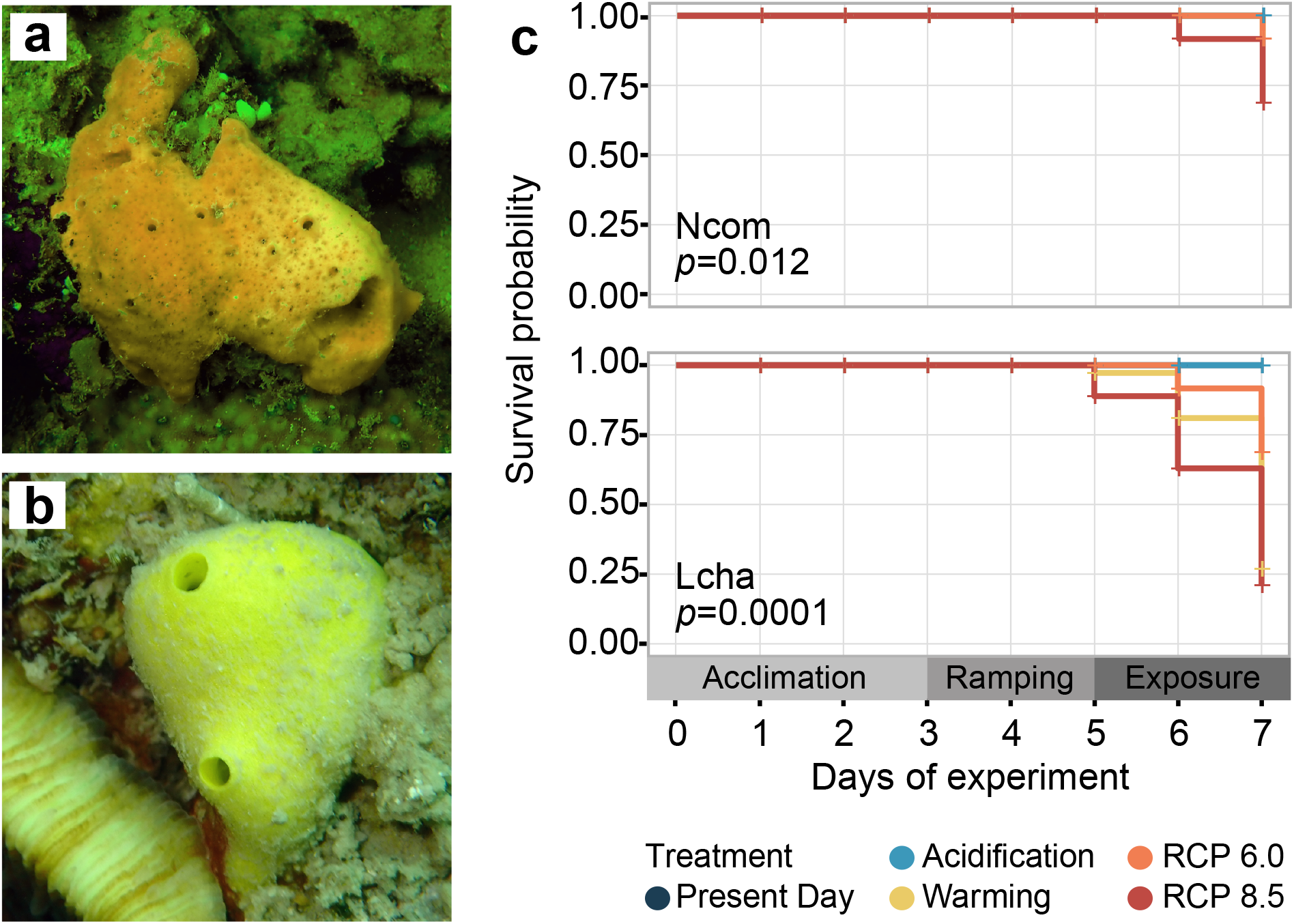
Sponge holobiont features and survivorship under future ocean conditions. Representative images of (A) *Neopetrosia compacta* and (B) *Leucetta chagosensis* in their natural habitats. (C) Survival probability of *N. compacta* (top) and *L. chagosensis* (bottom) throughout the duration of the experiment visualized using Kaplan-Meier survival analysis.

### Bacterial community shifts in the stress response of sponges

Rarefaction curves indicate the completeness of detected ASVs present in *N. compacta* (Fig. S5A) and *L. chagosensis* (Fig. S5B) across the different treatments. *Neopetrosia compacta* is associated with a fairly diverse and heterogenous bacterial assemblage (species richness = 188.00±27.78; Shannon diversity index = 3.68±0.02) with enrichment of the Chloroflexi-related SAR202 clade (22.23%) and phototrophic groups, including Nostocales (8.19%) and Synechococcales (0.12%). On the other hand, *L. chagosensis* harbored a less diverse microbiome (species richness = 150.33±53.58; Shannon diversity index = 2.04±0.66) composed primarily of Oceanospirillales (56.17%) and Deltaproteobacteria SAR324 clade (27.26%) (Fig. 2A-B; Fig. S6, S7A).

**Fig.2.**
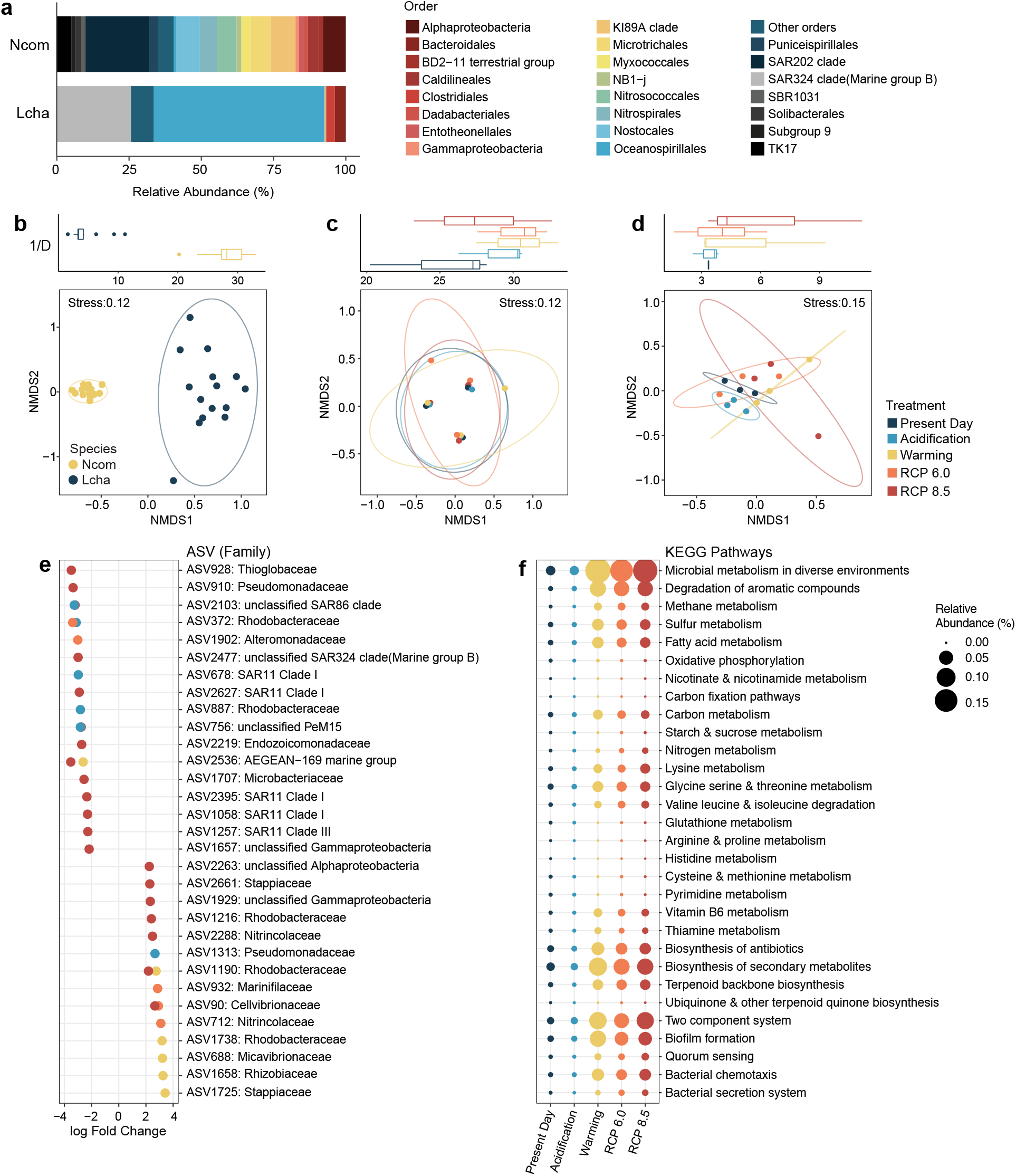
Bacterial community restructuring across stress treatments. (A) Relative abundance of major (≥ 1%) bacterial Orders in the *Neopetrosia compacta* and *Leucetta chagosensis* microbiomes. (B) NMDS clustering of sponge-associated microbial communities. Diversity and structure of (C) *N. compacta* and (D) *L. chagosensis* microbiomes under variable stress conditions. Graphs show NMDS clustering of samples. Box plots of Simpson diversity index (1/D) are shown above each graph. Colors represent different treatments (blue, Present Day; skyblue, Acidification; yellow, Warming; orange, RCP 6.0; red, RCP8,5). (E) Plot of differentially abundant ASVs in *L. chagosensis* relative to the Present Day samples. (F) Bubble plot of differentially enriched KEGG pathways in *L. chagosensis* relative to the Present Day samples. Bubble size indicates relative abundance.

To explore the possible roles of bacterial community dynamics in the stress response of the two sponge holobionts, we described the shifts in the taxonomic profiles of their bacterial associates following exposure to simulated conditions. The microbial community of *N. compacta* showed little change in structure when subjected to the various stressors (Fig. 2C). The few ASVs that showed a significant change in abundance relative to the present day treatment include ASV749 (Microbacteriaceae), which decreased in Acidification and RCP 6.0 conditions, and ASV842 (Endozoicomonadaceae), ASV2840 (Cellvibrionaceae), ASV73, ASV1088, ASV887, ASV2836, ASV1181, and ASV2292 (Rhodobacteraceae), ASV2836 (Alteromonadaceae), ASV1181 (Nitrincolaceae), and ASV2292 (Colwelliaceae), which increased in RCP 6.0 (Fig. S8).

In contrast, the bacterial assemblage of *L. chagosensis* exhibited apparent restructuring with the treatments, although not statistically supported (Fig. 2D, Table S3). A total of 37 ASVs exhibited changes in relative abundance, with 21 decreasing and 16 increasing (Fig. 2E). In the Acidification treatment, where 100% of the sponges survived, a decrease in Bacteroidales and Clostridiales and an increase in the most dominant symbiont, Oceanospirillales, was observed (Fig. S9B). On the other hand, in the treatments with high sponge mortality (i.e. Warming, RCP 6.0, and RCP 8.5), there was reduced abundance of ASV2477 (SAR324 clade) and ASV2219 (Endozoicomonadaceae). Presumptive opportunistic taxa, such as Vibrionales (ASV90, ASV688), Rhodobacterales (ASV1216, ASV1190, ASV1738, ASV2661, ASV1725), and Rhizobiales (ASV1658) (62), increased in relative abundance under these treatments (Fig. 2E; Fig. S9B).

Taxa that proliferated in the microbial community of *L. chagosensis* under stress conditions were predicted to invest more in the production of antimicrobial molecules (Fig. 2F), which may be advantageous in competitive colonization of the tissues of the sponge. The increased abundance of opportunistic taxa may be due to their capacity to form biofilms for efficient surface adhesion and active secretion of virulence factors to invade the host cell (63). These traits, along with the ability to sense and respond to environmental perturbations, as evidenced by enrichment of functions related to two-component signaling and bacterial chemotaxis, may support successful proliferation of certain taxa that will eventually outcompete other microbiome members (64). The resulting large-scale changes in the bacterial community of *L. chagosensis* are predicted to correlate to a shift in the functional and metabolic potential of the holobiont, which may further contribute to the decline of the host.

### Sponge immune response under ocean warming and acidification conditions

To determine how the host innate immune system is affected by acidification and warming, we described the expression patterns of immune-related genes in *N. compacta* and *L. chagosensis* using RNA-Seq. A total of 1 596 genes (Acidification = 74, Warming = 308, RCP 6.0 = 501, RCP 8.5 = 713) were found to be differentially expressed in *L. chagosensis*, whereas only 70 genes (Acidification = 3, Warming = 7, RCP 6.0 = 12, RCP 8.5 = 48) genes were differentially expressed in *N. compacta*.

Although reduced environmental pH levels have been reported to induce bacterial virulence (65), gene ontology (GO) enrichment analysis of differentially expressed genes in the calcareous sponge revealed that *L. chagosensis* is able to mount relevant responses, including endosome organization and antibacterial humoral response to avoid pathobiont invasion, under Acidification treatment (Fig. 3A). On the other hand, key defense mechanisms against microbial perturbations were differentially regulated in *L. chagosensis* under Warming, RCP 6.0, and RCP 8.5 conditions. Specifically, genes implicated in recognition of microbe-associated molecular patterns (MAMPs) and receptor-mediated endocytosis were repressed in *L. chagosensis*. In particular, scavenger receptors (SRCRs), secretin G-protein coupled receptors (GPCRs), and nucleotide-binding domain and leucine-rich repeat-containing genes (NLRs), exhibited reduced expression under Warming, RCP 6.0, and RCP 8.5 conditions (Fig. 3B). The reduction in the expression levels of these genes and the sensor proteins that they encode, along with the repression of bactericidal permeability-increasing protein (*BPI*) and lipopolysaccharide binding protein (*LBP*) (Fig. 3C), may result in impaired recognition of microbial cells or molecules, which, in turn, influences the regulation of downstream effectors of the immune response (66). Components of other principal machineries involved in the response to pathogen infection, such as autophagy, inflammation, and apoptosis, were similarly downregulated (Fig. 3A).

**Fig.3.**
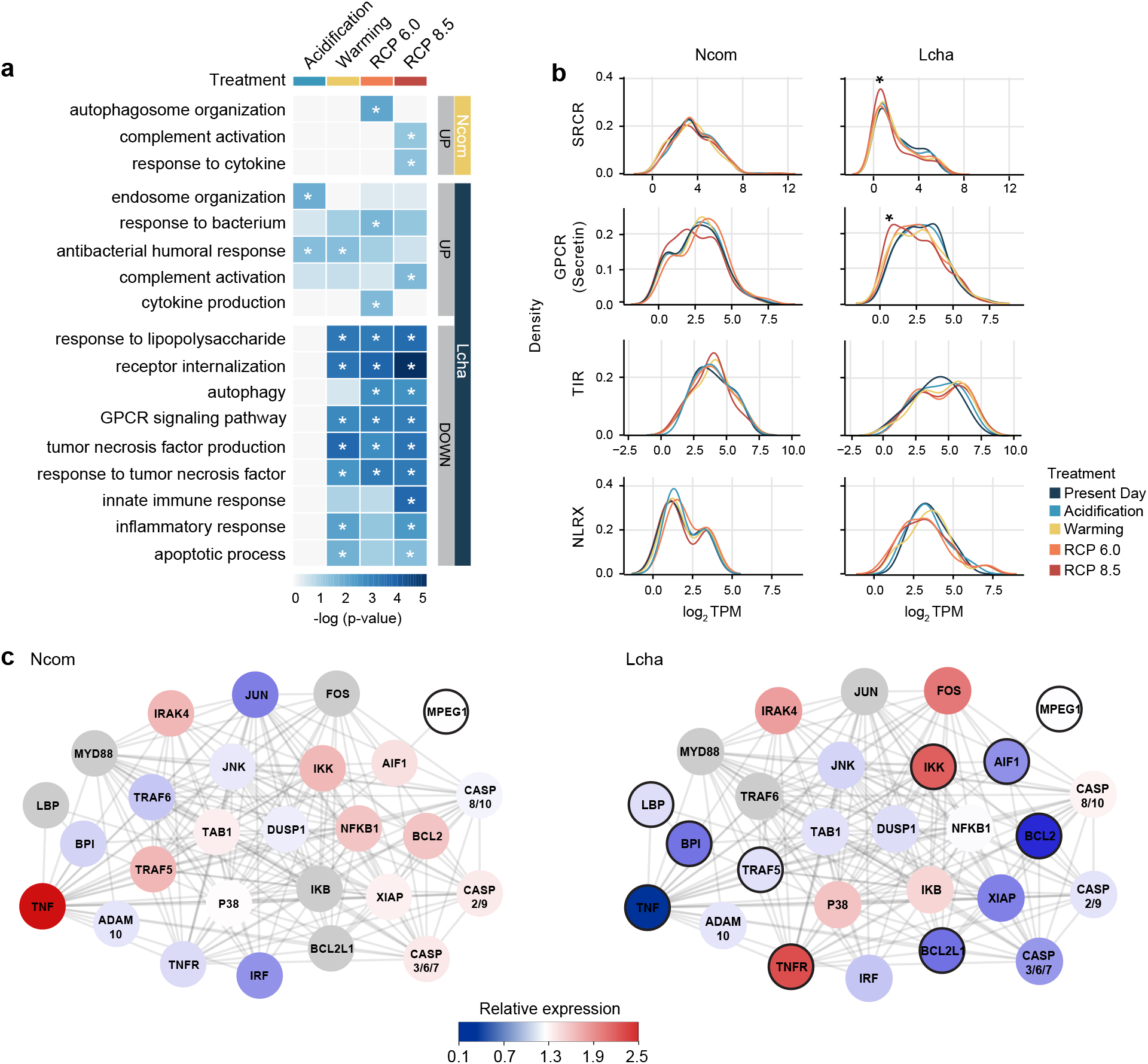
Immune responses to different treatment conditions. (A) Gene Ontology (GO) enrichment analysis for the up and downregulated transcripts in *Neopetrosia compacta* and *Leucetta chagosensis* under the different treatments. Only immune-related GO terms are presented. (B) Expression levels of major pattern recognition receptors are presented as the log2 transformed TPM values in *N. compacta* (left) compared with *L. chagosensis* (right). NLRX refers to NACHT-containing genes with bona fide NLR architecture. Colors represent different treatments. Asterisks indicate significant change (*p*<0.05) in expression relative to the Present Day samples, as determined through Welch’s t-test or Wilcoxon test. (C) Protein interaction network of immune-related genes in *L. chagosensis* (right) and *N. compacta* (left). Relative expression of genes was computed as the sum of TPM values relative to the Present Day samples (blue, low; red, high; gray, no match). Genes with at least one differentially expressed transcript are marked by black borders. The network is based on human protein-protein interactions.

Decreased expression of genes involved in tumor necrosis factor (*TNF*) signaling suggests that *L. chagosensis* may no longer be able to deploy synchronized expression of effector molecules that mediate diverse aspects of innate immunity (67). While the TNF receptor (*TNFR*) increased in expression, the *TNF* ligand, activator disintegrin and metalloproteinase domain-containing protein 10 (*ADAM10*), and the adapter protein TNF receptor-associated factor 5 (*TRAF5*) reduced in expression (Fig. 3C). The repression of responses regulated through this pathway is further supported by the downregulation of immune-related transcription factors, such as interferon regulatory factor 5 (*IRF5*) and nuclear factor NF-kappa-B p105 subunit (*NFKB1*), coupled with the increased levels of the *NF-kB* inhibitor (*IKB*) and inhibitor of *NF-kB* kinase (*IKK*) (68). Indeed, the downregulation of macrophage-expressed gene protein 1 (*MPEG1*), allograft inflammatory factor-1 (*AIF1*), initiator caspase *CASP2/9*, and executioner caspase *CASP3/6/7*, suggest inhibited antimicrobial, inflammatory, and apoptotic mechanisms (69–71). Genes with anti-apoptotic functions, including the apoptosis regulator (*BCL2*), Bcl-2-like protein 1 (*BCL2L1*), and X-linked inhibitor of apoptosis (*XIAP*) were negatively regulated, as well. These results generally suggest that *L. chagosensis* may not be able to restore immune homeostasis under combined warming and acidification conditions.

In contrast to the calcareous sponge, the demosponge *N. compacta* exhibited activation of the complement system and cytokine-induced processes under RCP 8.5 conditions (Fig. 3A). An increased level of *TNF*, along with the upregulation of interleukin-1 receptor-associated kinase-4 (*IRAK4), TRAF5*, and *NFKB1*, indicates that the TNF-NFkB and Myd88-dependent signaling pathways were activated (72, 73) (Fig. 3C). The increased expression of *AIF1*, *CASP2/9*, and *CASP3/7*, along with the *BCL2* and *XIAP*, suggest active inflammatory and apoptotic functions (69, 71). These indicate that *N. compacta* may be able to sustain its ability for symbiont recognition and pathogen clearance through the coordinated expression of signaling pathways and immune effector mechanisms even under the most extreme conditions in this study.

Demosponges and calcareans are characterized by disparate bacterial communities and immunological repertoires

Evaluation of the diversity patterns of bacterial communities in representatives from different sponge classes showed that demosponges typically harbor bacterial communities with a wide range of taxonomic diversity, whereas calcareans are generally associated with less diverse bacterial communities (Fig. 4A). Low microbial abundance has been consistently reported among calcareans through microscopy (74), metagenomics (51), and culture-based techniques (74). Comparisons of predicted microbiome functions indicate that there is functional differentiation among HMA and LMA sponge microbiomes (Fig. 4B). Generally, the bacterial associates of calcareans, along with LMA demosponges and homoscleromorphs, are enriched for functions related to metabolism, whereas the bacterial assemblage of HMA demosponges and homoscleromorphs are enriched for functions related to general cellular processes and genetic information processing (Fig. 4B; Table S4). In particular, functions related to cofactor and vitamin metabolism, transport, and catabolism are enriched in the microbiomes of calcareans. LMA demosponge microbiomes are enriched for xenobiotic biodegradation and metabolism, terpenoid and polyketides metabolism, and lipid metabolism. In contrast, the bacterial associates of HMA demosponges and homoscleromorphs are enriched with genes involved in transcription, translation, protein processing, signal transduction, cellular community, cell motility, cell growth and death, and energy metabolism. HMA species have been reported to host bacterial communities with convergent functional potential, whereas LMA species exhibit greater microbiome differentiation (52, 75). The functional variability of LMA sponge microbiomes is likely shaped by the metabolic requirements of the holobiont under emerging environmental conditions.

**Fig.4.**
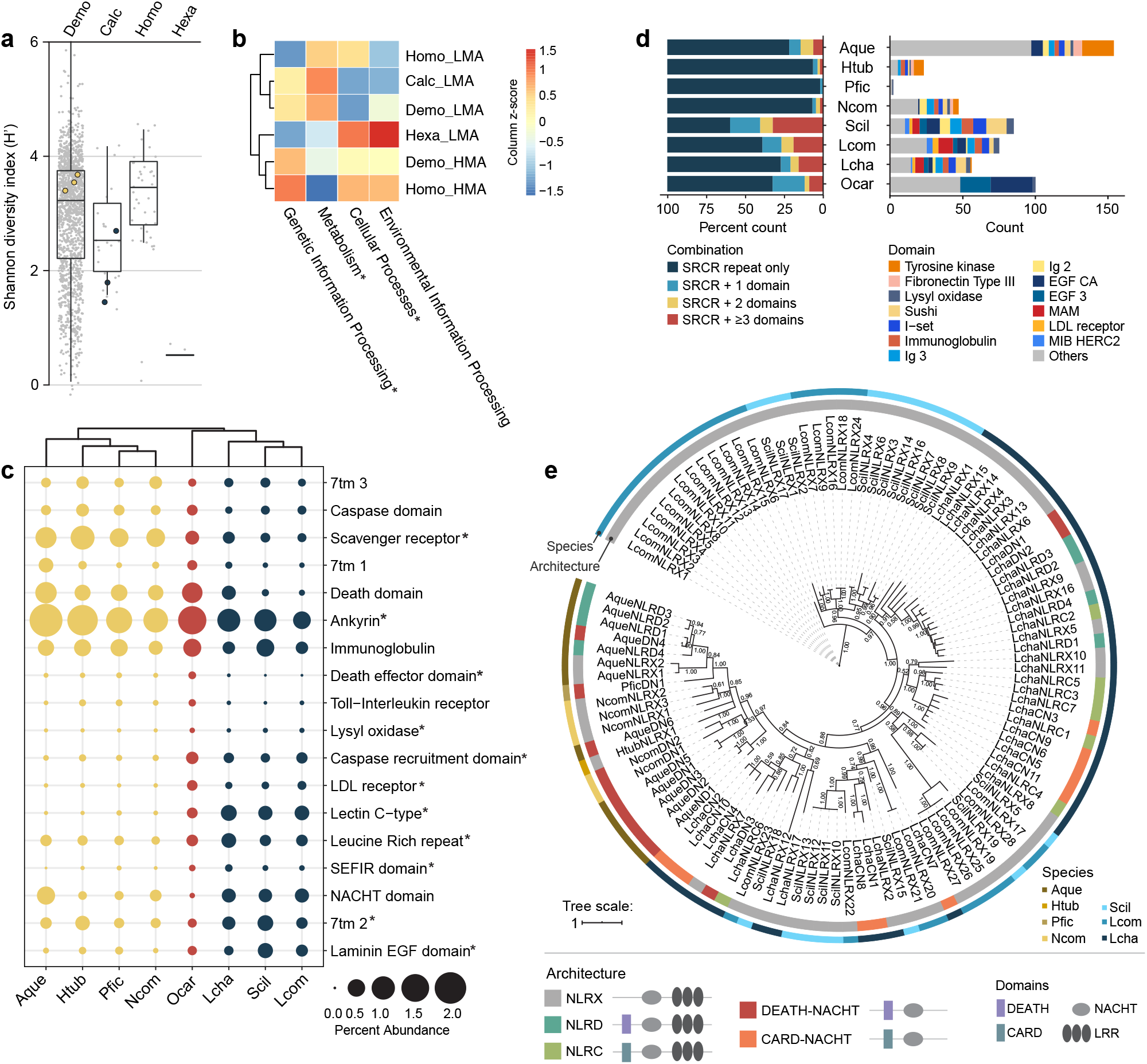
Microbiomes and immunological repertoire of sponges. (A) Taxonomic diversity (Shannon index) of bacterial communities of healthy sponge adults retrieved from the Sponge Microbiome Project. Yellow and blue circles represent diversity of the bacterial communities of *N. compacta* and *L. chagosensis*, respectively. (B) Enrichment of predicted functions (red, high; blue, low) in the microbiome associated with LMA or HMA sponges from different classes (Demospongiae, Demo; Calcarea, Calc; Homoscleromorpha, Homo; Hexactinellida, Hexa). Asterisks denote significant enrichment of functions in a specific sponge group, as determined through LDA-LEfSe. (C) Abundance of peptides containing selected Pfam domains across species. Bubble size indicates the percent of peptides containing a specific domain relative to the total number of predicted peptides in each species. Colors represent sponge class (yellow, Demospongiae; red, Homoscleromorpha; blue, Calcarea). Asterisks denote significantly higher counts of PFAM domains for a specific sponge class, as determined through LDA-LEfSe. (D) Abundance and diversity of SRCR-containing peptides across species. The percent of SRCR-containing peptides (left) and the count of SRCR peptides associated with other PFAM domains (right) are presented for each species. (E) Diversification of bona fide NLRs in Demosponges and Calcareans. The phylogenetic tree was derived from Bayesian analysis. Numbers on selected branches represent Bayesian posterior probabilities. Outer and inner color strips indicate species and peptide architectures, respectively. Species abbreviations: *Amphimedon queenslandica* (Aque), *Haliclona tubifera* (Htub), *Petrosia ficiformis* (Pfic), *Neopetrosia compacta* (Ncom), *Oscarella carmela* (Ocar), *Leucetta chagosensis* (Lcha), *Sycon ciliatum* (Scil), *Leucosolenia complicata* (Lcom).

Host phylogeny influences the diversity of the sponge microbiome (76), which may, in part, be due to species-specific immune receptor complements. Comparison of immune-related protein domains in representative sponge species revealed lineage-specific abundance patterns (Fig. 4C). SRCR domain-containing proteins were more abundant in demosponges. These domains are part of PRRs that recognize a wide array of bacterial ligands (77). SRCR-containing peptides in calcareans are associated with diverse combinations of immune or cell-adhesion domains, whereas in demosponges, the peptides consist mostly of multiple SRCR domains (Fig. 4D). Calcareans also possess a higher number of genes with secretin GPCR domains (Fig. 4C), a membrane receptor involved in sensing diverse physiological stimuli and in shaping immune responses toward extracellular pathogens and danger molecules (78).

NLRs are a group of intracellular receptors that detect foreign microbes that are able to evade extracellular defenses (79). These genes likely play a critical role in mediating host-symbiont interactions and in differentiating pathogenic from symbiotic microbes (80). Bona fide NLRs (NLRX) are characterized by both a central NACHT domain and C-terminal leucine-rich repeats (LRRs) (81). Calcareans possess an extensive family of NLRs, which group into a separate clade distinct from that of other metazoan NLRs (Fig. 4E; Fig. S10) (56). Twenty NLRX genes were identified in *S. ciliatum*, 29 in *L. complicata*, and 28 in *L. chagosensis*. In contrast, fewer NLRX genes were detected in the demosponges, with six in *A. queenslandica*, one in *H. tubifera*, and three in *N. compacta*. Among the 28 NLRX genes in *L. chagosensis*, 11 have a tripartite architecture with either a CARD (NLRC, n = 7) or DEATH domain (NLRD, n = 4) at the N-terminal. Other *L. chagosensis* NLRs that are phylogenetically related to NLRX genes possess only the central NACHT domain alongside either a CARD (CARD-NACHT, n = 11) or DEATH domain (DEATH-NACHT, n = 3). The co-expansion of tripartite NLRC, CARD-NACHT, and other CARD-containing genes in *L. chagosensis*, as well as in other calcareans is indicative of enhanced signaling potential from homotypic interactions that launch immune effector mechanisms (82). This lineage-specific expansion, coupled with the rich complement of other surface receptors, may have evolved to facilitate the maintenance of the low abundance and distinct microbiome in calcareans (52) through efficient selection or phagocytic clearance of interacting microorganisms.

### Sponge holobionts in the future ocean

Understanding the persistence of sponge holobionts in perturbed and extreme conditions requires the elucidation of the animal host, the bacterial partners, and their interactions. Our study revealed that bacterial complement diversity may define the adaptive capacities of sponge holobionts under future ocean conditions. The HMA sponge *N. compacta* exhibited greater tolerance to stress compared with LMA sponge *L. chagosensis*, which showed visible tissue necrosis and high mortality to the combined effects of warming and acidification.

The stress tolerance of *N. compacta* was supported by a stable microbiome with abundant phototrophic members (Fig. S7A, C). Other photosymbiotic sponges, *Carteriospongia foliascens* (Pallas, 1766) and *Cymbastela coralliophila* Hooper & Bergquist, 1992, have also been shown to have a higher resistance to future ocean conditions due to enhanced productivity of their cyanobacteria symbionts under elevated inorganic carbon concentration (83, 84). The notable increase in relative abundance of photoheterotrophic Rhodobacteraceae (Fig. S8) and the stable population of other photosymbionts in *N. compacta* (Fig. S9A) may have ameliorated the effects of stress from acidification and warming.

On the other hand, we propose that the susceptibility of *L. chagosensis* to stressors is linked to the instability of its microbiome, possibly stemming from low taxonomic diversity (Fig. S6) and low functional redundancy (Fig. S6; Fig. S7B) (85). While microbiome flexibility in low microbial abundant corals (86) and sponges (87) has been proposed as a mechanism for rapid adaptation (13), unstable phases during community restructuring may result in loss of essential functions and offer an opportunity for pathogen invasion. Large-scale changes in the predicted metabolic capabilities and pathogenic potential of the restructured *L. chagosensis* microbiome under stress is consistent with observations on the dysbiotic metagenomes of *R. odorabile* (10) and the coral, *Porites compressa* (88).

We speculate that the difference in bacterial community dynamics in the two sponges may be underpinned by differences in the host’s immune functions. Microbial recognition in sponges, such as in *Dysidea avara* (Schmidt, 1862) and *Aplysina aerophoba* (Nardo, 1833), involves the expression of genes encoding NLRs, SRCRs, and GPCRs, along with the activation of apoptotic functions (18). Further, recent studies provide evidence that symbiont recognition and maintenance among poriferans is mediated by TNF-NFkB dependent pathways (72, 89). Under the simulated stress conditions, sustained levels of surface receptors and expression of immune effectors in *N. compacta* may have allowed efficient symbiont recognition and pathogenic clearance, whereas the broadscale suppression of immune pathways in *L. chagosensis* may have disrupted the sponge-symbiont interactions and attenuation of the host’s defense mechanisms. Our results mirror reports on adaptive or dysbiotic events in other holobionts challenged by various environmental perturbations. For instance, the coral, *Montipora aequituberculata*, which had a stable bacterial community under elevated temperatures, exhibited regulation of the complement system and phagocytosis (90), while the dissociation of coral-algal symbiosis in *Orbicella faveolata* following a prolonged thermal anomaly was accompanied by overall reduced expression of genes implicated in the TNF pathway and apoptosis (17).

The HMA or LMA status of sponges correlates with differences in host physiology (91) and holobiont strategies for nutrient assimilation and processing (92). Although sponge pumping rates may vary across species and sponge body size (93), HMA species generally have slower filtration rates and a denser mesohyl, while LMA demosponges and calcareans can more rapidly take up large volumes of seawater through their tissues (91, 94, 95). For example, *Leucetta* can filter about 4.56L hr^−1^ (95) while *Neopetrosia problematica* (de Laubenfels, 1930) can only take up 0.53L hr^−1^ (94). Differences in pumping rate and tissue density may influence the degree of exposure to stressors. Species with higher pumping rates and lower tissue density may be more susceptible to perturbations as they may have more frequent encounters with pathobionts and are less protected against the external environment. Under elevated temperature and lowered pH conditions, sponge pumping capacity and skeletal strength may also be adversely affected, as observed in the glass sponge, *Aphrocallistes vastus* (Schulze, 1886) (96). Given their LMA status and synapomorphic calcitic spicules, calcareans are likely to be more negatively affected by future ocean conditions. It is worth noting, however, that *L. chagosensis* survived under reduced pH, which corroborates with the reported survival and proliferation of *L. complicata* at pH 7.7 (97). Indeed, elevated temperature seems to be more detrimental to sponges compared to reduced pH (83). However, the predicted co-occurrence of acidification and warming may cause the narrowing of organismal thermal tolerance thresholds (98). Surprisingly, the calcaronean sponge, *Sycettusa hastifera*, has been shown to tolerate thermo-acidic stress with little change to its microbiome and spicules (99). The resistance of *S. hastifera* to perturbed environmental conditions may be linked to its opportunistic and invasive traits (100).

Comparison of the lineage-specific patterns of microbiome diversity and immunological repertoire among poriferans provides broader insights into the adaptive capacity of different sponge groups in perturbed ocean conditions. Although sponges are generally predicted to be winners under future ocean scenarios, species and lineage-specific holobiont features may define their susceptibly or tolerance to various stress events. It is thus warranted to further investigate the roles of microbiome flexibility and immune functions in the stress response of other sponge species representing diverse groups with different evolutionary histories, morphologies, and microbiome densities. Given that sponges are critical members of the reef ecosystem, with key roles in nutrient cycling and substrate consolidation, revealing the mechanisms to their adaptive success or failure is pivotal in projecting the reef landscape in the future ocean.

## Supporting information

SuppInfo_spongeholOAW

## Acknowledgements

We thank Francis Kenith Adolfo, Robert Valenzuela, and Ronald De Guzman for field and hatchery assistance, and staff of the Bolinao Marine Laboratory for logistical support. This study was funded by the Department of Science and Technology Philippine Council for Agriculture, Aquatic and Natural Resources Research and Development (QMSR-MRRD-MEC-295-1449) to C.C.

