## Supplementary material for "Microbiome diversity and host immune functions may define the fate of sponge holobionts under future ocean conditions": SuppInfo_spongeholOAW

### List of Contents


| | Page |
|---|---|
| **Supplementary Methods** | 2 |
| **Supplementary Figures** | |
| **Fig. S1.** Phylogenetic relationships of representative sponge species | 3 |
| **Fig. S2.** Physico-chemical parameters throughout the duration of experiment | 4 |
| **Fig. S3.** Expression dependent N50 | 5 |
| **Fig. S4.** Morphological changes observed in *N. compacta* and *L. chagosensis* after sustained exposure for two days to different treatments | 6 |
| **Fig. S5.** Sequencing coverage of sponge-associated bacterial communities | 7 |
| **Fig. S6.** Richness and Shannon diversity index of ASVs and KOs of *N. compacta* and *L. chagosensis* microbiomes | 8 |
| **Fig. S7.** Distinguishing features of sponge microbiomes | 9 |
| **Fig. S8.** Differentially abundant ASVs in *N. compacta* subjected to Acidification or RCP 6.0 | 10 |
| **Fig. S9.** Taxonomic profiles of the bacterial communities of the two sponge species under different treatment conditions | 10 |
| **Fig. S10.** Phylogenetic analysis of bona fide NLRs of sponges and other metazoans | 11 |
| **Supplementary Tables** | |
| **Table S1.** Carbonate chemistry of seawater in treatment tanks | 12 |
| **Table S2.** Assembly statistics of *N. compacta* and *L. chagosensis* transcriptome | 13 |
| **Table S3.** Comparison of bacterial communities between species and across treatments | 13 |
| **Table S4.** Comparison of predicted KEGG level 2 functions of different sponge groups | 14 |
| **Table S5.** List of sponge species cited in the present study | 15 |
| **Supplementary Data** | |
| **Data S1.** Aligned and trimmed amino acid sequences corresponding to the NACHT domain (PF05729) that were included for phylogenetic comparisons of sponge NACHT domain-containing genes | 16-25 |
| **Data S2.** Aligned and trimmed 28S rRNA gene sequences of *N. compacta* and *L. chagosensis* and of representative sponge species retrieved from NCBI | 26-33 |
| **Data S3.** Aligned and trimmed amino acid sequences of bona fide NLRs of sponges and other metazoans | 34-57 |

**Supplementary Methods**

28S rRNA gene analysis

Total genomic DNA was extracted from the tissues of *Neopetrosia compacta* and *Leucetta chagosensis* using DNeasy PowerSoil Pro Kit (Mo Bio, Carlsbad, CA, USA). 28S rRNA gene was amplified from the extracted DNA using primers 28S-C2-fwd (5'-GAA AAG AAC TTT GRA RAG AGA GT-3') and 28S-D2-rev (5'- TCC GTG TTT CAA GAC GGG-3') (1). Amplified samples were sent to Macrogen, South Korea, for sequencing.

28S rRNA gene sequences of representative sponge species were retrieved from NCBI. Multiple sequence alignment of sponge 28S rRNA gene sequences was performed using Clustal Omega (2) and the aligned sequences were manually trimmed. The best-fit substitution model (GTR+G) was identified based on Bayesian Information Criterion using jModelTest v.2.1.7 (3). Phylogenetic analysis was carried out in MrBayes v.3.2 (4) with two-independent MCMC runs with four chains per run. The analysis was sampled every 100 trees until the average standard deviation of split frequencies was <0.01. The first 25% of trees were discarded as burn-in.

Morphological examination of *Neopetrosia compacta*

The voucher sponge sample forms a thick encrustation to hemispherical mass. Surface rough, like sandpaper, with slightly raised oscules, up to 10 mm diameter; texture stony; colour in life, intense turmeric yellow with a brownish yellow to reddish surface tinge in places; skeleton a very dense, spherical reticulation of robust, hastate oxeas, 234 (169–285) µm long, n=90; mucous-rich. The sponge is most closely comparable to *Neopetrosia compacta* (Ridley & Dendy, 1886) (Demospongiae, Haplosclerida, Petrosiidae) as compared to the original description from the Eastern Philippines (Ridley & Dendy 1886) and particularly, as re-described by Thiele (1912) from Aru Island.

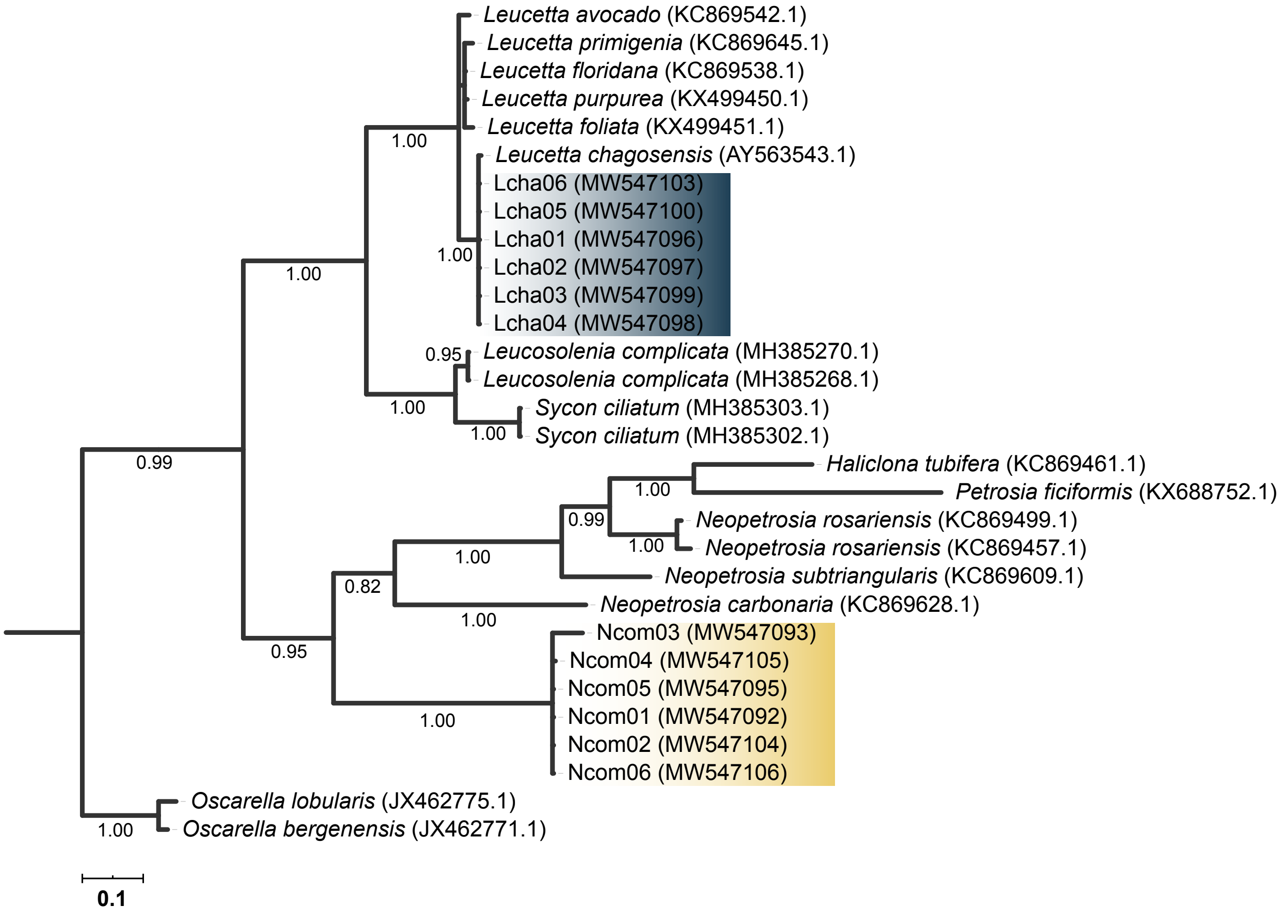


Fig. S1. Phylogenetic relationships of representative sponge species. The phylogenetic tree was derived from Bayesian analysis of 28S rRNA gene sequences. Numbers on selected branches represent Bayesian posterior probabilities. Shaded branches are sequences from the present study (yellow, *N. compacta*; blue, *L. chagosensis*).

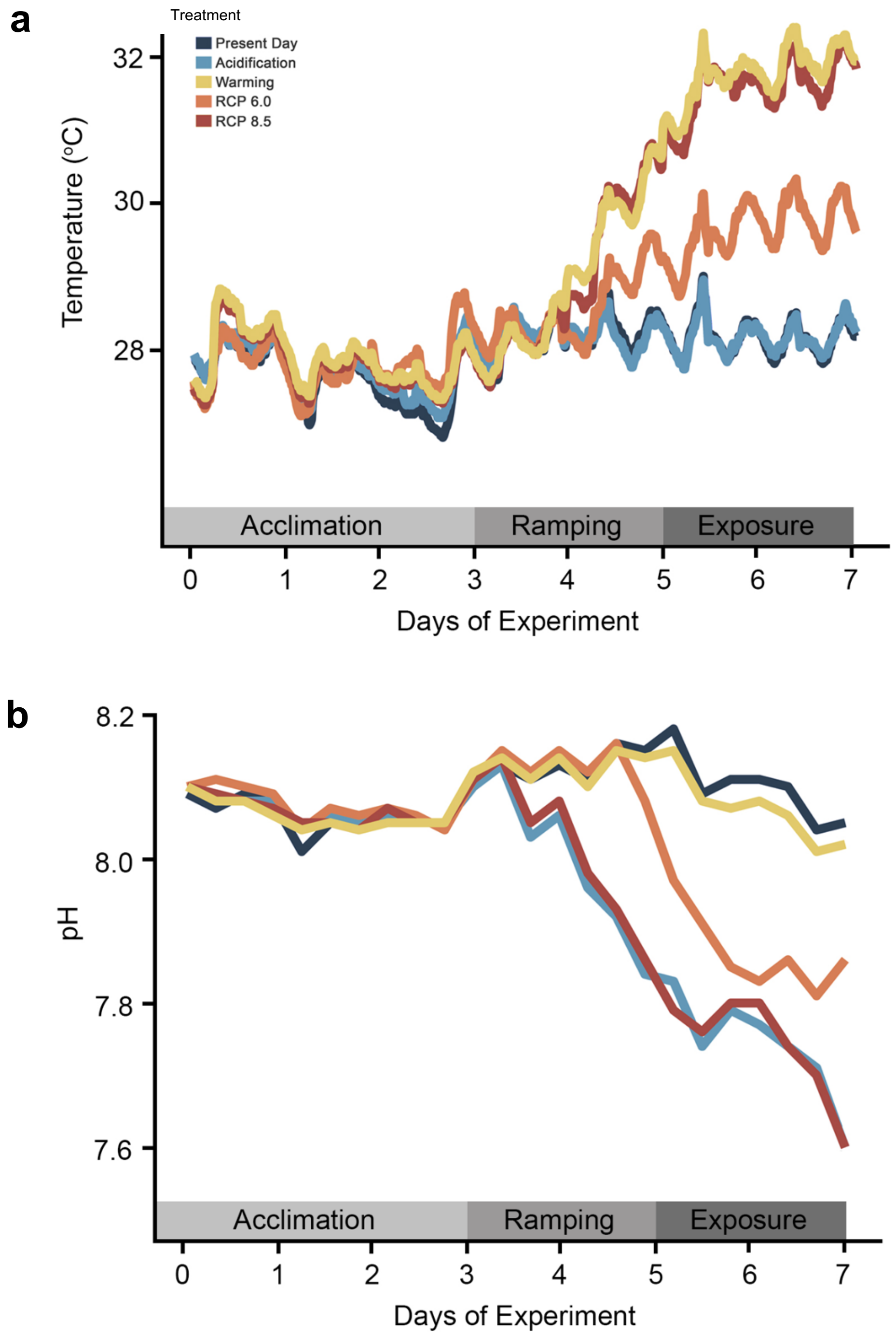


Fig. S2. Physico-chemical parameters throughout the duration of experiment. After acclimation, (A) temperatures and (B) pH levels were changed gradually until the desired conditions were reached. Experiment conditions were maintained and terminated as mortalities were observed. Different colors represent the various treatments.

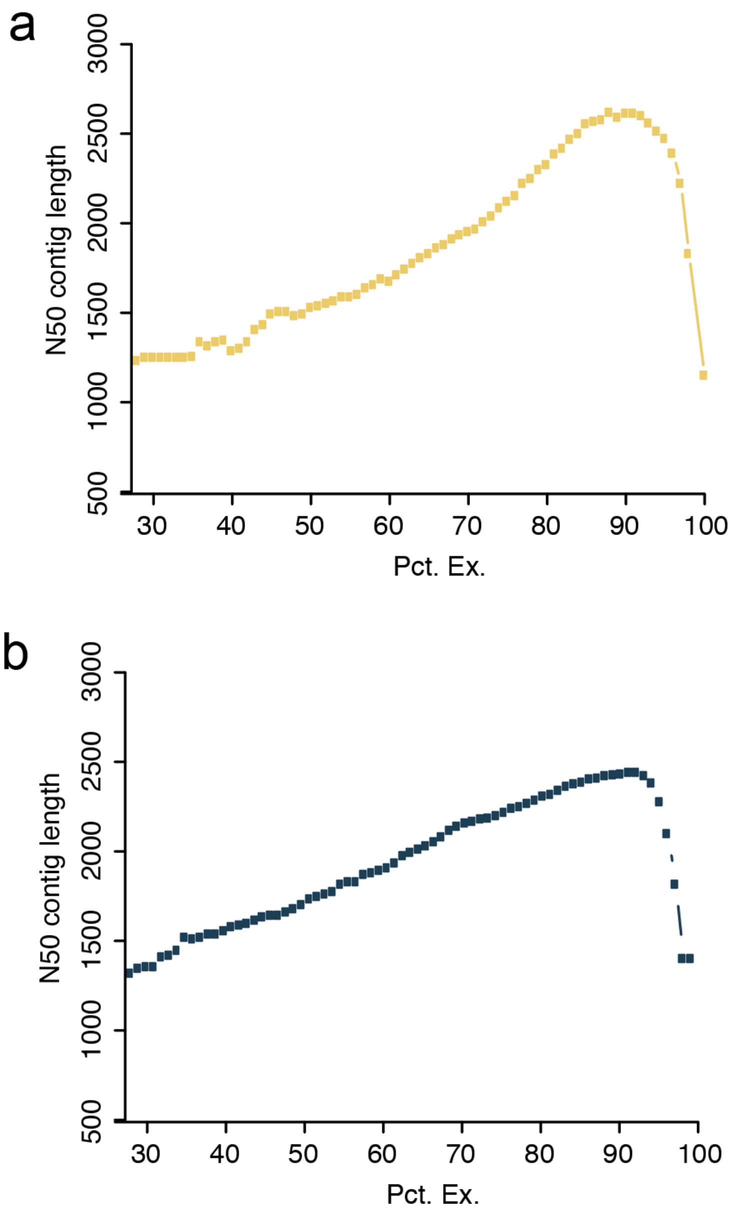


Fig. S3. Expression dependent N50 (ExN50). The plots show the fraction of the expressed data for specific N50 contig in (A) *N. compacta* and (B) *L. chagosensis* transcriptomes.

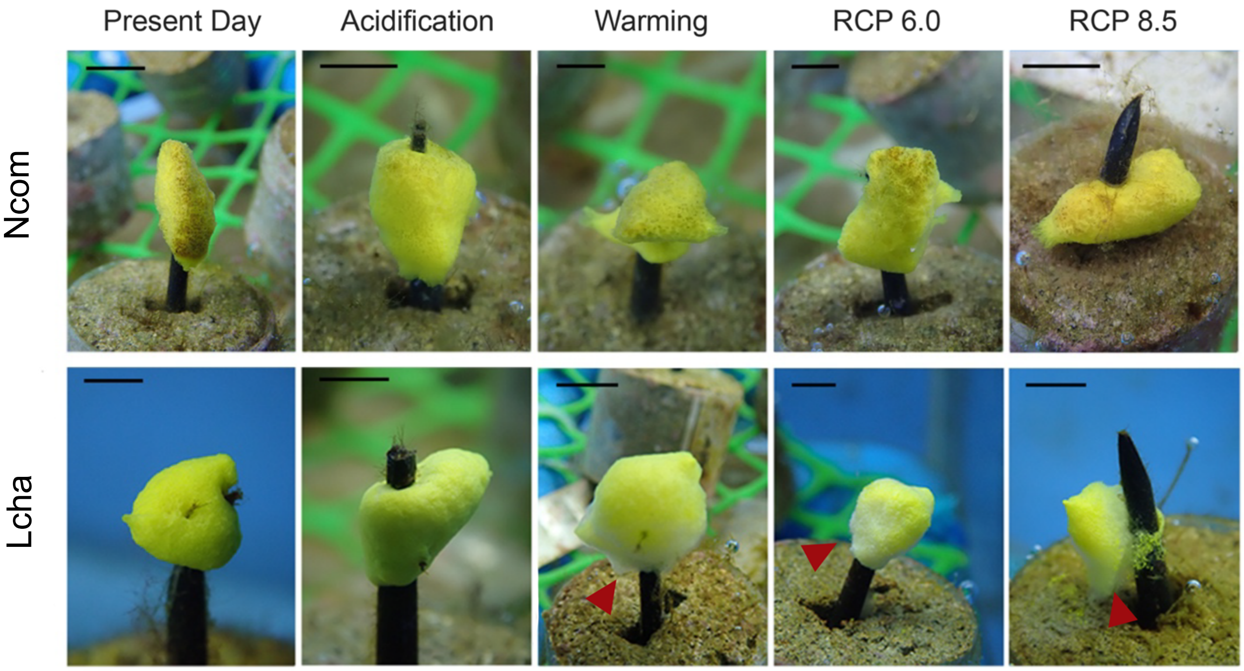


Fig. S4. Morphological changes observed in *N. compacta* (top) and *L. chagosensis* (bottom) after sustained exposure for two days to different treatments. Arrowheads point to necrotic and disintegrated tissues. Scale bars indicate 0.5 cm.

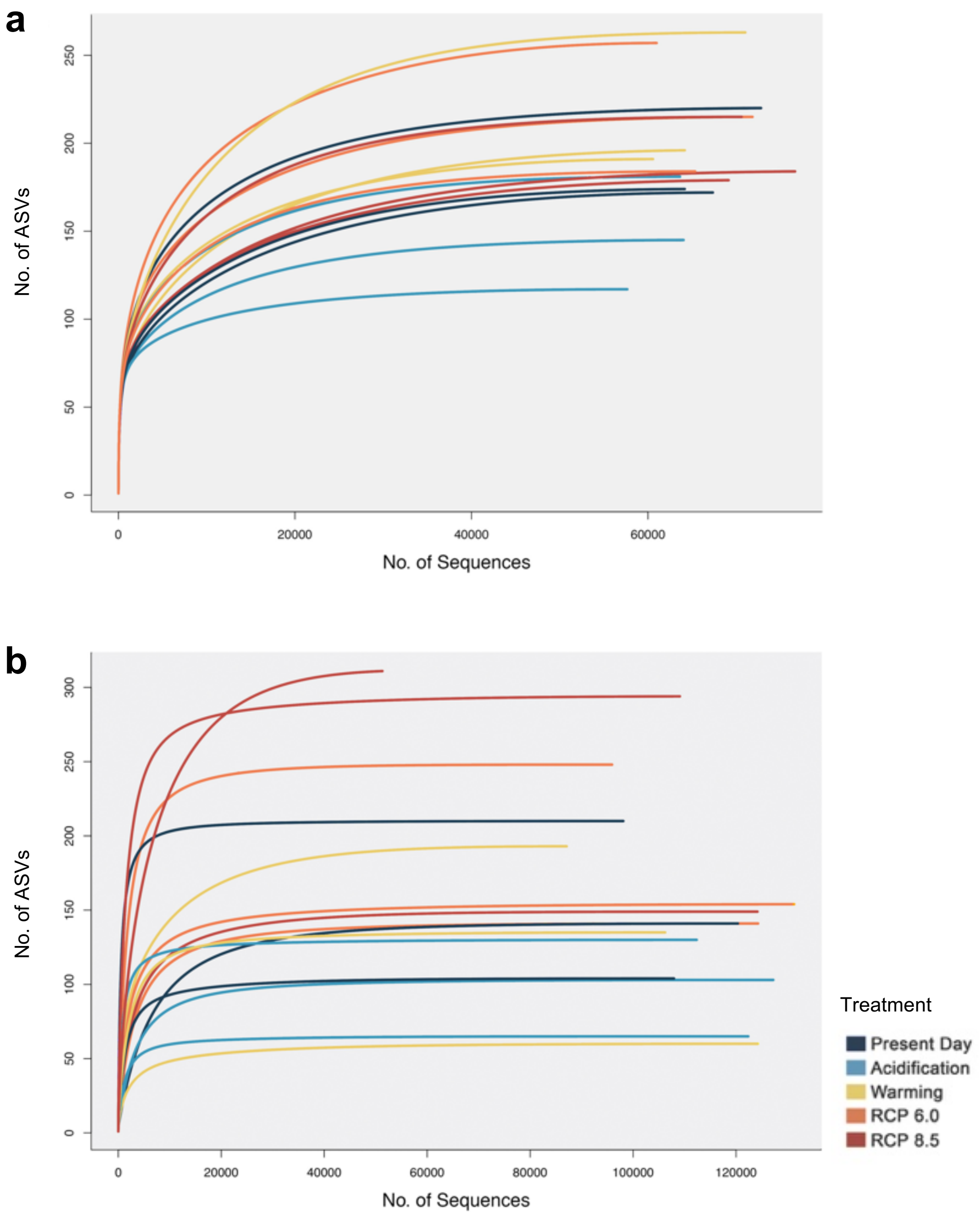


Fig. S5. Sequencing coverage of sponge-associated bacterial communities. Rarefaction curves of ASVs from 16S rRNA gene sequences of (A) *N. compacta* and (B) *L. chagosensis*. Colors represent the different treatments.

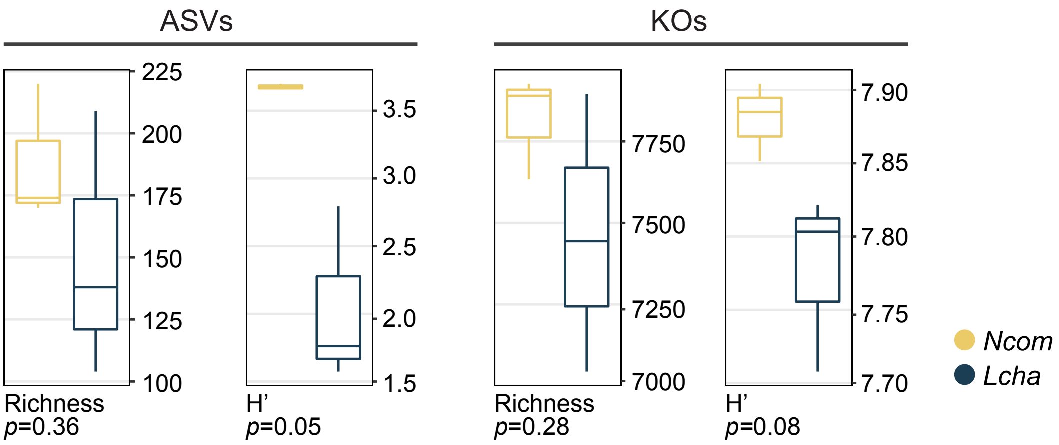


Fig. S6. Richness and Shannon diversity index (H’) of ASVs and KOs of *N. compacta* and *L. chagosensis* microbiomes.

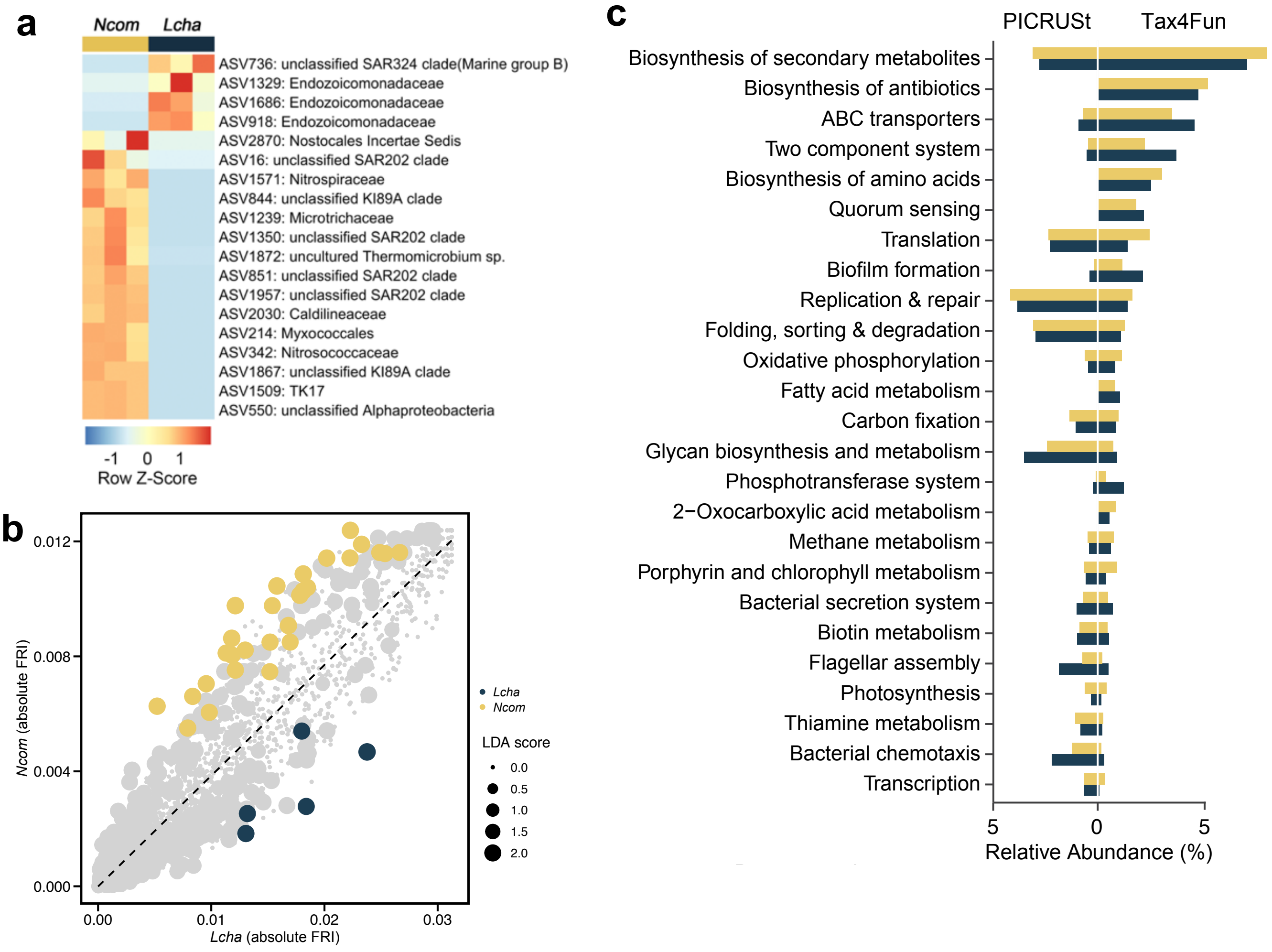


Fig. S7. Distinguishing features of sponge microbiomes. (A) Differentially abundant ASVs between *N. compacta* (yellow) and *L. chagosensis* (blue) presented as relative abundance (row z-scores). (B) Redundant KOs (LDA score ≥ 2.0, *p*<0.05) for each species are indicated by the colored points. The absolute FRI was computed for each KO inferred from the bacterial communities of the two sponge species. The FRI is based on the fraction of all ASVs that are capable of performing a particular KO function and their phylogenetic relationships. A high FRI indicates that a specific function is redundant or ubiquitous in a bacterial community (5). (C) Enriched KEGG pathways predicted through PICRUSt (left) and Tax4Fun (right). ASVs (LDA score ≥ 4.0, *p*<0.05) and predicted functions (LDA score ≥ 2.0, *p*<0.05) that distinguish between the *N. compacta* and *L. chagosensis* microbiomes were determined using Linear Discriminant Analysis effect size (LDA-LEfSe) (6) based on relative abundance values.

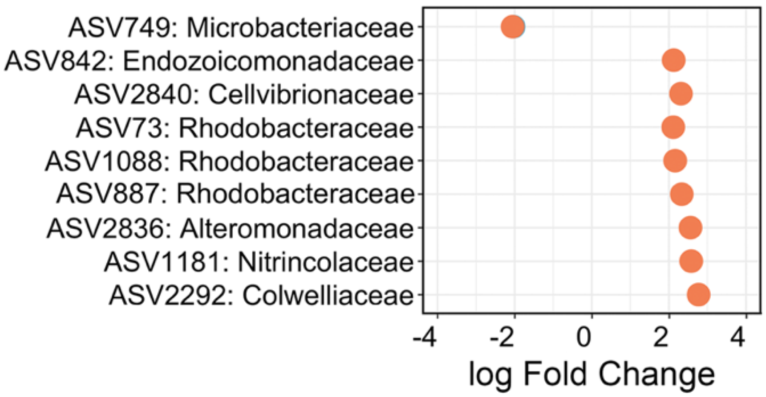


Fig. S8. Differentially abundant ASVs, presented as abundance fold change, in *N. compacta* subjected to Acidification (skyblue) or RCP 6.0 (orange) relative to the Present Day samples.

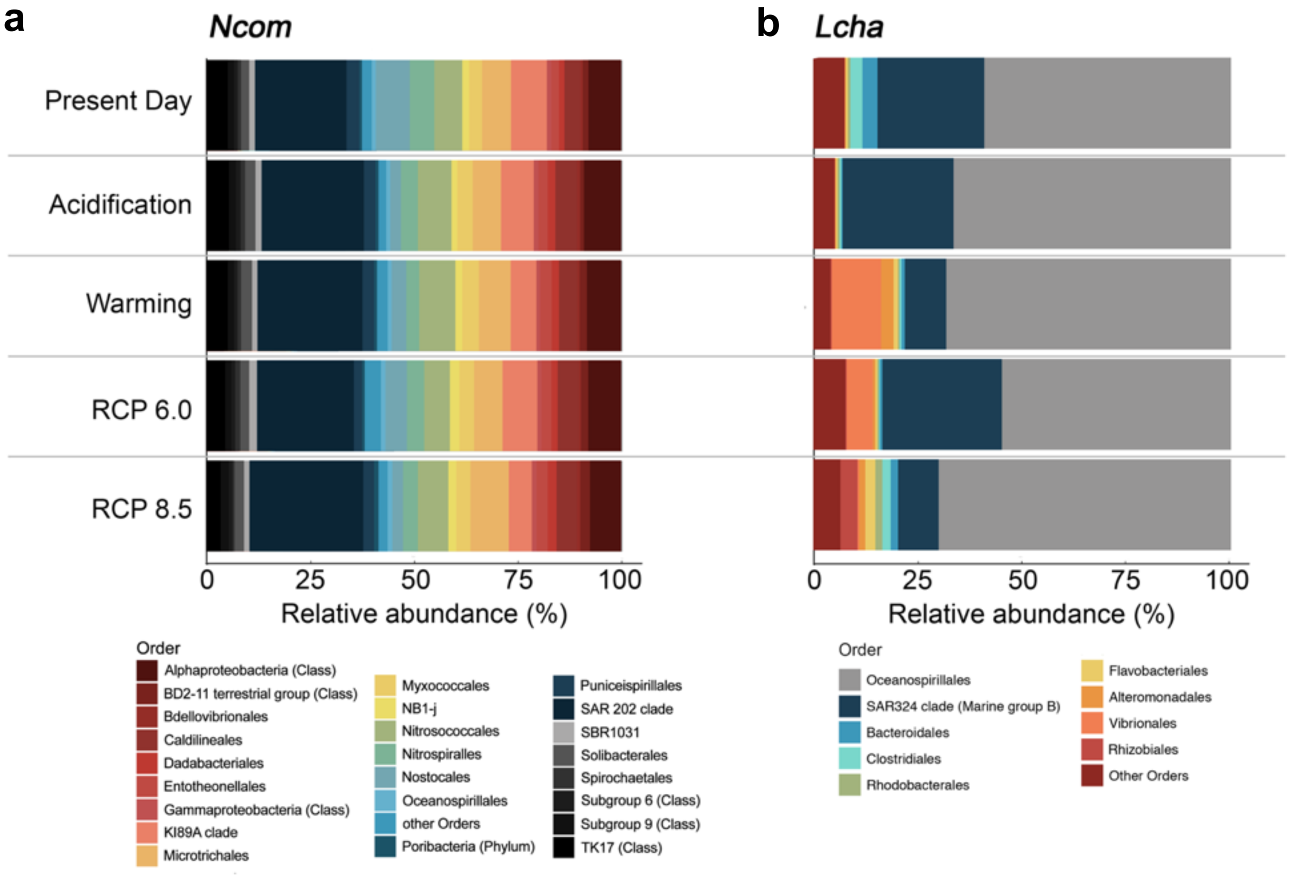


Fig. S9. Taxonomic profiles of the bacterial communities of the two sponge species under different treatment conditions. Relative abundance of major (≥ 1%) bacterial Orders in (A) *N. compacta* and (B) *L. chagosensis* are presented.

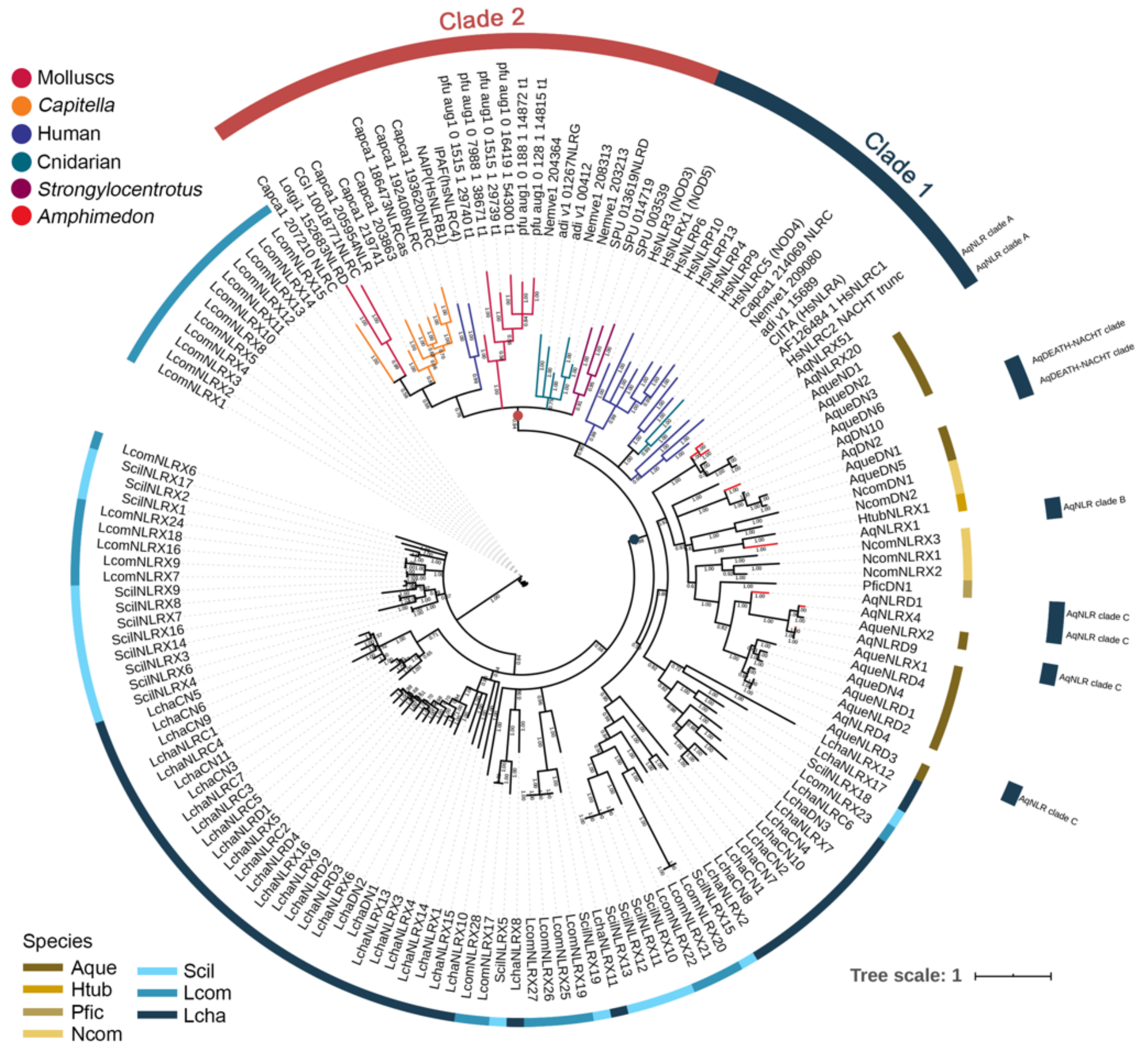


Fig. S10. Phylogenetic analysis of bona fide NLRs of sponges and other metazoans. The phylogenetic tree was derived from Bayesian analysis of aligned NLR sequences from the work of Yuen et al. (7) and from the present study. Numbers on selected branches represent Bayesian posterior probabilities. Outer color strips and annotations indicate the two metazoan NLR clades and the four AqNLR clades based on Yuen et al. (7) Colors of selected branches represent metazoan groups or species. Inner strips indicate the sequences of sponge species in the present study.

Table S1. Carbonate chemistry of seawater in treatment tanks computed using the CO2SYS package (8).

| | | | | | | Forms of Inorganic C (umol/ kg-SW) | | | $CaCO_3$ satn state | |
|---|---|---|---|---|---|---|---|---|---|---|
| Treatment tank | Total Alkalinity (umol/kg-SW) | Dissolved Inorganic Carbon (umol/ kg-SW) | pH | Total Inorganic Carbon | $pCO_2$ | $CO_2$ | $HCO_3^-$ | $CO_3^{--}$ | Omega Ca | Omega Ar |
| PD_A | 2211.55 | 2032.07 | 7.857 | 2016.544 | 646 | 17.93 | 1852.83 | 145.78 | 3.55 | 2.34 |
| PD_B | 2212.17 | 2005.32 | 7.919 | 1993.876 | 547.3 | 15.51 | 1818.47 | 159.89 | 3.89 | 2.56 |
| PD_C | 2213.52 | 2004.63 | 7.918 | 1991.396 | 548.5 | 15.34 | 1813.63 | 162.42 | 3.95 | 2.6 |
| PD_D | 2220.35 | 2033.39 | 7.869 | 2022.643 | 628.1 | 17.65 | 1857.47 | 147.52 | 3.59 | 2.36 |
| Acidification_A | 2229.9 | 2199.08 | 7.465 | 2186.912 | 1777.5 | 50.17 | 2072.51 | 64.23 | 1.57 | 1.03 |
| Acidification_B | 2226.08 | 2198.55 | 7.462 | 2186.335 | 1787.2 | 51 | 2072.76 | 62.57 | 1.53 | 1 |
| Acidification_C | 2219.33 | 2201.36 | 7.433 | 2188.068 | 1914.6 | 54.36 | 2074.69 | 59.02 | 1.44 | 0.95 |
| Acidification_D | 2218.51 | 2198.73 | 7.437 | 2187.169 | 1893.2 | 54.16 | 2074.11 | 58.89 | 1.44 | 0.94 |
| Warming_A | 2217.46 | 2005.27 | 7.934 | 1995.48 | 528.2 | 15.1 | 1817.88 | 162.49 | 3.96 | 2.6 |
| Warming_B | 2226.38 | 2024.58 | 7.913 | 2012.695 | 560.9 | 15.99 | 1839.12 | 157.58 | 3.84 | 2.52 |
| Warming_C | 2225.15 | 2030.58 | 7.894 | 2020.719 | 589.6 | 16.84 | 1852.17 | 151.71 | 3.69 | 2.43 |
| Warming_D | 2224.18 | 2028.44 | 7.902 | 2019.188 | 578.2 | 16.63 | 1850.4 | 152.16 | 3.71 | 2.44 |
| RCP6.0_A | 2224.84 | 2130.51 | 7.671 | 2119.045 | 1053 | 31 | 1994.3 | 93.75 | 2.27 | 1.49 |
| RCP6.0_B | 2218.83 | 2102.82 | 7.728 | 2092.821 | 909.7 | 26.76 | 1961.3 | 104.77 | 2.55 | 1.67 |
| RCP6.0_C | 2222.28 | 2107.12 | 7.729 | 2095.324 | 908.2 | 26.7 | 1963.26 | 105.36 | 2.56 | 1.68 |
| RCP6.0_D | 2212.54 | 2107.75 | 7.705 | 2094.968 | 961.5 | 28.27 | 1966.82 | 99.88 | 2.43 | 1.59 |
| RCP8.5_A | 2236.15 | 2194.49 | 7.505 | 2184.821 | 1607.6 | 46.75 | 2070.58 | 67.49 | 1.64 | 1.08 |
| RCP8.5_B | 2220.92 | 2194.38 | 7.466 | 2182.254 | 1757.7 | 51.11 | 2069.48 | 61.66 | 1.5 | 0.98 |
| RCP8.5_C | 2221.45 | 2186.85 | 7.491 | 2175.916 | 1652.9 | 48.34 | 2063.1 | 64.48 | 1.57 | 1.03 |
| RCP8.5_D | 2228.69 | 2199.69 | 7.477 | 2188.306 | 1716.9 | 50.37 | 2075.51 | 62.43 | 1.52 | 0.99 |

PD=Present Day

Letters (A, B, C, D) in treatment tank codes indicate replicate tanks for each treatment

Table S2. Assembly statistics of *N. compacta* and *L. chagosensis* transcriptome.

| | *N. compacta* | *L. chagosensis* |
|---|---|---|
| No. of transcripts | 69 202 | 91 886 |
| N50 | 1 150 | 1 409 |
| % GC | 46.53 | 45.94 |
| Smallest contig | 300 | 300 |
| Largest contig | 33 061 | 46 605 |
| Alignment rate (%) | 88.77 | 87.29 |
| Mean orf (%) | 69.06 | 59.54 |
| BUSCO Metazoa *odb9* (%) | 89.70 | 90.80 |

Table S3. Comparison of bacterial communities between species and across treatments. Microbiome differentiation was statistically determined through PERMANOVA and ANOSIM tests. Significant comparisons are indicated by bold font.

| Comparison | | PERMANOVA | | ANOSIM | |
|---|---|---|---|---|---|
| Group A | Group B | $R^2$ | Pr(>F) | R | Sig |
| All Ncom | All Lcha | 0.61551 | **0.001** | 0.9913 | **0.001** |
| All Ncom | | 0.14319 | 0.993 | -0.2148 | 0.927 |
| All Lcha | | 0.18269 | 0.954 | -0.1244 | 0.925 |
| Ncom_PD | Lcha_PD | 0.65762 | 0.1 | 1 | 0.1 |
| Ncom_PD | Ncom_Acidification | 0.06671 | 0.7 | -0.3704 | 0.9 |
| Ncom_PD | Ncom_Warming | 0.10787 | 0.7 | -0.1481 | 0.7 |
| Ncom_PD | Ncom_RCP6.0 | 0.06476 | 0.8 | -0.3333 | 0.8 |
| Ncom_PD | Ncom_RCP8.5 | 0.13246 | 0.7 | -0.1481 | 0.6 |
| Lcha_PD | Lcha_Acidification | 0.0696 | 0.7 | -0.1852 | 0.7 |
| Lcha_PD | Lcha_Warming | 0.12891 | 0.8 | -0 | 0.6 |
| Lcha_PD | Lcha_RCP6.0 | 0.03713 | 1 | -0.2963 | 1 |
| Lcha_PD | Lcha_RCP8.5 | 0.16807 | 0.7 | -0.1481 | 1 |

Table S4. Comparison of predicted KEGG level 2 functions of different sponge groups was determined through LDA-LEfSe. Presented are KEGG functions which are significantly enriched in specific sponge group.

| KEGG level 2 function | Sponge group | LDA score | p-value |
|---|---|---|---|
| Metabolism of cofactors and vitamins | Calcarea | 3.8138959 | 0.00517243 |
| Transport and catabolism | Calcarea | 3.60905739 | 2.14E-05 |
| Lipid metabolism | Demospongiae_LMA | 4.25831365 | 0.00028159 |
| Metabolism of terpenoids and polyketides | Demospongiae_LMA | 3.91976679 | 0.00240384 |
| Xenobiotics biodegradation and metabolism | Demospongiae_LMA | 4.10381139 | 0.02009266 |
| Transcription | Demospongiae_HMA | 3.53707926 | 1.49E-05 |
| Cell growth and death | Homoscleromorpha_HMA | 3.87494797 | 0.01116506 |
| Cell motility | Homoscleromorpha_HMA | 4.10800255 | 5.73E-09 |
| Cellular community-prokaryotes | Homoscleromorpha_HMA | 4.11275168 | 0.0005958 |
| Energy metabolism | Homoscleromorpha_HMA | 3.83670935 | 3.67E-06 |
| Folding, sorting, and degradation | Homoscleromorpha_HMA | 3.90671805 | 1.92E-05 |
| Signal transduction | Homoscleromorpha_HMA | 3.7700863 | 2.11E-07 |
| Translation | Homoscleromorpha_HMA | 3.50723491 | 0.01645081 |

Table S5. List of sponge species cited in the present study.

| Species | Taxonomic authority | Reference |
|---|---|---|
| *Neopetrosia compacta* | Ridley & Dendy, 1886 | (9) |
| *Leucetta chagosensis* | Dendy, 1913 | (10) |
| *Rhopaloeides odorabile* | Thompson, Murphy, Bergquist & Evans, 1987 | (11) |
| *Coelocarteria singaporensis* | Carter, 1883 | (12) |
| *Haliclona (Reniera) tubifera* | George & Wilson, 1919 | (13) |
| *Stylissa carteri* | Dendy, 1889 | (14) |
| *Xestospongia testudinaria* | Lamarck, 1815 | (15) |
| *Sycon ciliatum* | Fabricius, 1780 | (16) |
| *Amphimedon queenslandica* | Hooper & van Soest, 2006 | (17) |
| *Petrosia (Petrosia) ficiformis* | Poiret, 1789 | (18) |
| *Leucosolenia complicata* | Montagu, 1814 | (19) |
| *Oscarella carmela* | Muricy & Pearse, 2004 | (20) |
| *Carteriospongia foliascens* | Pallas, 1766 | (21) |
| *Cymbastela coralliophila* | Hooper & Bergquist, 1992 | (22) |
| *Dysidea avara* | Schmidt, 1862 | (23) |
| *Aplysina aerophoba* | Nardo, 1833 | (24) |
| *Neopetrosia problematica* | de Laubenfels, 1930 | (25) |
| *Aphrocallistes vastus* | Schulze, 1886 | (26) |

Data S1. Aligned and trimmed amino acid sequences corresponding to the NACHT domain (PF05729) that were included for phylogenetic comparisons of sponge NACHT domain-containing genes.

>LcomNLRX1
RLVAIVSGAGSGKSTLCL-KILEEFGEKGTHPAC-AQRLKSRFEFPILIRCRELDGGE--
----SLEELLGLQCS--DLDFTDIE---RKSVIKHL-RHH--ANKVLLVVDGLDEAIPEA
LGSNSELRKLLDG---------AKSLVGASVILTG-RLCDAIKDVVISR-TRAVCYALAG
F-SGQAFKDYGD-----DEFRAA

>LcomNLRX2
RLVAIVSGAGSGKSTLCL-KILEEFGEKGTHPAC-AQRLKSRFEFPILIRCRELDGGE--
----SLEELLGLQCS--DLDFTDIE---RKSVIKHL-RHH--ANKVLLVVDGLDEAIPEA
LGSNSELRKLLDG---------AKSLVGASVILTG-RLCDAIKDVVISR-TRAVCYALAG
F-SGQAFKDYGD-----DEFRAA

>LcomNLRX3
RLVAIVSGAGSGKSTLCL-KILEEFGEKGTHPAC-AQRLKSRFEFPILIRCRELDGGE--
----SLEELLGLQCS--DLDFTDIE---RKSVIKHL-RHH--ANKVLLVVDGLDEAIPEA
LGSNSELRKLLDG---------AKSLVGASVILTG-RLCDAIKDVVISR-TRAVCYALAG
F-SGQAFKDYGD-----DEFRAA

>LcomNLRX4
RLVAIVSGAGSGKSTLCL-KILEEFGEKGTHPAC-AQRLKSRFEFPILIRCRELDGGE--
----SLEELLGLQCS--DLDFTDIE---RKSVIKHL-RHH--ANKVLLVVDGLDEAIPEA
LGSNSELRKLLDG---------AKSLVGASVILTG-RLCDAIKDVVISR-TRAVCYALAG
F-SGQAFKDYGD-----DEFRAA

>LcomNLRX5
RLVAIVSGAGSGKSTLCL-KILEEFGEKGTHPAC-AQRLKSRFEFPILIRCRELDGGE--
----SLEELLGLQCS--DLDFTDIE---RKSVIKHL-RHH--ANKVLLVVDGLDEAIPEA
LGSNSELRKLLDG---------AKSLVGASVILTG-RLCDAIKDVVISR-TRAVCYALAG
F-SGQAFKDYGD-----DEFRAA

>LcomNLRX8
RLVAIVSGAGSGKSTLCL-KILEEFGEKGTHPAC-AQRLKSRFEFPILIRCRELDGGE--
----SLEELLGLQCS--DLDFTDIE---RKSVIKHL-RHH--ANKVLLVVDGLDEAIPEA
LGSNSELRKLLDG---------AKSLVGASVILTG-RLCDAIKDVVISR-TRAVCYALAG
F-SGQAFKDYGD-----DEFRAA

>LcomNLRX10
RLVAIVSGAGSGKSTLCL-KILEEFGEKGTHPAC-AQRLKSRFEFPILIRCRELDGGE--
----SLEELLGLQCS--DLDFTDIE---RKSVIKHL-RHH--ANKVLLVVDGLDEAIPEA
LGSNSELRKLLDG---------AKSLVGASVILTG-RLCDAIKDVVISR-TRAVCYALAG
F-SGQAFKDYGD-----DEFRAA

>LcomNLRX11
RLVAIVSGAGSGKSTLCL-KILEEFGEKGTHPAC-AQRLKSRFEFPILIRCRELDGGE--
----SLEELLGLQCS--DLDFTDIE---RKSVIKHL-RHH--ANKVLLVVDGLDEAIPEA
LGSNSELRKLLDG---------AKSLVGASVILTG-RLCDAIKDVVISR-TRAVCYALAG
F-SGQAFKDYGD-----DEFRAA

>LcomNLRX12
RLVAIVSGAGSGKSTLCL-KILEEFGEKGTHPAC-AQRLKSRFEFPILIRCRELDGGE--
----SLEELLGLQCS--DLDFTDIE---RKSVIKHL-RHH--ANKVLLVVDGLDEAIPEA
LGSNSELRKLLDG---------AKSLVGASVILTG-RLCDAIKDVVISR-TRAVCYALAG
F-SGQAFKDYGD-----DEFRAA

>LcomNLRX13
RLVAIVSGAGSGKSTLCL-KILEEFGEKGTHPAC-AQRLKSRFEFPILIRCRELDGGE--
----SLEELLGLQCS--DLDFTDIE---RKSVIKHL-RHH--ANKVLLVVDGLDEAIPEA
LGSNSELRKLLDG---------AKSLVGASVILTG-RLCDAIKDVVISR-TRAVCYALAG
F-SGQAFKDYGD-----DEFRAA

>LcomNLRX14
RLVAIVSGAGSGKSTLCL-KILEEFGEKGTHPAC-AQRLKSRFEFPILIRCRELDGGE--
----SLEELLGLQCS--DLDFTDIE---RKSVIKHL-RHH--ANKVLLVVDGLDEAIPEA
LGSNSELRKLLDG---------AKSLVGASVILTG-RLCDAIKDVVISR-TRAVCYALAG
F-SGQAFKDYGD-----DEFRAA

>LcomNLRX15
RLVAIVSGAGSGKSTLCL-KILEEFGEKGTHPAC-AQRLKSRFEFPILIRCRELDGGE--
----SLEELLGLQCS--DLDFTDIE---RKSVIKHL-RHH--ANKVLLVVDGLDEAIPEA
LGSNSELRKLLDG---------AKSLVGASVILTG-RLCDAIKDVVISR-TRAVCYALAG
F-SGQAFKDYGD-----DEFRAA

>LchaNLRX17
RHIAIVGSSGAGKSVACQ-LLCSNVSAG-------V--LKERLSRVLCWDFRDVQVRK--
--ATGITELLQIATQ----CSDATT---CRPLAEELLASR--GKGVLLIFDGLEQFSAA-
---DSVIWSLI----------NGYLLSLSHVVVTT-RPCGIVKRLTNF---VFYQLQLDE
V-LE-------------------

>LchaNLRX10
----------------------SAYLKRK------A--ELSRYDFVFVVPARNLTEAN--
--GDNVFSLL-NLHH----YLSAEE---VADVLPHLK-DN--ADRVLILLDGADEMGSS-
LSKSKAVEDLL----------DGPSLSGRTLLVTS-RVSDLADRFMKE---ACSCAMILE
L-TDEQLDSMVRLRLNAK-----

>ScilNLRX18
-RIGLHGAPSLGKSQVCQ-RLLLNWAYR-------R--QLARFYLVVHWELRDASVQS--
--ASSLQDLLLALGV----QDDM-----VSSAVPALQRMR--GRGCLFILDGLDEMSPQ-

TAAGQFVRQLM----------DGRALPEACLLVTS-RPCAQSEALFAK---YTIQLDILG
F-SDQQAEEFISCHLAT------

>LcomNLRX23
------------------------------------------------------------
-------------------------------------RER--GRGCLFVLDGLEELQAG-
NTTSTYIRRLL----------RGKVLPDACLLVTS-RACSEADQLFTT---YSSTAEILG
F-SDSQAESFIRLQIPE------

>LchaDN3
KHVGIIGGTGCGKTNLCT-SITTFYS---------K--CWPMFSLVLFWRLYDPSVQK--
--AENLKELLCAVGP----SWTTIR---ALRLAETLLATA--GRGVLVILDGVDQLEAD-
--ENACIWRLL----------EGSVLQEARLLVTS-RPCFLAKHYFDT---YDINLELLG
F-TEEQVSHFIYHCLGED-----

>LchaNLRX7
KRVGMYGSSGCGKTFCCT-ALTQLYASQ-------Q--LWKHFRLVLFWRLSDPIVQE--
--AKTLVQLLFALLP----SASLRR---RQRLASVLNTSN--GKGILIILDGVDQLEGG-
--ERAFVRHLL----------SGEAFKEACLLVTS-RPCSLAENLFSG---YNVQLVLKG
Y-TRKQVVSMLDRRLG-------

>LchaNLRC6
QQIGLYGGAGCGKTASCL-KALKERSEG-------R--LWNGFQLVLLWHLRKPDVQR--
--ASTLKDLLRALPV----PLSERQ---STRLSEHLEESD--GRGVLLVLDGIDELRRP-
S-NNAYVVRLL----------ERTRLHRSSILVTS-RPCYEAEQYFKT---YDVCYEIIG
F-TDDQVLSYIHYYLEEE-----

>LchaCN4
KRVSIYGGAGCGKTSCLL-KIASMYAEK-------R--LWPEAHAFLLWKLRDREVQK--
--AQNLEQLLRLLKL----SLTEEK---CKELAAVLEASE--GEGVILALDGIDELDTQ-
--KNGIVWSLL----------DGSVLSEACVLATS-RPSSIAKTFFNE---YDVNLELLG
F-TEEQVDQFVEQQFVDK-----

>LchaCN10
KRVGIYGGAGCGKTSSLM-KAASMYANG-------Q--LWQGARALLLWKLRNPDVQT--
--ADCLEQLLLQLMI----GLTEQQ---CKHFTEVLLSCA--GEGVILALDGVDELDAQ-
--QEGYIWSLL----------DGSALSKACVLATS-RSCSTAKTYFDK---YEVNLELLG
F-TEEQTGQFIQQQLGDQ-----

>LchaCN2
QRVGIYGGAGCGKTSCLT-KAASLYAEK-------Q--LWQGARTFLLWKLRDPNVQE--
--ASSLTQLLRQLSL----DLTEKK---CEELASVLSSSD--GEDVILALDGIDELDTK-
--VKGYVRRLL----------DGSALKKAYVLATS-RPCSTARTYFDR---YGVNLELLG
F-TEEQVDQFIQQQLGDD-----

>AqueDN1
RFVLIEGEPGIGKSTLAK-ELVLRWANR-------SDKLLSNYDIVFFIQLRFETYHK--
--ATSIEDLFVDLDN----QT--IN---MTDLNIEIKKRK--GAGILWILDGFDELPSH-
LR-NSILMTLI----------KGVILPKSTVIVTS-RPVASDLLLELLKDDNSKRISLRG
F-DSTKIGEFALKYFNDK-----

>AqueDN5
RFVLIEGEPGIGKSTLAK-ELVLRWAKG-------SDELMNNYDIVFLIQLRFETYHK--
--ATSIEHLFVDLDD----QS--IN---MTELHVEIKKRK--GAGILWILDGFDELPSH-
LK-NSVLMKLI----------KGDNLSKSTVIVTS-RPVASDQLLHFLHEHDSKRISLRG
F-DSTKIEEYALQYFNDK-----

>AqueND1
LRVVIDGPPGIGKTTLCR-KLLNMWSNG-------TLV-HQQYDLVLYCPLRNSKIAT--
--ATTLADLF--EY-----RC--CE---VPIVAKWFEKRN--GEGLLIIFDGWDELSEQ-
LR-SSLAASII----------HREQLDQCSVIVTS-RSYASSSLLKMDT--LSRHVQVIG
F-SKKEISKVIIRTLQKD-----

>AqueDN2
LRVAIDGPPGIGKTTLCR-KLLNMWSKG-------SHE-LQHYDLVLYCPFRHKAIAE--
--ATKLVDLF--VY-----ES--PK---VSKVVDWILERE--GKGLLIIFDGWDELSTQ-
LR-SSLATKII----------CRNQLVRSSVIVTS-RTYATASIYQLEC--LNLNVHVIG
F-AADEINYVIKGMLSQK-----

>AqueDN3
LRVAIDGPPGIGKTTLCR-KLLNMWSKG-------SHE-FQHYDIVLYCPFRHKAVAE--
--ATELADLLTCAY-----KS--PK---VSEVADWILERE--GKGLLIIFDGWDELSTQ-
LQ-SSLATSII----------CRDQLVYSSVIVTS-RTYATASLLQLEC--LNQNVHVIG
F-VAAEIDEVIKGTLSSP-----

>NcomDN1
-KILLVGVAGSGKTTISW-HACQQWAEG-------N--MFQEFEFLIYLSLADPHIQS--
--ASSFKDLIGHPCN----EV--C-----NVVAKAIEGPK--KKKVCFVMDGWENLPTTL
QE-STFLRSLIQG------NASEV------VIVTS-RPIAAGSIVFSM----FFTYNIRE
F-SDDDIAFCAYQYLPTK-----

>NcomDN2
TKVLLEGVAGSGKTTVTW-YACRGWAEE-------E--MFHEFDYLIHLTLADPELHS--
--AKTIEDIIPHPSS----EM--R-----KAVADAIIKLN--GRGCCFILDGWEDLPSQA
RISNSILACLLHG------NNVRMALPQCSFLVTS-RPAVSDSLINSV----SRVITMEG
F-PEDAIDSYANRYFCTQ-----

>HtubNLRX1
-STLISGRPGIGKTVLLT-KVCKDWGKG-------N--CLKNIQVIIHVSLRKLHCKS--
-PQPSLLDMIELHFT----DS--DR---SKQFCRLIEAKG--GKNVCFAFDGLDEYPLL-
NT-NDIVSEII----------GKQKLPMASVIVTS-RPTASHIVKQKM----KKHAEIIG

F-MPHQIESYINEYYTSS-----

>AqueDN6
KVIIIEGCPGIGKTTLAS-KLCQEWSEK-------R--QMLEFKLLLYIPLRSPLMRT--
AQ--SIDELLEYYG-----DN--YT----SNDVLLIKKNQ--GRDVVFILDGWDELRPS-
CRIDMFFPGLI----------YGKFLPESTIVVTS-RPGATIDIRRHA----NRIVEILG
F-TEDQIKQYIMSYFKTN-----

>NcomNLRX3
KMVLIQGAPGVGKTLLAR-CICQRWAKG-------E--ILTEYQLVLFVPLRAFPANS--
-GSLSLHNIVELYLC----GS--GI----DDAVRELSEHN--GKKVLLILEGWDELPPE-
LREFTIFNDLV----------MGTKLPEASIMVTS-RPIVVDELYKTI---KDRQIEVLG
F-KEKQIKEYLKHNLKAE-----

>NcomNLRX1
SAVLIEGEQGKGKSMFSL-YLCKKWAKG-------D--MLNQFNIVLLVPLHRFSPES--
ALQLTTRELIEIYLP----GS--AG----SKASNDIEFSD--GEKVLFILDGWDELTPV-
LQ-NGFFRDLL----------LGKILTKASVLVTS-RPTAALQLGQLV----SRAYQLLG
F-SQYEVQRFLRLHVPD------

>NcomNLRX2
HKVLVEGAPGIGKTMLAQ-YLSREWAEG-------R--LLKEFNLVLLVPLRRFSSKS--
ADSLTIQDLVQIYLS----GD--LG----KKASKQLELSG--GEKLLIILEGWDELPSE-
LRDMTLFHDLL----------IGHKLPKASVLVTS-RPTVAGDLYQFV----NRRIEVLG
F-LEEQIEEYVKFHTRDK-----

>PficDN1
TRILVKGAPGIGKSSLAI-ELCKKWSIG-------E--LGSHFKLVILLRLKVPKIQK--
--AKKIEDLI---MP----ER--Y-----RNCIESLGDCL--GEGVLFILEGYDELPDEQ
RELQTLME--E----------IEEELPKASVMVTS-RPWATLG--IGEF--FQEHVQILG
F-TESSRREYVDSVLSSD-----

>AqueNLRX1
SWILIEGVSGIGKSTLAY-EMLKQWKDG-------T--ALQNYSYVLLLRFRNENVHQYW
SPSK-VTQLIQECLN----EQ--YH---EQPDI---LSNS--GQDLLLILEGYDELPKKE
LEHAQVFY--Q----------LNRNFHNAVVVITT-HPSFSYQLSDKIL--FAKKIEIIG
F-DKPNQDQYIKVAFKND-----

>AqueNLRX2
SWILIEGVSGIGKSTLAY-EMLKQWKDG-------T--ALQQYSYVLLLRLRNENVHQQW
SPSPDVPQLIEDCLN----EQ--YI---EPPEIKSILDNK--GHNLLLILEGYDELPKGK
FKQMEVFR--Q----------LKSDFHNAVVAITI-HPSFLYQLSENIL--FTKEIEILG
F-NKSSQNRYIEIAFKND-----

>AqueNLRD4
DAILIEGAPGIGKSMLAF-EICSRWVKG-------K--ALMGYTLLLLFRLREKFVQD--

--CETVKELLGCFLV----GQ--SW---KEEVVRDIIDNS--GEGVIIILEGFDELPHE-
LTPDSVFL--K----------LSFELPSASFIFTS-RPSAKHCLKQEIL--FERHVEVIG
F-TKPSIEKYVTEFFKGN-----

>AqueDN4
EAVLIEGAPGIGKSMLAF-EICSRWVKG-------E--ALKKYPLLLLLRLRDRVIKN--
--CESVRELLGCFLK----EQ--SW---KDAVVQDIIDNG--GEGLIVILEGFDELPEH-
LTQDSVFF--Q----------FSTVLPCASLVFTS-RPSAKHFLRLKVE--FELHVEVIG
F-THENINEYIQKFCKGN-----

>AqueNLRD1
DAILIEGAPGIGKSMLAF-EMCSRWVKG-------E--ALKKYPLLLLLRLRDRVIQN--
--CKSVKDLLGCFLK----EQ--SW---KDAAVQDIIDNG--GEGLIVILEGFDELPEH-
LTRGSLFF--Q----------FSNELPCATLVFTS-RPSAKHYLKNEIE--FERHVEVIG
F-KKENIDEYIEKFCNNN-----

>AqueNLRD2
EAVLIEGAPGIGKSMLAF-EICSRWIKG-------E--ALKKYPLLLLLRLRDKFIQN--
--CKTVKDLLGCFLK----EQ--SW---KEEAVQHIFDKS--GEGLIVILEGFDELPED-
LTASSVFL--Q----------ISEELPFTSLIYTS-RPSAKHSLKQEIS--FSRHIEVLG
F-TGKSINEYIQLFFEND-----

>AqueNLRD3
EAVLIEGAPGIGKSMLAF-EICSRWVKG-------E--ALKKYPLLLLLRLRDRVIQN--
--CKSVRELLGCFLT----EQ--NW---KDATVQHIFNKS--GKGLIVILEGFDELPED-
LTPGSVFL--N----------ISEELPFASLIYTS-RPSAKHYLRQEIT--FSHHIEVIG
F-TSKSISQYIQIFFQDN-----

>LcomNLRX20
RNILAAGPPGSGKSYLFTKIIPFLWSC--------DLLWKGRFDLVAFFDLCREDVRN--
--AADMKELLATFTE--DTMMDDGE--RSAAYFHSA-RMM--AERLCLIFDGLSECSVA-
--CSEFIRGIL----------GRTRWSACHVIVLT-RHLTDATIVDKRF-QYHRFLEVSG
L-TRCAMDGIITSRVSAD-----

>LcomNLRX21
RNILAAGPPGSGKSYLFTKIIPFLWSC--------DLLWKGRFDLVAFFDLCREDVRN--
--AADMKELLATFTE--DTMMDDGE--RSAAYFHSA-RMM--AERLCLIFDGLSECSVA-
--CSEFIRGIL----------GRTRWSACHVIVLT-RHLTDATIVDKRF-QYHRFLEVSG
L-TRCAMDGIITSRVSAD-----

>LchaCN1
RRQIATGCAGAGKTTAFTLKAPYEWAKEG------SDFWQQFRLF-FFGSLNDTKWRD--
--SKSLNDVFGLGEF----GLTKRA---QTDVLTFI-RNH--SEKVLLVADSFEEAPQL-
--HSSLLWMVLSG--------KEL--PQLNVMVSS-RPCKMASWLSRNF-PFHQRLEVMG
F-TLEKILLYVKEYFSHD-----

>LchaCN8
RRLIATGCSGAGKTTAFTIKAPYEWAKEG------SDFWQQFKLF-FYGSFNDTKWRD--
--AESLADVFHLNEF----QLTEAE---QKEVLAYI-REH--PDKVLLVADSLEEAPEP-
--QSSMLLKVLSG--------KERGLLNLNVVVSS-RPCRITAVLSKDC-PFDQRVEVAG
F-TPDKIKRYVEWFFSQD-----

>SciNLRX10
---IAVASAGCGKTFAFTKVAPLKWAM--------GKLCQRKKLL-IARELHHEDVKM--
--AKSLSGLLGLEGI----GIEDSH--DRQVICEYV-RAQ--PDALCLVLDGLDEINLS-
--CSSFVQGVI----------QGEELPGVHLIVTS-RPCPDVFSLSAMP-HFQQHVELVG
F-QPDDVQMYVNKVLRSE-----

>SciNLRX11
---IAVASAGCGKTFAFTKVAPLKWAM--------GKLCQRKKLL-IARELHHEDVKM--
--AKSLSGLLGLEGI----GIEDSH--DRQVICEYV-RAQ--PDALCLVLDGLDEINLS-
--CSSFVQGVI----------QGEELPGVHLIVTS-RPCPDVFSLSAMP-HFQQHVELVG
F-QPDDVQMYVNKVLRSE-----

>SciNLRX12
---IAVASAGCGKTFAFTKVAPLKWAM--------GKLCQRKKLL-IARELHHEDVKM--
--AKSLSGLLGLEGI----GIEDSH--DRQVICEYV-RAQ--PDALCLVLDGLDEINLS-
--CSSFVQGVI----------QGEELPGVHLIVTS-RPCPDVFSLSAMP-HFQQHVELVG
F-QPDDVQMYVNKVLRSE-----

>SciNLRX13
---IAVASAGCGKTFAFTKVAPLKWAM--------GKLCQRKKLL-IARELHHEDVKM--
--AKSLSGLLGLEGI----GIEDSH--DRQVICEYV-RAQ--PDALCLVLDGLDEINLS-
--CSSFVQGVI----------QGEELPGVHLIVTS-RPCPDVFSLSAMP-HFQQHVELVG
F-QPDDVQMYVNKVLRSE-----

>LcomNLRX22
--------AGCRETELFLTKSPRDWAI--------GKLWREFDLL-VARELRSDSVHK--
--ARNVSDLFALEYY----GVRSLE--EQQIVSKFV-QRN--PARVCLILVGLDEIQMS-
--CSSFMQQVI----------NGETLRGIRLLLTS-KPSPEVFKLSVSS-PFDRHIEVAG
F-LPQNTRNYICKALSPN-----

>SciNLRX15
RRVLAVASAGCGKTVLFTIKIPHDWAS--------GDLWAAHFDLLCVVQLNDVAARS--
--AHNTEELLQLATL----GLTAAE---QAELAQYV-HAH--PERVCLILDGLDECHVY-
--CSLFVQQVIYD--------QCPCLAGMRVIITS-RPCIATSVLTQRA-RVDRRLEIIG
F-THSSVYDFVRKYLTGE-----

>LchaNLRX2
TRVLAIGTAGSGKSTAFTVKAPHDWSL--------GQLWPETVLFQCL-KLRDRSVWK--
--AQSVSELFQLNAL----GLNDAE---RTEVENFI-VKN--PGRVLLACDGLDECVVQ-
--FGLLWSV---L--------QGKALAGVRLLMTS-RPCELLIDLSRET-AIDSHVRLFG

F-TEENVLVFVDRYVGGA-----

>LchaNLRX15
-------------SRFCF-RVLLQWACYK---------VFRQFAIVFYISMRDHERSC--
--TTSVASLLRLDVM----GFKSPE---QRSILAYL-NKE--SHKVLLIVDSC---THDI
SQQGSAVQQLFDG----------VLFPNASIIISA-QPSLSLLQLTARC-E--RHYCFQ-
-----------------------

>LchaNLRX1
-RCLLLSPPGGGKTMTCH-QILNLWAGGK---------SFKHFKAVVYLTGKEDQRVC--
--TSKVGEFLALNDS----EPTDRKSLTERLIQSHV-ASH--GEELLIILDAADESSSDS
LFDGGILATLFSPAKSLS--SGLGKLTDASVIVTS-RPCPASDYLVQGG-DCTAVLWLCG
F-TDYNLQQLLRKRLGEE-----

>LchaNLRX14
KRSVIFGGGGSGKTLLCR-KILSDWDNIR------LVNSFKKFWAVMYISGRNTGRIH--
--ATNASTFLGINDR----NLSDRQ---CCEIVGFM-AEN--SERLLVLIDGWDEVSGSG
LLGGTVLTDLLRR---------RGCFSQASILITS-RPCPEVFQVLELC-KISRFFSLVG
L-KEKRIRELACRKLGDH-----

>LchaNLRX4
QRALVIGPGGSGKSLTCS-KIIADWIDRT---------AFRQFLAVVHISAKDTDHWQ--
--ARSSSDLLCLDSH----GYSQSQ---QTRLLLQL-RKR--SEKLLFLIDGSDEAAGEG
LLPGTALHDVLNR----------KLFRNASVIITT-RPSPSCYNLLNIC-D--KCFYLAG
F-TEKHLNEFVRNRLEP------

>LchaNLRX13
KSALILGMSGSGKSLTVL-KILSEWIGEA-------KSAVKQFGMVVLVTGRETTRLN--
--AKAVEDFFQLWRL----GYNERQ---QRYILDYF-KESEDAKSVLILIDGWNEGGEAC
LLDGTVLKDMLRGER-------SKLFPGCSVIITS-RPTESIYPLLEEC-D--ARYNLLG
L-SNQQLRQLLSRRLNEK-----

>LchaNLRX3
-RSLVLGGGGSGKSLTCY-KILAEWSGDD-------PSPFKEFDAVFYITGREQKRLS--
--AKSRAQLLRFNSM----GFDKER---ERQLVEYF-TEH--SEKMLFLIDGGDELSNGG
LLQGLAMRRVLRG----------QLFPKACVIITS-RPCPGAYKLLNVC-Q--RHFALVG
F-TDDNLHHLIKCRLGQQ-----

>LchaDN1
-SALLFGEAGSGKSLTTL-KILSEWNKNE----LQSTLPFKKFDMIVYVTGRESIRLQ--
--AKEIQNFWQLWRF----GYSGGQ---QRHILHHY-EHH--SEKVLFVIDAWDECGNDS
PLEDTVIMQILHG----------KLFRNSSVIITS-RPSPSAYPLLKVC-K--RRYSLVG
F-KNKRLRELFDRRLGEQ-----

>LchaDN2
KSALLFGEAGSGKSLSAL-KVLSEWSKKE----LLSTSPFKQFDVVVYVTGRESTRMQ--

--AKDVQNFWLLWQC----GYSEDQ---QEHILHHY-ERH--SEKVLFLIDAWDECRSDG
LLKDTVMMKILHK----------KLFCNSSVIITS-RPSPNA------------------
----------------------

>LchaNLRX6
KSAIIFGSAGSGKSLTVL-KILSEWWSQN------KTSPFERFDVVIHITGRETSRIR--
--TKSIQELWQLWRM----GYTDHQ---QQFIVKHY-SQH--SEKVLFLVDAWDEAGDNG
LLEDTALKRILHG----------ELFQYSPVIITS-RPAPNTYLLLHKC-K--RRYSLVG
F-NNDRLKELFDRRLEEP-----

>LchaNLRD4
RSTILFGSAGSGKSLTTL-KILSEWSSRN------SGSPFKKFDVIVHLTGKERSRLR--
--TTDVHELWRLQRM----DFSDEQ---QLIIIDYF-AHH--SEKVLFLIDGWDEAGDDG
ILEGSVLKAILDG----------NLFPQSSTIITS-RPVPTAYPLLEKC-R--YCCLLVG
F-SDKRLQEFFHQRLGEQ-----

>LchaNLRD2
QRTLLFGSAGSGKSLTML-KILSEWIAKD------SASPFKRFDAIVYITGRERSRLS--
--TMDVNDLWRIKYL----GYTEQQ---EEFLLDHF-SRH--SDKVLFLIDGWDECWDDN
LLEDTVMKQILCG----------DLFPQSAVIITS-RPAPNAYPLLEKC-K--HRYALVG
F-SDRRLQELFNRRLGEQ-----

>LchaNLRX16
--------------------------------------------------RERLRLI--
--TKEVHDLWQMKQM----GFNERQ---GLFILDHF-SHH--SDRVLFLLDGWDEAGDSG
VLEDSALKQILDG----------DLFPQSAIIITS-RPAPNAYPLLEKC-K--QRYTLVG
F-NDKRLRELFNRRLGET-----

>LchaNLRX5
KSALLFGSAGSGKTLTTL-KILSEWNSKE------TTSPFQQFDVIVYVTGKERARLR--
--AKDIHDFWRLEQM----GFNEKQ---QQFILNHY-SSL--SEKVLFLIDGWDEGGDDD
LLGDSALRKILQR----------DTFPQSAVIITT-RPAPNVHPLLETC-R--HRYSLVG
F-NDSRLQELFHQRLEES-----

>LchaNLRX9
-RALIFGSAGSGKSLTTL-KILSEWTSQS------CASLFKRFDAVVYITGRERSRLT--
--TDNVSNLWQLQHN----GFNKQQ---ECFLLNHF-SEC--SDKVLFLVDGWEEAGEHG
LLEDMVLNKILNG----------ELFPQSAVIITS-RPAPTAYHLLEKC-K--NRYSLTG
F-NDRRLQELFNRRLSES-----

>LchaNLRD3
-SALLFGSAGSGKSLTTL-KILSEWMDSD------TKSAFNKFDVIIYIAGRERSRIS--
--TSDVRELWQLEQM----GFNEQQ---QLFFLDHF-LQN--SSKVLYIIDGWDEAGDDG
LLENTVLKRILHR----------ELFPESSIIITS-RPAPNAFPLLEKC-L--RRYSLVG
F-NDRRLRELFCYRLGDS-----

>LchaNLRC2
-SALLFGLAGSGKSLTTL-KILSEWTSKT------SLSMFKQFDAIIYATGKERSRLR--
--AMDVAEFWQLQQM----GFNEQQ---SHFILGYF-LQN--SERVLFLIDGWDEAGDDG
LLEGSVLKNIVDG----------ELFPQSSIVITS-RPAPNAYPLLEKC-Y--HRYSLVG
F-NDKRLKELFNRHLNET-----

>LchaNLRD1
-SALLFGSAGSGKSLTTL-KILSEWNSES------STSAFKQFDAVVYLTGKERSRLG--
--AANVQDLWRLQQM----GFSEQE---ELFILNHF-SQH--SEKVLFLVDGWDEAGDHG
ILDDSVLKKILDG----------DLFPQSAIIITT-RPVPNAYPLLEKC-K--QRYSLVG
F-NNTHLQELFHRQMEEA-----

>LchaNLRX12
HVVAITGSAGSGKTTVLR-QLAAAWAKLKCG--PAHLHWVSRYRFVLYLDVSVTDSGN--
--ISLRSQLLPQN--------SLLR---VGLLLKEISENG--GKEVLVLLDGAHIRIHPI
L-SSSEIGQLLQC----------QLLPLSTVVVTHNDDCPLPTIYRRDGRL--LELEVAE
L-TLREREDLVKLYVGDN-----

>SciINLRX19
RQYLVYGPPGVGK-----------------------------------------------
------------------------------------------------------------
--TVALSYKLPFEWAT----EKALKIYKFVIVVPL-RESHLAAELDPAI-L--LLSRISR
LREDAELRHKIYKVLEREEKQLL

Data S2. Aligned and trimmed 28S rRNA gene sequences of *N. compacta* and *L. chagosensis* and of representative sponge species retrieved from NCBI.

>Ncom03
--------------------TTTCCGCCGGGGCACGTC---------
ATCGGCGATGGCTCTCATCCCTCGAGGCG-----------GCAAAGGGTTGCC-------
TCCTCCGA-------GGTCAAGTGGCCGGGA---------------------------------------------------
CGGCCTTGCGGTCTTCTCGGAACGC----------A-TACTTT--GCCGTTGGGG--
GCGTGCGAGCG------------------------------------------
AGCAACGAGTGAGTTCGTTCCGTC-----GGCGTCCAT---TCACCTG---
GAGCAAAGGCAGGTGCGGGCCAGAGAGTTTGGCCGGGCGGCCCACG-
CGTCCGGTAGTCTCGAGC--------------GAGTGCTGCC-----------
CCGTCCTCGCGGCGGGCGGCTCACTCGAGTTCTG---CCGGCGTGCGGTGT—
CCGCCTCGCCCGCGCCTA

>Ncom04
-------------CGGTGCACTTTCCGCCGGGGCACGTC---------
AGCGGCGATTGCTCTCATCCCTCGAGGCG-----------GCAGAGGGTTGCC-------
TCCTCGGA-------GGTCAGGTGGCCGGGA---------------------------------------------------
CGGCCTTGTGGTCGTCTCGGAGCGT----------A-TACTTT--GCCGTTGGGG--
GCGTGGAAGCG------------------------------------------
ATCGAGGAGTGGGTTCGTTCCGTC-----GGCGTCCAT---TCGCCTG---
GAGCAAAGGCAGGTGCGGGCCAGAGAGTTTGGCCGGGCGGCCCAGG-
CGTCCGGTAGTCTCGAGC--------------GAGTGCTGCC-----------
CCGTCCTCGCGGCGGGCGGCTCACTCGAGTTCTG---CCGGCGTGCGG----------------
-------

>Ncom05
-------------CGGTGCACTTTCCGCCGGGGCACGTC---------
AGCGGCGATTGCTCTCATCCCTCGAGGCG-----------GCAGAGGGTTGCC-------
TCCTCGGA-------GGTCAGGTGGCCGGGA---------------------------------------------------
CGGCCTTGTGGTCGTCTCGGAGCGT----------A-TACTTT--GCCGTTGGGG--
GCGTGGGAGCG------------------------------------------
ATCGAGGAGTGGGTTCGTTCCGTC-----GGCGTCCAT---TCGCCTG---
GAGCAAAGGCAGGTGCGGGCCAGAGAGTTTGGCCGGGCGGCCCAGG-
CGTCCGGTAGTCTCGAGC--------------GAGTGCTGCC-----------
CCGTCCTCGCGGCGGGCGGCTCACTCGAGTTCTG---CCGGCGTGCGGTGT--
CCGCCTCGCCCGCGCCTA

>Ncom01
-------------CGGTGCACTTTCCGCCGGGGCACGTC---------
AGCGGCGATTGCTCTCATCCCTCGAGGCG-----------GCAGAGGGTTGCC-------
TCCTCGGA-------GGTCAGGTGGCCGGGA---------------------------------------------------
CGGCCTTGTGGTCGTCTCGGAGCGT----------A-TACTTT--GCCGTTGGGG--

GCGTGGGAGCG----------------------------------------
ATCGAGGAGTGGGTTCGTTCCGTC-----GGCGTCCAT---TCGCCTG---
GAGCAAAGGCAGGTGCGGGCCAGAGAGTTTGGCCGGGCGGCCCAGG-
CGTCCGGTAGTCTCGAGC--------------GAGTGCTGCC-----------
CCGTCCTCGCGGCGGGCGGCTCACTCGAGTTCTG---CCGGCGTGCGGTGT--
CCGCCTCGCCCGCGC---

>Ncom02
-------------CGGTGCACTTTCCGCCGGGGCACGTC---------
AGCGGCGATTGCTCTCATCCCTCGAGGCG-----------GCAGAGGGTTGCC-------
TCCTCGGA-------GGTCAGGTGGCCGGGA---------------------------------------------------
CGGCCTTGTGGTCGTCTCGGAGCGT----------A-TACTTT--GCCGTTGGGG--
GCGTGGGAGCG----------------------------------------
ATCGAGGAGTGGGTTCGTTCCGTC-----GGCGTCCAT---TCGCCTG---
GAGCAAAGGCAGGTGCGGGCCAGAGAGTTTGGCCGGGCGGCCCAGG-
CGTCCGGTAGTCTCGAGC--------------GAGTGCTGCC-----------
CCGTCCTCGCGGCGGGCGGCTCACTCGAGTTCTG---CCGGCGTGCGGTGT--
CCGCCTCGCCCGCGCCTA

>Ncom06
-------------CGGTGCACTTTCCGCCGGGGCACGTC---------
AGCGGCGATTGCTCTCATCCCTCGAGGCG-----------GCAGAGGGTTGCC-------
TCCTCGGA-------GGTCAGGTGGCCGGGA---------------------------------------------------
CGGCCTTGTGGTCGTCTCGGAGCGT----------A-TACTTT--GCCGTTGGGG--
GCGTGGGAGCG----------------------------------------
ATCGAGGAGTGGGTTCGTTCCGTC-----GGCGTCCAT---TCGCCTG---
GAGCAAAGGCAGGTGCGGGCCAGAGAGTTTGGCCGGGCGGCCCAGG-
CGTCCGGTAGTCTCGAGC--------------GAGTGCTGCC-----------
CCGTCCTCGCGGCGGGCGGCTCACTCGAGTTCTG---CCGGCGTGCGGTGT--
CCGCCTCGCCCGCGCC—

>*Oscarella lobularis* (JX462775.1)
GCAAAGCA---CTTGTTGCAGATTCAGTCGCGTCACGT------------CG--------------
TCGGGAGTCG--G-----------CCGGCGGATCCAATCGGACC---
GTCGTCGGCTCGACGGCGGCGAGGCACGCCG----CAATTCTGTCCCGTG-----C--
GAGCCAACG---------------------TCGGTTCG------TCGACGACGTTCGGCTGTCGC-
GTCAGGTGTCTCGTGACTTCGGTCGGGAGCGTTATAGTCG-
CGATAGCAAGGGCGTC-GAGACGGAG----CGAGGAGACGAAATG-----
TCGGTGGCGAA-------CCGCGCTCGCT--TTCGGGCG---GGCTA------
GAGTCGTTCGACGCGCGTCGTG-------------CGT---CGCGTTT---TGGACTG---------
CGTGCAGTGCTCGATGCGATCG-------TG----------
CGGGCGCGTCGTTTCGTCGTCATAAGCGGTTCGA

>*Oscarella bergenensis* (JX462771.1)
GCAAAGCA---CTCGTTGCAGATTCAGTCGCGTCACGT------------CG--------------
TCGGGAGTCG--G-----------TTGGCGGATCCAATCGGACC---

GTCGTCAGCTCGTCGGTGGCGAGGCGCGGCG----CAATTCTGTCCCGTG-----C--
GAGCCAACG---------------------TCGGTTCG------TCGACGACGTTCGGCTGTTGC-
GTCAGGTGTCTCGTGGCTTCGGTCGGGAGCGTTATAGGCG-
TGATAGCAAGGGCGTC-GAGACGGAC----CGAGGAGACGAAATG-----
TCGGTGGCGAA-------CCGCGCTCGCT--TTCGGGTG---GGCTA------
GAGTCGTTCGACGCGCGTCGTG-------------CGT---CGCGTTT---TGGACTG---------
CGTGCAGTGCTCGATGCGATCG-------TG----------
CGGGCGAGTCGTTTCGTCGTCATAAGCGGTTCGA

>*Neopetrosia carbonaria* (KC869628.1)
GGCGGTTG---GGCGAGCGGAGTTCATCCGGACTTCGAG----AG----
GAGTGCGGTGCATTTTCCGATCGTCTATGGTAACCGTCAGCGAGAGGTGAGCCCG
TCGGTTTTCGGTCGGCG--GAGGAGGTGGCGGGACTCGGTTTGCGAGA-
TCCGTTC-------GTAGAGCTCCTACCG----------------------------
AGCGAGCGGCGTGGCGATCCTCTGGGATGTTGGAGCGTCTT---
TTCGCTCTTGGGAA-----------GTAGGTGCGGGTCT---TCCCGTG-----AGGGAA----------
---------------GGTTGCG---------GGCACCGGGTG-------G------TTGTCGTTCGGTCG----T-
TTCGGTGGACGTGTGCGGCGGT-CGGG--TGA---
ATTCTCGTCGGTTGTGCAGTTTTCGGTTCGGGTG-------GTCGAGTTGCC-CGGG----
TGGT------CACGTTGCCGCGCCCA

>*Neopetrosia subtriangularis* (KC869609.1)
GGTTGTCGTGCTGCATGTGGAGTTCAGCCGGTGTC-GAA----AG----CCGCT-
GGTGCAGTCTCCCATGG---AGCTGGACCGTCAGTTGG-
GAAGAGACTGGAGGCCTGTTGATG--------TCGGTGGTTGGG-----------AGG-
TATGCCC-------
TGGGGTCGCCTTCGGGTGCCCCAGGGTGACATATAGCCATTCCGCCGTGGTCTCT
GTCGTCTTCCAGG-TGTGATTGGGACGACTG---TTCGTCCGAAATGGG-----------
AAGGTGCAGTCTTCCTCCTTGTG-----
AGGGAGTTCGGGCCGGCTCGGCGGTCGCGTTGCGTTGC----------GGCACCGTGTT-
----------------CCGTT---------CT-
CTCCCCATTCTGCCTCGGCAAGTGGGGTTGGAGTGCGGCACG----GACTGCAT-----
TTGACAGCGT------------CCCTAC-CGGG----TCGT-GATCCAAGTTGACTGTGCCCA

>*Leucetta chagosensis* (AY563543.1)
-GCTAGCAATGTGCCACGACGGTTCAGCCGTTTTG-GGC-----G----GTGAG-
GTGGCTG---CTGTCGG---A-----TCCGA------------------A-GGACCGCTGTAGTCG--
TCGTCGCTGTCCGAA-----------ATG-------C-------GGTGCACTCTTTCGTGGTACGC-----
GTC--AGCGTCGGTTCGCAGCGGGCGATAAGGCC----TT---
GTGGAAGGTAGCTAGGGGCCTTGCCCCTAGTGTT-ATAGACGC---
AGGAGCTGGGCTC-GC-----------------TACGGACCG---AGGAGAGAGAGTATCGTTGC-
---------AGCGCATG--------------------CCGC--AAGGCTAGCT-ACCTGCCGTCTGTCTC-
GCTTCACGGAC-TGAGGACTGCACG----CAGTGCTC-----GGACTGGCGG-------
AGTGGTCCGGC--G-G----TGGA-GTG---GTATAAATGCGCTCA

>Lcha05

GGCTAGCAATGTGCCACGACGGTTCAGCCGTTTTG-GGC-----G----GCGAG-
GTGGCTG---CTGTCGG---A-----TCCGA-----------------A-GGACCGCTGTAGTCG--
TCGTCGCTGTCCGAA-----------ATG-------C-------GGTGCACTCTTTCGTGGTACGC-----
GTC--AGCGTCGGTTCGCAGCGGGCGATAAGGCC----TT---
GTGGAAGGTAGCTAGGGG-CTTGCCCCTAGTGTT-ATAGACGC---
AGGAGCTGGGCTC-GC----------------TACGGACCG---AGGAGAGAGAGTATCGTTGC-
---------AGCGCATG-------------------CCGC--AAGGCTAGCT-ACCTGCCGTCTGTCTC-
GCTTCACGGAC-TGAGGACTGCACG----CAGTGCTC-----GGACTGGCGG-------
AGTGGTCCGGC--G-G----TGGA-GTG---GTATAAATGC-----

>Lcha06
GGCTAGCAATGTGCCACGACGGTTCAGCCGTTTTG-GGC-----G----GCGAG-
GTGGCTG---CTGTCGG---A-----TCCGA-----------------A-GGACCGCTGTAGTCG--
TCGTCGCTGTCCGAA-----------ATG-------C-------GGTGCACTCTTTCGTGGTACGC-----
GTC--AGCGTCGGTTCGCAGCGGGCGATAAGGCC----TT---
GTGGAAGGTAGCTAGGGG-CTTGCCCCTAGTGTT-ATAGACGC---
AGGAGCTGGGCTC-GC----------------TACGGACCG---AGGAGAGAGAGTATCGTTGC-
---------AGCGCATG-------------------CCGC--AAGGCTAGCT-ACCTGCCGTCTGTCTC-
GCTTCACGGAC-TGAGGACTGCACG----CAGTGCTC-----GGACTGGCGG-------
AGTGGTCCGGC--G-G----TGGA-GTG---GTATAAATGCGCTCA

>Lcha01
--------ATGTGCCACGACGGTTCAGCCGTTTTG-GGC-----G----GCGAG-GTGGCTG---
CTGTCGG---A-----TCCGA-----------------A-GGACCGCTGTAGTCG--
TCGTCGCTGTCCGAA-----------ATG-------C-------GGTGCACTCTTTCGTGGTACGC-----
GTC--AGCGTCGGTTCGCAGCGGGCGATAAGGCC----TT---
GTGGAAGGTAGCTAGGGG-CTTGCCCCTAGTGTT-ATAGACGC---
AGGAGCTGGGCTC-GC----------------TACGGACCG---AGGAGAGAGAGTATCGTTGC-
---------AGCGCATG-------------------CCGC--AAGGCTAGCT-ACCTGCCGTCTGTCTC-
GCTTCACGGAC-TGAGGACTGCACG----CAGTGCTC-----GGACTGGCGG-------
AGTGGTCCGGC--G-G----TGGA-GTG---GTATAAA--------

>Lcha02
GGCTAGCAATGTGCCACGACGGTTCAGCCGTTTTG-GGC-----G----GCGAG-
GTGGCTG---CTGTCGG---A-----TCCGA-----------------A-GGACCGCTGTAGTCG--
TCGTCGCTGTCCGAA-----------ATG-------C-------GGTGCACTCTTTCGTGGTACGC-----
GTC--AGCGTCGGTTCGCAGCGGGCGATAAGGCC----TT---
GTGGAAGGTAGCTAGGGG-CTTGCCCCTAGTGTT-ATAGACGC---
AGGAGCTGGGCTC-GC----------------TACGGACCG---AGGAGAGAGAGTATCGTTGC-
---------AGCGCATG-------------------CCGC--AAGGCTAGCT-ACCTGCCGTCTGTCTC-
GCTTCACGGAC-TGAGGACTGCACG----CAGTGCTC-----GGACTGGCGG-------
AGTGGTCCGGC--G-G----TGGA-GTG---GTATAAA--------

>Lcha03
-----------TGCCACGACGGTTCAGCCGTTTTG-GGC-----G----GCGAG-GTGGCTG---
CTGTCGG---A-----TCCGA-----------------A-GGACCGCTGTAGTCG--

TCGTCGCTGTCCGAA-----------ATG-------C-------GGTGCACTCTTTCGTGGTACGC-----
GTC--AGCGTCGGTTCGCAGCGGGCGATAAGGCC----TT---
GTGGAAGGTAGCTAGGGG-CTTGCCCCTAGTGTT-ATAGACGC---
AGGAGCTGGGCTC-GC-----------------TACGGACCG---AGGAGAGAGAGTATCGTTGC-
---------AGCGCATG--------------------CCGC--AAGGCTAGCT-ACCTGCCGTCTGTCTC-
GCTTCACGGAC-TGAGGACTGCACG----CAGTGCTC-----GGACTGGCGG-------
AGTGGTCCGGC--G-G----TGGA-GTG---GTATAAA--------

>Lcha04
GGCTAGCAATGTGCCACGACGGTTCAGCCGTTTTG-GGC-----G----GCGAG-
GTGGCTG---CTGTCGG---A-----TCCGA------------------A-GGACCGCTGTAGTCG--
TCGTCGCTGTCCGAA-----------ATG-------C-------GGTGCACTCTTTCGTGGTACGC-----
GTC--AGCGTCGGTTCGCAGCGGGCGATAAGGCC----TT---
GTGGAAGGTAGCTAGGGG-CTTGCCCCTAGTGTT-ATAGACGC---
AGGAGCTGGGCTC-GC-----------------TACGGACCG---AGGAGAGAGAGTATCGTTGC-
---------AGCGCATG--------------------CCGC--AAGGCTAGCT-ACCTGCCGTCTGTCTC-
GCTTCACGGAC-TGAGGACTGCACG----CAGTGCTC-----GGACTGGCGG-------
AGTGGTCCGGC--G-G----TGGA-GTG---GTATAAA--------

>*Leucetta avocado* (KC869542.1)
GGCTAGCAATGTGCCACGACGGTTCAGCCGTTTTG-GGC-----G----GCGAG-
GTGGCTG---CTGTCGG---A-----TCCGA------------------ATGGACCGCTGTAGTCG--
TCGTCGCTGTCCGAG-----------ACG-------C-------GGTGCACTCTTTCGTGGTACGC-----
GTC--AGCGTCGGTTCGCAGCGGGCGAAAAGGCC----TT---
GTGGAAGGTAGCTCGGGG-CTTGCTCTGAGTGTT-ATAGACGC---
AGGTGCTGGGTCT-GC-----------------TGCGGACCG---AGGAGAGAGAGTATCGTTGC-
---------AGCGCATG--------------------CCGC--AAGGCTAGCT-ACCCGTCGTCTGGCTC-
GCTTCACGGAC-CGAGGACTGCACG----CAGTGCTT-----GGACTGGCGG-------
AGTGGTCAGGC--G-G----TGGA-GTG---GTATAAATGCGCTCA

>*Leucetta primigenia* (KC869645.1)
GGCTAGCAATGTGCCACGACGGTTCAGCCGTTTTG-GGC-----G----GTGAG-
GTGGCTG---CTGTCGG---A-----TCCGA------------------ATGGACCGCTGTAGTCG--
TCGTCGCTGTCCGAG-----------ACG-------C-------GGTGCACTCTTTCGTGGTACGC-----
GTC--AGCGTCGGTTCGCAGCGGGCGATGAGGCC----TT---
GCGGAAGGTAGCTCGGGG-CTTGCTCTGAGTGTT-ATAGACGC---
AGGTGCTGGGTCC-GC-----------------TGCGGACCG---AGGAGAGAGAGTATCGTTGC-
---------AGCGCATG--------------------CCGC--AAGGCTAGCT-ACCCGCTGTCTGGCTC-
GCTTCACAAAC-TGAGGACTGCACG----CAGTGCTC-----GGATTGGCGG-------
AGTGGTCAGGC--G-G----TGGA-GTG---GTATAAATGCGCTTA

>*Leucetta floridana* (KC869538.1)
GGCTAGCAATGTGCCACGACGGTTCAGCCGTTTTG-GGC-----G----GCGAG-
GTGGCTG---CTGTCGG---A-----TCCGA------------------ATGGACCGCTGTAGTCG--
TCGTCGCTGTCCGAG-----------ACG-------C-------GGTGCACTCTTTCGTGGTACGC-----
GTC--AGCGTCGGTTCGCAGCGGGCGATGAGGCC----TT---

GCGGAAGGTAGCTCGGGG-CTTGCTCTGAGTGTT-ATAGACGC---
AGGTGCTGGGTCC-GC-----------------TGCGGACCG---AGGAGAGAGAGTATCGTTGC-
---------AGCGCATG--------------------CCGC--AAGGCTAGCT-ACCCGCCGTCTGGCTC-
GCTTCACGGAC-TGAGGACTGCACG----CAGTGCTC-----GGATCGGCGG-------
AGTGGTCAGGC--G-G----TGGA-GTG---GTATAAATGCGCTCA

>*Leucetta purpurea* (KX499450.1)
GGCTAGCAATGTGCCACGACGGTTCAGCCGTTTTG-GGC-----G----GCGAG-
GTGGCTG---CTGTCGG---A-----TCCGA------------------ATGGACCGCTGTAGTCG--
TCGTCGCTGTCCGAG-----------ACG-------C-------GGTGCACTCTTTCGTGGTACGC-----
GTC--AGCGTCGGTTCGCAGCGGGCGATGAGGCC----TT---
GCGGAAGGTAGCTCGGGG-CTTGCTCCGAGTGTT-ACAGACGC---
AGGTGCTGGGTCC-GC-----------------TGCGGACCG---AGGAGAGAGAGTATCGTTGC-
---------AGCGCATG--------------------CCGC--AAGGCTAGCT-ACCCGCCGTCTGGCTC-
GCTTCACGGAC-TGAGGACTGCACG----CAGTGCTC-----GGATCGGCGG-------
AGTGGTCAGGC--G-G----TGGA-GTG---GTATAAATGCGCTCA

>*Leucetta foliate* (KX499451.1)
GGCTAGCAATGTGCCACGACGGTTCAGCCGTTTTG-GGC-----G----GCGAG-
GTGGCTG---CTGTCGG---A-----TCCGA------------------ATGGACCGCTGTAGTCG--
TCGTCGCTGTCCGAG-----------ACG-------C-------GGTGCACTCTTTCGTGGTACGC-----
GTC--AGCGTCGGTTCGCAGCGGGCGATGAGGCC----TT---
GCGGAAGGTAGCTCGGAG-CTTGCTCCGAGTGTT-ATAGACGC---
AAGTGCTGGGTCC-GC-----------------TGCGGACCG---AGGAGAGAGAGTATCGTTGC-
---------AGCGCATG--------------------CCGC--AAGGCTAGCT-ACCTGCCGTCCGGCTC-
GCTTCACGGAC-TGAGGACTGCACG----CAGTGCTC-----GGACCGGCGG-------
AGTGGTCAGGC--G-G----TGGA-GTG---GTATAAATGCGCTCA

>*Leucosolenia complicata* (MH385270.1)
GGCTAGCAATGCACGTTGGTGGTTCAGCCGGCTAG-TCT----TG----TCGAG-
TCGTCGT---TTCGCGG---A-----TCCGA------------------ATGGACCGCGGGCGGCG--
GCATCGGTGGGCGAG-----------CTG-------T-------GGTGCACTCCATCGGCGTGCGC---
--GTC--AGCGTCGGTTCTCGACGGGCGATAAGGTG---CGTG-
CGGGGAAGGTAGCTCGGT-CTTCGGACGGAGTGTT-
ATAGACCCGCTCGGCGCTGGGTTC-GTC----------------GTGGGACCG---
AGGAGAGAGATGATCGTTGC----------GGCACCTG--------------------
CCTTTCGGGGCTAGCC-AGCTGCTGTCAGGTGCGGCGTCACACGT-
TGAGGACTGCACG----CAGTGCTT-----GGTGTGGAGG-------CGTGATCTGGC--CGG--
--TGGT-GTG—GGTATAGAGGTGCCTA

>*Leucosolenia complicate* (MH385268.1)
GGCTAGCAATGCACGTTGGTGGTTCAGCCGGCTAG-TCT----TG----TCGAG-
TCGTCGT---TTCGCGG---A-----TCCGA------------------ATGGACCGCGGGCGGCG--
GCATCGGTGGGCGAG-----------CTG-------T-------GGTGCACTCCATCGGCGTGCGC---
--GTC--AGCGTCGGTTCTCGACGGGCGATAAGGTG---CGTG-
CGGGGAAGGTAGCTCGGT-CTTCGGACGGAGTGTT-

ATAGACCCGCTCGGCGCTGGGTTC-GTC----------------GTGGGACCG---
AGGAGAGAGATGATCGTTGC----------GGCACCTG--------------------
CCTTTCGGGGCTAGCC-AGCTGCTGTCAGGTGCGGCGTCACACGT-
TGAGGACTGCACG----CAGTGCTT-----GGTGTGGAGG-------CGTGATCTGGC--CGG--
--TGGT-GTG—GGTATAGAGGTGCCTA

>*Sycon ciliatum* (MH385303.1)
GGCTAGCAATGCACGTCTTTGGTTCAGCCGGCTAG-TCT----TG----TCGAG-
TTGCCGT---TGTGTGG---A-----TCCGA------------------ATGGACTGCCTGCGGTA--
ACGGCGGTGGGCGAG-----------CTA-------T-------GGTGCACTCCAAGGGCGTGCGC---
--GTC--AGCGTCGGTTCTCGACGGACGACAAGGTA---CGT---
GGGAAAGGTAGCTCGGT-CTTCGGACGGAGTGTT-ACAGACCCGT---
GTGCTGGGTTC-GTT----------------GTGGGACCG---AGGAGAGAGATGATCGCTGC-----
-----GGCACCTG--------------------CCTCTCGGGGCTAGCC-AGCTGTCGTCAGGTGT-
GCGCCAGGTGC-CGAGGACTGCACG----CAGTGCTC-----GGTGTTGAGG-------
TGTGATCTGGT---GG----TGGT-GTG--GGTATAGAGGTGCCTA

>*Sycon ciliatum* (MH385302.1)
GGCTAGCAATGCACGTCTTTGGTTCAGCCGGCTAG-TCT----TG----TCGAG-
TTGCCGT---TGTGTGG---A-----TCCGA------------------ATGGACTGCCTGCGGTA--
ACGGCGGTGGGCGAG-----------CTA-------T-------GGTGCACTCCAAGGGCGTGCGC---
--GTC--AGCGTCGGTTCTCGACGGACGACAAGGTA---CGT---
GGGAAAGGTAGCTCGGT-CTTCGGACGGAGTGTT-ACAGACCCGT---
GTGCTGGGTTC-GTT----------------GTGGGACCG---AGGAGAGAGATGATCGCTGC-----
-----GGCACCTG--------------------CCTCTCGGGGCTAGCC-AGCTGTCGTCAGGTGT-
GCGCCAGGTGC-CGAGGACTGCACG----CAGTGCTC-----GGTGTTGAGG-------
TGTGATCTGGT---GG----TGGT-GTG--GGTATAGAGGTGCCTA

>*Neopetrosia rosariensis* (KC869499.1)
GGTTGTCGTGCTCCGCTGCGAGTTCAATTGGTGGAGGTCGGAAACGGCCACTGCC
AGTGCAATCTCC---TGTGGAGTTGGACCGTCAGTTGGGGAGAGACGGA---
GTCCTGTG--------TAGTCGGCGGTCGGG-----------AGG-TGCGCCCCGGGT------TAC--
CCTCGGGTGCCCGTGGTGAC-
GTATAGCCGCTCCGTCGTGGGCTCTGTCGTCTTCCAGG-
TGCGATTCGGACGTCTG---TTCGTCCA--GAA---------TGGGAAGGTGCAGTCTTCC---
CCGTTCGCG-GGGGAGTTCGGGCCATCTCGGCGGTCGCGCT----GCCT---------
TGCGGTGT-CG-------T------GTTCCGCTCTCTCCCCGCTTTGCCT----------
CGGCGGGT-GGGGCAGAGTGCGGCACG---GCC-TGCATTGGTC-----------------
GGTGTCCCGTC-CGGG----CTGT-GATCCAAGTCGACTGTGCCGA

>*Neopetrosia rosariensis* (KC869457.1)
GGTTGTCGTGCTCCGCTGCGAGTTCAATTGGTGGAGGTCGGAAACGGCCACTGCC
AGTGCATTCTCC---TGTGGAGTTGGACCGTCAGTTGGGGAGAGACGGA---
GTCCTGTG--------TAGTCGGCGGTCGGG-----------AGG-TGCGCCCCGGGT------TAC--
TCTCGGGTGCCCGTGGTGAC-
GTATAGCCGCTCCGTCGTGGGCTCTGTCGTCTTCCAGG-

TGCGATTCGGACGTCTG---TTCGTCCG--GAA---------TGGGAAGGTGCAGTCTCCC---
TCGTTCGCG-GGGGAGTTCGGGCCATCTCGGCGGTCGCGCC----GCCT---------
TGCGGTGT-CG-------T------GTTCCGCTCGCTCCCCGCTTTGCCT----------
CGGCGGGT-GGGGCAGAGTGCGGCACG---TCT-CGCATTGGTC----------------
GGTGTCCCGTC-CGGG----GTGT-GATCCAAGTCGACTGTGCCGA

>*Haliclona tubifera* (KC869461.1)
GGTTGTCGTGCTCCGGTTGG-GTTCATCTGGTGGTGT--------
TCACGCCGACAGTGCATTCCCAGCCGG--A-GTTGGACCGTCAGTTGGGAAGAGA---
TCGGGGCTTGTAT--------TGTCGGCGTTGGGG-----------AGG-TAGGTCTC-----
GTGGTTGCCC--TCGGGTGCCCG--
GGATCACATATAGCCCTGCGCCGTGGTCTCGGTCGTCTTTCAGG-TGCGTT--
GGACGGCTA---CTCGTCCA--TAA---------TTGGTAGGTGCAGTCT----CCCCTTG-----
CGGGAGGTCGGGCCCTCGCCTAGGT-TACCTG-GTGGCG---------TGAAG-CG-TG----
---T------GTTCCGTTCCCTCGCCGTTGG-ACCCAGTCT-GCG--GTGTA--------G-----
GGTGCG---GCACTGCTCGCGAC--------AT-------TGTGTCCGA----AGG----TGTT-
GTGTCC-AAGTTCTGCGCCAA

>*Petrosia ficiformis* (KX688752.1)
GGTGGTCGTGCTTCGTTGGG-
GTTCATCGGACTTGCCTGCCATTGCAGAGCTGTCCGAGCATTCTCCCTCTGAAAGT
TGGAACCGTCAGTCGGGGAGGGTCGTGCAGGACTGATTTGC-----
GTGTCAGTGGTCGGG-----------TAG-
GTTGCCTCTGGACGTGAAAGCTTGCTTGAGTGTTTGGAGGAGCTTATAAGTCGTTC
CGCTGTGACCTGCGTCGTCTCCCATT-TGCGAG--GTG--TCGA---
ATCGCCCCTGAAA---------
GGGGAAGGTGCAGACTTTCTCCACTAACGTTGCGAGAGGGTTGAGAAGCTCCGG
GAGGTAGCT--GTGGCC---------TGTTTCAG-AG-------C------GTTCCGT-
CCTTTGCCTTAGTAGTTCAGTCT-GCTCGGGGAAC-GGGAAGGCGTTCGAAGCA---
GCTGTTCACTGT-C--------CC-------TGTTTACC-----TGG----CAGC-TTTTCA-
AGGAACTGTGCCGA

Data S3. Aligned and trimmed amino acid sequences of bona fide NLRs of sponges and other metazoans.

>LcomNLRX1
RLVAIV-SGAGSGKSTL-CLKILEEFGEKG---------------THPACAQRLKSRFEF
P-ILIR-CRELD--GG--------ES--------LE--ELLGLQCSD-------LDFTDI
ERKSVIKHLR----HH--ANKVLLVVDGLDE---AIPEALGSN-----------------
--------SELRKL--------L--DGA---------KSLVGASVILTGR-L------CD
AIKDVVISRT------RA-VCYALAGFSGQAFKDYGDDEFRAA

>LcomNLRX2
RLVAIV-SGAGSGKSTL-CLKILEEFGEKG---------------THPACAQRLKSRFEF
P-ILIR-CRELD--GG--------ES--------LE--ELLGLQCSD-------LDFTDI
ERKSVIKHLR----HH--ANKVLLVVDGLDE---AIPEALGSN-----------------
--------SELRKL--------L--DGA---------KSLVGASVILTGR-L------CD
AIKDVVISRT------RA-VCYALAGFSGQAFKDYGDDEFRAA

>LcomNLRX3
RLVAIV-SGAGSGKSTL-CLKILEEFGEKG---------------THPACAQRLKSRFEF
P-ILIR-CRELD--GG--------ES--------LE--ELLGLQCSD-------LDFTDI
ERKSVIKHLR----HH--ANKVLLVVDGLDE---AIPEALGSN-----------------
--------SELRKL--------L--DGA---------KSLVGASVILTGR-L------CD
AIKDVVISRT------RA-VCYALAGFSGQAFKDYGDDEFRAA

>LcomNLRX4
RLVAIV-SGAGSGKSTL-CLKILEEFGEKG---------------THPACAQRLKSRFEF
P-ILIR-CRELD--GG--------ES--------LE--ELLGLQCSD-------LDFTDI
ERKSVIKHLR----HH--ANKVLLVVDGLDE---AIPEALGSN-----------------
--------SELRKL--------L--DGA---------KSLVGASVILTGR-L------CD
AIKDVVISRT------RA-VCYALAGFSGQAFKDYGDDEFRAA

>LcomNLRX5
RLVAIV-SGAGSGKSTL-CLKILEEFGEKG---------------THPACAQRLKSRFEF
P-ILIR-CRELD--GG--------ES--------LE--ELLGLQCSD-------LDFTDI
ERKSVIKHLR----HH--ANKVLLVVDGLDE---AIPEALGSN-----------------
--------SELRKL--------L--DGA---------KSLVGASVILTGR-L------CD
AIKDVVISRT------RA-VCYALAGFSGQAFKDYGDDEFRAA

>LcomNLRX8
RLVAIV-SGAGSGKSTL-CLKILEEFGEKG---------------THPACAQRLKSRFEF
P-ILIR-CRELD--GG--------ES--------LE--ELLGLQCSD-------LDFTDI
ERKSVIKHLR----HH--ANKVLLVVDGLDE---AIPEALGSN-----------------
--------SELRKL--------L--DGA---------KSLVGASVILTGR-L------CD
AIKDVVISRT------RA-VCYALAGFSGQAFKDYGDDEFRAA

>LcomNLRX10

RLVAIV-SGAGSGKSTL-CLKILEEFGEKG---------------THPACAQRLKSRFEF
P-ILIR-CRELD--GG--------ES--------LE--ELLGLQCSD-------LDFTDI
ERKSVIKHLR----HH--ANKVLLVVDGLDE---AIPEALGSN-----------------
--------SELRKL--------L--DGA---------KSLVGASVILTGR-L------CD
AIKDVVISRT------RA-VCYALAGFSGQAFKDYGDDEFRAA

>LcomNLRX11
RLVAIV-SGAGSGKSTL-CLKILEEFGEKG---------------THPACAQRLKSRFEF
P-ILIR-CRELD--GG--------ES--------LE--ELLGLQCSD-------LDFTDI
ERKSVIKHLR----HH--ANKVLLVVDGLDE---AIPEALGSN-----------------
--------SELRKL--------L--DGA---------KSLVGASVILTGR-L------CD
AIKDVVISRT------RA-VCYALAGFSGQAFKDYGDDEFRAA

>LcomNLRX12
RLVAIV-SGAGSGKSTL-CLKILEEFGEKG---------------THPACAQRLKSRFEF
P-ILIR-CRELD--GG--------ES--------LE--ELLGLQCSD-------LDFTDI
ERKSVIKHLR----HH--ANKVLLVVDGLDE---AIPEALGSN-----------------
--------SELRKL--------L--DGA---------KSLVGASVILTGR-L------CD
AIKDVVISRT------RA-VCYALAGFSGQAFKDYGDDEFRAA

>LcomNLRX13
RLVAIV-SGAGSGKSTL-CLKILEEFGEKG---------------THPACAQRLKSRFEF
P-ILIR-CRELD--GG--------ES--------LE--ELLGLQCSD-------LDFTDI
ERKSVIKHLR----HH--ANKVLLVVDGLDE---AIPEALGSN-----------------
--------SELRKL--------L--DGA---------KSLVGASVILTGR-L------CD
AIKDVVISRT------RA-VCYALAGFSGQAFKDYGDDEFRAA

>LcomNLRX14
RLVAIV-SGAGSGKSTL-CLKILEEFGEKG---------------THPACAQRLKSRFEF
P-ILIR-CRELD--GG--------ES--------LE--ELLGLQCSD-------LDFTDI
ERKSVIKHLR----HH--ANKVLLVVDGLDE---AIPEALGSN-----------------
--------SELRKL--------L--DGA---------KSLVGASVILTGR-L------CD
AIKDVVISRT------RA-VCYALAGFSGQAFKDYGDDEFRAA

>LcomNLRX15
RLVAIV-SGAGSGKSTL-CLKILEEFGEKG---------------THPACAQRLKSRFEF
P-ILIR-CRELD--GG--------ES--------LE--ELLGLQCSD-------LDFTDI
ERKSVIKHLR----HH--ANKVLLVVDGLDE---AIPEALGSN-----------------
--------SELRKL--------L--DGA---------KSLVGASVILTGR-L------CD
AIKDVVISRT------RA-VCYALAGFSGQAFKDYGDDEFRAA

>LcomNLRX6
-SCLVL-SGAGTGKTTL-CRYILKAYTSAV--CQRS-LLT---------------ERCPY
P-VYIR-CQDME--RFH------AND--------WM--TVLGLN----E-DHLCRILSAE
DRQQIMRYFL-D--EN--SDKLLFILDGADE---VEGESVLPQ-------G---------
--------AILKKD--------F--LRK-------DREEMLQSRVIITSR-P------CP

QASELVDACS---------AWYSLAGFDSEQLRDYLYGRLG--

>LcomNLRX7
------------------------------SRHHAG-CLP---------------KRFKC
V-IHVE-CRDVE--VIN------SVD--------WT--VFLGLH----D-E--SLNLDAE
ERECVLRYV--Q--EH--AAETLVIIDGLDE---AGKEGLRPE-------S---------
--------AALSFL--------R--RQR-------RKSKLMSSSYIVTGR-P------CQ
QMYDLLSAVD---------CHYELHGFSPAQLHTFVSDRLGET

>LcomNLRX9
------------------------------SRHHAG-CLP---------------KRFKC
V-IHVE-CRDVE--VIN------SVD--------WT--VFLGLH----D-E--SLNLDAE
ERECVLRYV--Q--EH--AAETLVIIDGLDE---AGKEGLRPE-------S---------
--------AALSFL--------R--RQR-------RKSKLMSSSYIVTGR-P------CQ
QMYDLLSAVD---------CHYELHGFSPAQLHTFVSDRLGET

>LcomNLRX16
RGVIVM-GGAGKGKTMV-CRRALSDFTEEESRRFTG-CLS---------------EHFTC
V-IHIE-CRDVE--VAN------SPD--------WS--VFLGLQ----D-M--SLGLDTE
ERGWVMDYL--R--DH--SEDVLLVIDGVDE---VGKEGFAKE-------S---------
--------AMLAFL--------K--RLP-------RKTLLLKSSFLLTSR-P------CQ
QAYDLVPACD---------AHYQVRGFSLNQLSTFLNDRLGE-

>LcomNLRX18
RGVIVM-GGAGKGKTMV-CRRALSDFTEEESRRFTG-CLS---------------EHFTC
V-IHIE-CRDVE--VAN------SPD--------WS--VFLGLQ----D-M--SLGLDTE
ERGWVMDYL--R--DH--SEDVLLVIDGVDE---VGKEGFAKE-------S---------
--------AMLAFL--------K--RLP-------RKTLLLKSSFLLTSR-P------CQ
QAYDLVPACD---------AHYQVRGFSLNQLSTFLNDRLGEN

>LcomNLRX24
RGVIVM-GGAGKGKTMV-CRRALSDFTEEESRRFTG-CLS---------------EHFTC
V-IHIE-CRDVE--VAN------SPD--------WS--VFLGLQ----D-M--SLGLDTE
ERGWVMDYL--R--DH--SEDVLLVIDGVDE---VGKEGFAKE-------S---------
--------AMLAFL--------K--RLP-------RKTLLLKSSFLLTSR-P------CQ
QAYDLVPACD---------AHYQVRGFSLNQLSTFLNDRLGE-

>SciINLRX17
----------------------------RK--FRRD-AVL---------------SNFNY
V-VHLP-CRDVD--RVC------SYN--------WS--TLLGLD----E-Q--CLHLDTH
QQDQMIEYL--E--DQ--SQQVLFIVDGLDE---IGKDVLEER-------S---------
--------ALSALV--------H--RE-----------RFADSRILLTSR-P------CQ
LASTLAKECE---------EHYWLAGFNDHQLERYCLRNLGED

>SciINLRX1
-RCMLM-SGAGKGKTMA-CRRVMADYSGDE--HAPG-LMQ---------------EHFDF

V-IPIS-CRDTD--RVS------TSE--------WT--EFLGLD----H-H--ALDITSD
EREEVLRYL--S--AN--SERVLLLLDGVDE---GGKDAFQYT-------S---------
--------AACQFM--------D--ET--------KKNRLVNASVVVTSR-P------CN
KATDLVRQCA---------AYYRLTGFTDAQLRDFCCQQLG--

>SciINLRX2
-SCMLM-SAAGKGKTMA-CRRVMADYANTK---TSS-ILR---------------SQFDF
V-IPIS-CRGTE--RVS------TDC--------WS--KFLGLD----H-S--ALHLTTT
EREEVLRYL--S--AH--SDRVLILLDGVDE---GGKAAFRST-------S---------
--------AAHHFM--------A--ET--------NKNLLVNASVIITSR-P------CE
KATDLVAQCM---------VHYRLTGFTDTQLKDFCCQQLQLE

>SciINLRX3
ESCIVI-GNAGQGKTTL-SKRVIADIMETT----TG-NL----------------AKIEF
I-FYVP-CRDLT--RIQ------SAD--------WC--VFLGLD----Q-L----RISTQ
DQSSLLNYL--S--EH--SDQVLIVIDGIDE---LGSGGFGKG-------S---------
--------AAQAVI--------L--RSSSS-----NHSLLPDAIVVATSR-P------CD
EAYHHIQNFR---------LRFRLLGFSTDQLFAYCSRRLGEE

>SciINLRX4
KRCMLI-GGAGQGKTTV-CRKILSEFLEKE----RT-EL----------------RQFKY
V-FYIP-CRSME--RVT------TED--------WI--VLLGLK----S-F------TAD
ESHPVMEYL--A--TH--SAEVLVIFDGVDE---AGP-GFHAS-------S---------
--------AALALI--------K--NNSDDDSSNAMPSRLPDATVIVTSR-P------CQ
LAYDLVPLCT---------LYFRLTGFSETQLKEFCFKHLDRD

>SciINLRX6
KRCMLI-GGAGQGKTTV-CRKILSEFLEKE----RT-EL----------------RQFKY
V-FYIP-CRSME--RVT------TED--------WI--VLLGLK----S-F------TAD
ESHPVMEYL--A--TH--SAEVLVIFDGVDE---AGP-GFHAS-------S---------
--------AALALI--------K--NNSDDDSSNAMPSRLPDATVIVTSR-P------CQ
LAYDLVPLCT---------LYFRLTGFSETQLKEFCFKHLDRD

>SciINLRX7
RSCLVV-AAAGQGKTTL-CRRVIAEVMDAK----AG-PL----------------SKFRF
V-FYIP-CRDAE--RLS------TRN--------WS--EFLGLH----D-D--EMGLNDE
EKKAVMLHL--Q--TR--SDEVLIVLDGIDE---ASIDGLAEG-------T---------
--------AAYALV--------K--RGTSGV----AKTRLLNATVIATSR-Q------CR
GALELSAMCR---------MGFRLKGFSKEQLAKYFIHQLD--

>SciINLRX14
SGCMLI-GSAGQGKTTL-CKRIIADVSDAA--KRSS-PL----------------NKLKY
V-FYIA-CRHTE--RVT------SPD--------WC--VLLGLD----G-L----GVNTE
EQHQLLQYL--S--RH--SDEVLVILDGIDE---AGAHGLGAE-------S---------
--------AAAALL--------Q--RQSDGTS---SRSHLLGAVVVATSR-P------CE
EAHTLVEHFS---------QHYRLVGFSESQLGEFCRQHLGET

>SciINLRX16
SGCMLI-GSAGQGKTTL-CKRIIADVSDAA--KRSS-PL----------------NKLKY
V-FYIA-CRHTE--RVT------SPD--------WC--VLLGLD----G-L----GVNTE
EQHQLLQYL--S--RH--SDEVLVILDGIDE---AGAHGLGAE-------S---------
--------AAAALL--------Q--RQSDGTS---SRSHLLGAVVVATSR-P------CE
EAHTLVEHFS---------QHYRLVGFSESQLGEFCRQHLGET

>SciINLRX8
SSCMVI-GGAGQGKTTF-CKRVIAEVTDAV----SG-PL----------------AKFKF
V-FYIP-CRNLE--RVR------SES--------WI--ELLGLD----A-A--SMDLNGE
EQRSVLRHL--Q--TH--SENVLVILDGIDE---AGGDGLLEQ-------S---------
--------AAKALV--------K---RSGGAS---SASRLLNATVVVTSR-P------CQ
GAYNLVQACK---------RRYRLRGFNESQLEEFCVCQLGD-

>SciINLRX9
SSCMVI-GGAGQGKTTF-CKRVIAEVTDAV----SG-PL----------------AKFKF
V-FYIP-CRNLE--RVR------SES--------WI--ELLGLD----A-A--SMDLNGE
EQRSVLRHL--Q--TH--SENVLVILDGIDE---AGGDGLLLQ-------S---------
--------AAKALV--------K---RS--------ASHLLNATVVITSR-P------CQ
GAYDLVQACK---------LRYRLRGFSESQLHEFCIQQLGE-

>LchaNLRC1
NSCMVV-GPGGVGKTLL-LEWLMVSWAEGR-----L-EEL---------------SEFEL
V-IRLN-RRDAT--ALA------SKT--------AI--GVFEGALRR-Q-C----GVDGA
ELKEIVEHF--R--VN--SRPLLLLVDSPEE---GGEAWSKND-------G---------
----------LQML--------V--EK----------RSLERCSVIITSR-P------CS
LAYNLARWCQ---------LRYYAVGFSDKQLYDLLERILGE-

>LchaCN9
DSCLVV-GPGGVGKTLL-LQWLMVSWAEGR-----L-EEL---------------SEFEF
V-LYLS-GRDVK--MSK------TNT--------VV--DLLQSALEL---Y----GLIGT
ELKDMCEYL--Q--GN--SSRLLVLLDSADE---GGEAWSQSK-------G---------
----------LGML--------L--EK----------RSLEDCSFIVTSR-P------CS
QA-----------------------------------------

>LchaCN6
DSCMII-APGGAGKTLL-LKWLMMQWAEGS-----M-EEL---------------SEFEF
V-IYVR-GRDLA--ALE------SET--------AV--GVLQSALKQ---Y----DLSDN
ELCEIGQYF--K--DN--SSRLLVLLDSADE---CGEAWSH-------------------
------------------------------------------------------------
-------------------------------------------

>LchaCN5
DSCLVI-GPGGVGKTLL-LQWLMVSWAEGR-----A-EEL---------------SEFEL
V-IYVR-GRDTT--ALE------SDT--------GV--GVLQAALKL---Y----DLKEA

ELKDLGEYL--R--GN--SRHLLVLLDSADE---GGQAWLNSK-------G---------
----------LQML--------L--NR----------RSLQECSLIVTSR-P------CS
LAYDLVQVCM---------LQYYAVGFSDAQLDKLLERRLGTA

>LchaCN11
DSCLVI-GPGGVGKTLL-LQWLMVSWAEGG-----V-EEL---------------SKFEF
V-IYVS-GRDER--ALK------SAT--------AI--GVLQSALKR---Y----VSSDA
ELEEIGKYL--R--EN--SSHLLVLLDSADE---GGKAWSQSE-------G---------
----------LQML--------L--ER----------RSLQDCSFIVTSR-P------CS
LAYDLVQLCM---------LRYYVVGFTDEHLDELLGRRLGTD

>LchaNLRC4
DSCLVI-GAGGTGKTLL-LQWLMVSWAEGS-----V-EEL---------------STFEY
V-IYVS-GREAR--ALK------SDT--------AV--GVLQSALKL---Y----GLSDA
DLSGMAEYL--S--QD--SSRLLVLLDSADE---GGDAWSESD-------G---------
----------LRML--------L--DR----------RSLQDCSFIVTSR-P------CS
LAYSLLPLCT---------LRYYNVGFSDERLDELLVRRLGEH

>LchaNLRC5
SSSLLV-GQAGGGKTMT-IRKLLLSWAEEE-----E-AIR---------------KKFDF
V-FYVS-GQDHK--ALN------GKT--------AV--DLLHLE----A-Y----GFDHD
EQRTIGSHL--S--ES--SERVLVVLDGADE---GGDSWGKSE-------G---------
----------VKQL--------L--DR----------KLLHNCTFVITTR-P------SM
RAYELVPYCE---------EHFHLSGLDERRLKELVTRRLGET

>LchaNLRC7
RSCLLL-GQAASGKTLT-LLKLLSCWAERS-----E-EFP---------------NDFEF
V-FYVS-GRDER--ALK------GTS--------FV--DVLQLD----Q-F----GLDEG
ERRQMAIYL--R--EH--SDKVLILFDSVDE---AGEKWGRSD-------G---------
----------VRQI--------L--QR----------KALTKCSFVVTGR-P------CL
KAYELMPRCV---------QHFRLAGLNDCRLDELLRRRLGMD

>LchaCN3
RSCLLL-GHAASGKTLT-MYKLLSCWAERN-----E-EIL---------------KEFEF
V-FYVS-GRDKR--ALE------GTS--------FV--DVLQLE----R-F----GLDKE
EVRQMAKHL--R--EH--SDKVLVLFDGVDE---GGDKWGKSD-------G---------
----------IQQI--------L--QR----------RALRRCSFVATGR-P------CV
KAFELIPRCV---------QHFQLAGLNDYHFEELLRRRLGDK

>LchaNLRC3
NSCLLL-GHAGSGKTLT-MHKLLSCWAERS-----E-EFL---------------NVFEF
V-FYVS-GRDRR--ALE------GAS--------FV--DVLRLE----T-F----GLGKE
EEHQVARYL--Q--DN--SDKVLVFFDGVDE---GGEKWGSND-------G---------
----------IQQI--------L--QR----------KALGRCSFVVTGR-P------CL
KAFELIPRCV---------QHFHLAGLNNSGLEELLRRRLGAE

>LchaNLRX14
KRSVIF-GGGGSGKTLL-CRKILSDWDNIR----LV-NSF---------------KKFWA
V-MYIS-GRNTG--RIH------ATN--------AS--TFLGIN----D-R----NLSDR
QCCEIVGFM--A--EN--SERLLVLIDGWDE---VSGSGLLQG-------G---------
--------TVLTDL--------L--RR---------RGCFSQASILITSR-P------CP
EVFQVLELCK------IS-RFFSLVGLKEKRIRELACRKLGDH

>LchaNLRX4
QRALVI-GPGGSGKSLT-CSKIIADWIDRT--------AF---------------RQFLA
V-VHIS-AKDTD--HWQ------ARS--------SS--DLLCLD----S-H----GYSQS
QQTRLLLQL--R--KR--SEKLLFLIDGSDE---AAGEGLLTP-------G---------
--------TALHDV--------L--NR----------KLFRNASVIITTR-P------SP
SCYNLLNICD---------KCFYLAGFTEKHLNEFVRNRLEP-

>LchaNLRX13
KSALIL-GMSGSGKSLT-VLKILSEWIGEA-----K-SAV---------------KQFGM
V-VLVT-GRETT--RLN------AKA--------VE--DFFQLW----R-L----GYNER
QQRYILDYF--K--ESEDAKSVLILIDGWNE---GGEACLLQD-------G---------
--------TVLKDM--------L--RGE-------RSKLFPGCSVIITSR-P------TE
SIYPLLEECD---------ARYNLLGLSNQQLRQLLSRRLNEK

>LchaNLRX3
-RSLVL-GGGGSGKSLT-CYKILAEWSGDD-----P-SPF---------------KEFDA
V-FYIT-GREQK--RLS------AKS--------RA--QLLRFN----S-M----GFDKE
RERQLVEYF--T--EH--SEKMLFLIDGGDE---LSNGGLLDQ-------G---------
--------LAMRRV--------L--RG----------QLFPKACVIITSR-P------CP
GAYKLLNVCQ---------RHFALVGFTDDNLHHLIKCRLGQQ

>LchaDN1
-SALLF-GEAGSGKSLT-TLKILSEWNKNE--LQST-LPF---------------KKFDM
I-VYVT-GRESI--RLQ------AKE--------IQ--NFWQLW----R-F----GYSGG
QQRHILHHY--E--HH--SEKVLFVIDAWDE---CGNDSPLQE-------D---------
--------TVIMQI--------L--HG----------KLFRNSSVIITSR-P------SP
SAYPLLKVCK---------RRYSLVGFKNKRLRELFDRRLGEQ

>LchaDN2
KSALLF-GEAGSGKSLS-ALKVLSEWSKKE--LLST-SPF---------------KQFDV
V-VYVT-GREST--RMQ------AKD--------VQ--NFWLLW----Q-C----GYSED
QQEHILHHY--E--RH--SEKVLFLIDAWDE---CRSDGLLEK-------D---------
--------TVMMKI--------L--HK----------KLFCNSSVIITSR-P------SP
NA-----------------------------------------

>LchaNLRX6
KSAIIF-GSAGSGKSLT-VLKILSEWWSQN----KT-SPF---------------ERFDV
V-IHIT-GRETS--RIR------TKS--------IQ--ELWQLW----R-M----GYTDH
QQQFIVKHY--S--QH--SEKVLFLVDAWDE---AGDNGLLQE-------D---------

--------TALKRI--------L--HG----------ELFQYSPVIITSR-P------AP
NTYLLLHKCK---------RRYSLVGFNNDRLKELFDRRLEEP

>LchaNLRD4
RSTILF-GSAGSGKSLT-TLKILSEWSSRN----SG-SPF---------------KKFDV
I-VHLT-GKERS--RLR------TTD--------VH--ELWRLQ----R-M----DFSDE
QQLIIIDYF--A--HH--SEKVLFLIDGWDE---AGDDGILQE-------G---------
--------SVLKAI--------L--DG----------NLFPQSSTIITSR-P------VP
TAYPLLEKCR---------YCCLLVGFSDKRLQEFFHQRLGEQ

>LchaNLRD2
QRTLLF-GSAGSGKSLT-MLKILSEWIAKD----SA-SPF---------------KRFDA
I-VYIT-GRERS--RLS------TMD--------VN--DLWRIK----Y-L----GYTEQ
QEEFLLDHF--S--RH--SDKVLFLIDGWDE---CWDDNLLRE-------D---------
--------TVMKQI--------L--CG----------DLFPQSAVIITSR-P------AP
NAYPLLEKCK---------HRYALVGFSDRRLQELFNRRLGEQ

>LchaNLRX16
------------------------------------------------------------
--------RERL--RLI------TKE--------VH--DLWQMK----Q-M----GFNER
QGLFILDHF--S--HH--SDRVLFLLDGWDE---AGDSGVLQE-------D---------
--------SALKQI--------L--DG----------DLFPQSAIIITSR-P------AP
NAYPLLEKCK---------QRYTLVGFNDKRLRELFNRRLGET

>LchaNLRX5
KSALLF-GSAGSGKTLT-TLKILSEWNSKE----TT-SPF---------------QQFDV
I-VYVT-GKERA--RLR------AKD--------IH--DFWRLE----Q-M----GFNEK
QQQFILNHY--S--SL--SEKVLFLIDGWDE---GGDDDLLQG-------D---------
--------SALRKI--------L--QR----------DTFPQSAVIITTR-P------AP
NVHPLLETCR---------HRYSLVGFNDSRLQELFHQRLEES

>LchaNLRX9
-RALIF-GSAGSGKSLT-TLKILSEWTSQS----CA-SLF---------------KRFDA
V-VYIT-GRERS--RLT------TDN--------VS--NLWQLQ----H-N----GFNKQ
QECFLLNHF--S--EC--SDKVLFLVDGWEE---AGEHGLLQE-------D---------
--------MVLNKI--------L--NG----------ELFPQSAVIITSR-P------AP
TAYHLLEKCK---------NRYSLTGFNDRRLQELFNRRLSES

>LchaNLRD3
-SALLF-GSAGSGKSLT-TLKILSEWMDSD----TK-SAF---------------NKFDV
I-IYIA-GRERS--RIS------TSD--------VR--ELWQLE----Q-M----GFNEQ
QQLFFLDHF--L--QN--SSKVLYIIDGWDE---AGDDGLLQE-------N---------
--------TVLKRI--------L--HR----------ELFPESSIIITSR-P------AP
NAFPLLEKCL---------RRYSLVGFNDRRLRELFCYRLGDS

>LchaNLRC2

-SALLF-GLAGSGKSLT-TLKILSEWTSKT----SL-SMF---------------KQFDA
I-IYAT-GKERS--RLR------AMD--------VA--EFWQLQ----Q-M----GFNEQ
QSHFILGYF--L--QN--SERVLFLIDGWDE---AGDDGLLQE-------G---------
--------SVLKNI--------V--DG----------ELFPQSSIVITSR-P------AP
NAYPLLEKCY---------HRYSLVGFNDKRLKELFNRHLNET

>LchaNLRD1
-SALLF-GSAGSGKSLT-TLKILSEWNSES----ST-SAF---------------KQFDA
V-VYLT-GKERS--RLG------AAN--------VQ--DLWRLQ----Q-M----GFSEQ
EELFILNHF--S--QH--SEKVLFLVDGWDE---AGDHGILQD-------D---------
--------SVLKKI--------L--DG----------DLFPQSAIIITTR-P------VP
NAYPLLEKCK---------QRYSLVGFNNTHLQELFHRQMEEA

>LchaNLRX12
HVVAIT-GSAGSGKTTV-LRQLAAAWAK-------L-KCGKEEASNHVPAHLHWVSRYRF
V-LYLD-VSVTD--SG-------NIS--------LK--EIIRS-QLL-P------QNSLL
RVGLLLKEIS-E--NG--GKEVLVLLDGAHI---RIHPILSSS-----------------
---------EIGQL--------L--QC----------QLLPLSTVVVTHN-D------DC
PLPTIYRRD------GRL-LELEVAELTLREREDLVKLYVGDN

>LchaNLRX17
RHIAIV-GSSGAGKSVA-CQLLCSNVSA-------G-VLK---------------ERLSR
V-LCWD-FRDVQ--VRK------ATG--------IT--ELLQI-ATQ--------CSDAT
TCRPLAEELL-A--SR--GKGVLLIFDGLEQ---FSAA------------D---------
--------SVIWSL--------I--NG----------YLLSLSHVVVTTR-P------CG
I-VKRLTNF-------VF-YQLQLDEVLE--------------

>LchaNLRX10
-----------------------SAYLK-------RKAEL---------------SRYDF
V-FVVP-ARNLT--EAN------GDN--------VF--SLL-N-LHH--------YLSAE
EVADVLPHLK----DN--ADRVLILLDGADE---MGSSLS---------KS---------
--------KAVEDL--------L--DG----------PSLSGRTLLVTSR-V------SD
L-ADRFMKE-------AC-SCAMILELTDEQLDSMVRLRLNAK

>ScilNLRX18
-RIGLH-GAPSLGKSQV-CQRLLLNWAY-------R-RQL---------------ARFYL
V-VHWE-LRDAS--VQS------ASS--------LQ--DLLLA-LGV--------QDDM-
-VSSAVPALQ-R--MR--GRGCLFILDGLDE---MSPQTATTTATTTTGAG---------
--------QFVRQL--------M--DG----------RALPEACLLVTSR-P------CA
Q-SEALFAK-------YT-IQLDILGFSDQQAEEFISCHLAT-

>LcomNLRX23
------------------------------------------------------------
------------------------------------------------------------
-----------R--ER--GRGCLFVLDGLEE---LQAGNT---------TS---------
--------TYIRRL--------L--RG----------KVLPDACLLVTSR-A------CS

E-ADQLFTT-------YS-STAEILGFSDSQAESFIRLQIPE-

>LchaDN3
KHVGII-GGTGCGKTNL-CTSITTFYS----------KCW---------------PMFSL
V-LFWR-LYDPS--VQK------AEN--------LK--ELLCA-VGP--------SWTTI
RALRLAETLL-A--TA--GRGVLVILDGVDQ---LEAD-----------EN---------
--------ACIWRL--------L--EG----------SVLQEARLLVTSR-P------CF
L-AKHYFDT-------YD-INLELLGFTEEQVSHFIYHCLGED

>LchaNLRX7
KRVGMY-GSSGCGKTFC-CTALTQLYAS-------Q-QLW---------------KHFRL
V-LFWR-LSDPI--VQE------AKT--------LV--QLLFA-LLP--------SASLR
RRQRLASVLN-T--SN--GKGILIILDGVDQ---LEGG-----------ER---------
--------AFVRHL--------L--SG----------EAFKEACLLVTSR-P------CS
L-AENLFSG-------YN-VQLVLKGYTRKQVVSMLDRRLG--

>LchaNLRC6
QQIGLY-GGAGCGKTAS-CLKALKERSE-------G-RLW---------------NGFQL
V-LLWH-LRKPD--VQR------AST--------LK--DLLRA-LPV--------PLSER
QSTRLSEHLE-E--SD--GRGVLLVLDGIDE---LRRPS----------NN---------
--------AYVVRL--------L--ER----------TRLHRSSILVTSR-P------CY
E-AEQYFKT-------YD-VCYEIIGFTDDQVLSYIHYYLEEE

>LchaCN4
KRVSIY-GGAGCGKTSC-LLKIASMYAE-------K-RLW---------------PEAHA
F-LLWK-LRDRE--VQK------AQN--------LE--QLLRL-LKL--------SLTEE
KCKELAAVLE-A--SE--GEGVILALDGIDE---LDTQ-----------KN---------
--------GIVWSL--------L--DG----------SVLSEACVLATSR-P------SS
I-AKTFFNE-------YD-VNLELLGFTEEQVDQFVEQQFVDK

>LchaCN10
KRVGIY-GGAGCGKTSS-LMKAASMYAN-------G-QLW---------------QGARA
L-LLWK-LRNPD--VQT------ADC--------LE--QLLLQ-LMI--------GLTEQ
QCKHFTEVLL-S--CA--GEGVILALDGVDE---LDAQ-----------QE---------
--------GYIWSL--------L--DG----------SALSKACVLATSR-S------CS
T-AKTYFDK-------YE-VNLELLGFTEEQTGQFIQQQLGDQ

>LchaCN2
QRVGIY-GGAGCGKTSC-LTKAASLYAE-------K-QLW---------------QGART
F-LLWK-LRDPN--VQE------ASS--------LT--QLLRQ-LSL--------DLTEK
KCEELASVLS-S--SD--GEDVILALDGIDE---LDTK-----------VK---------
--------GYVRRL--------L--DG----------SALKKAYVLATSR-P------CS
T-ARTYFDR-------YG-VNLELLGFTEEQVDQFIQQQLGDD

>AqueDN1
RFVLIE-GEPGIGKSTL-AKELVLRWAN-------R-SDK------------LL-SNYDI

V-FFIQ-LRFET--YHK------ATS--------IE--DLFVD-LDN----------QTI
NMTDLNIEIK-K--RK--GAGILWILDGFDE---LPSHLRNNS-----------------
---------ILMTL--------I--KG----------VILPKSTVIVTSR-P------VA
S-DLLLELL----KDDNS-KRISLRGFDSTKIGEFALKYFNDK

>AqueDN5
RFVLIE-GEPGIGKSTL-AKELVLRWAK-------G-SDE------------LM-NNYDI
V-FLIQ-LRFET--YHK------ATS--------IE--HLFVD-LDD----------QSI
NMTELHVEIK-K--RK--GAGILWILDGFDE---LPSHLKSNS-----------------
---------VLMKL--------I--KG----------DNLSKSTVIVTSR-P------VA
S-DQLLHFL----HEHDS-KRISLRGFDSTKIEEYALQYFNDK

>AqueND1
LRVVID-GPPGIGKTTL-CRKLLNMWSN-------G-TLV-------------H-QQYDL
V-LYCP-LRNSK--IAT------ATT--------LA--DLF---EY-----------RCC
EVPIVAKWFE-K--RN--GEGLLIIFDGWDE---LSEQLRRSS-----------------
---------LAASI--------I--HR----------EQLDQCSVIVTSR-S------YA
S-SSLLKMD----T--LS-RHVQVIGFSKKEISKVIIRTLQKD

>AqueDN2
LRVAID-GPPGIGKTTL-CRKLLNMWSK-------G-SHE-------------L-QHYDL
V-LYCP-FRHKA--IAE------ATK--------LV--DLF---VY-----------ESP
KVSKVVDWIL-E--RE--GKGLLIIFDGWDE---LSTQLRESS-----------------
---------LATKI--------I--CR----------NQLVRSSVIVTSR-T------YA
T-ASIYQLE----C--LN-LNVHVIGFAADEINYVIKGMLSQK

>AqueDN3
LRVAID-GPPGIGKTTL-CRKLLNMWSK-------G-SHE-------------F-QHYDI
V-LYCP-FRHKA--VAE------ATE--------LA--DLLTC-AY-----------KSP
KVSEVADWIL-E--RE--GKGLLIIFDGWDE---LSTQLQESS-----------------
---------LATSI--------I--CR----------DQLVYSSVIVTSR-T------YA
T-ASLLQLE----C--LN-QNVHVIGFVAAEIDEVIKGTLSSP

>NcomDN1
-KILLV-GVAGSGKTTI-SWHACQQWAE-------G-NMF---------------QEFEF
L-IYLS-LADPH--IQS------ASS--------FK--DLIGH-PCN------------E
VCNVVAKAIE-G--PK--KKKVCFVMDGWEN---LPTTLQE--------S----------
--------TFLRSLIQG----NA--SE----------V------VIVTSR-P------IA
A-GSIVFSM--------F-FTYNIREFSDDDIAFCAYQYLPTK

>NcomDN2
TKVLLE-GVAGSGKTTV-TWYACRGWAE-------E-EMF---------------HEFDY
L-IHLT-LADPE--LHS------AKT--------IE--DIIPH-PSS------------E
MRKAVADAII-K--LN--GRGCCFILDGWED---LPSQARISS------N----------
--------SILACLLHG----NN--VR----------MALPQCSFLVTSR-P------AV
S-DSLINSV--------S-RVITMEGFPEDAIDSYANRYFCTQ

>HtubNLRX1
-STLIS-GRPGIGKTVL-LTKVCKDWGK-------G-NCL---------------KNIQV
I-IHVS-LRKLH--CKS---P--QPS--------LL--DMIEL-HFT----------DSD
RSKQFCRLIE-A--KG--GKNVCFAFDGLDE---YPLLNTSND------I----------
----------VSEI--------I--GK----------QKLPMASVIVTSR-P------TA
S-HIVKQKM--------K-KHAEIIGFMPHQIESYINEYYTSS

>AqueDN6
KVIIIE-GCPGIGKTTL-ASKLCQEWSE-------K-RQM---------------LEFKL
L-LYIP-LRSPL--MRT--AQ----S--------ID--ELLEY-YG------------DN
YTSNDVLLIK-K--NQ--GRDVVFILDGWDE---LRPSCRSID------M----------
---------FFPGL--------I--YG----------KFLPESTIVVTSR-P------GA
T-IDIRRHA--------N-RIVEILGFTEDQIKQYIMSYFKTN

>NcomNLRX3
KMVLIQ-GAPGVGKTLL-ARCICQRWAK-------G-EIL---------------TEYQL
V-LFVP-LRAFP--ANS---G--SLS--------LH--NIVEL-YLC-----------GS
GIDDAVRELS-E--HN--GKKVLLILEGWDE---LPPELRREF------T----------
---------IFNDL--------V--MG----------TKLPEASIMVTSR-P------IV
V-DELYKTI-------KD-RQIEVLGFKEKQIKEYLKHNLKAE

>NcomNLRX1
SAVLIE-GEQGKGKSMF-SLYLCKKWAK-------G-DML---------------NQFNI
V-LLVP-LHRFS--PES--AL--QLT--------TR--ELIEI-YLP-----------GS
AGSKASNDIE-F--SD--GEKVLFILDGWDE---LTPVLQK-N------G----------
---------FFRDL--------L--LG----------KILTKASVLVTSR-P------TA
A-LQLGQLV--------S-RAYQLLGFSQYEVQRFLRLHVPD-

>NcomNLRX2
HKVLVE-GAPGIGKTML-AQYLSREWAE-------G-RLL---------------KEFNL
V-LLVP-LRRFS--SKS--AD--SLT--------IQ--DLVQI-YLS-----------GD
LGKKASKQLE-L--SG--GEKLLIILEGWDE---LPSELRKDM------T----------
---------LFHDL--------L--IG----------HKLPKASVLVTSR-P------TV
A-GDLYQFV--------N-RRIEVLGFLEEQIEEYVKFHTRDK

>PficDN1
TRILVK-GAPGIGKSSL-AIELCKKWSI-------G-ELG---------------SHFKL
V-ILLR-LKVPK--IQ----K--AKK--------IE--DLIM----P------------E
RYRNCIESLG-D--CL--GEGVLFILEGYDE---LPDEQRESN------WL---------
--------QT-LME--------E--IE----------EELPKASVMVTSR-P------WA
T-LGIGE--------FFQ-EHVQILGFTESSRREYVDSVLSSD

>AqNLRX4
--ILIE-GVSGIGKSTL-AYEMLKQWKD-------G-T--------------ALR-RYSY
V-LLLR-LRNEN--VHQQWSQ--SPD--------VA--QLIEE-CLN----------EQY

```
IEPPEIKSIL-D--NK--GHNLLLILEGYDE---LPKEKFEF-------QV---------
--------NVFQ----------K--LK----------SHFDNAVVIITTQ-P------SL
S-YQLTGNI------LFT-KTIEILGFN---------------

>AqueNLRX2
SWILIE-GVSGIGKSTL-AYEMLKQWKD-------G-TAL---------------QQYSY
V-LLLR-LRNEN--VHQQWSP--SPD--------VP--QLIED-CLN-E---------QY
IEPPEIKSIL-D--NK--GHNLLLILEGYDE---LPKGKFKFQ-----------------
---------MEVFR--------Q--LK----------SDFHNAVVAITIH-P------SF
L-YQLSENI------LFT-KEIEILGFNKSSQNRYIEIAFKND

>AqNLRD9
--ILIE-GVSGIGKSTL-AYEMLKQWKD-------G-T--------------ALQ-NYSY
V-LLLR-FRNEN--VHQYWSP--S-K--------VT--QLIQE-CLN----------EQY
HEQP---DIL-S--NS--GQDLLLILEGYDE---LPKKELEC-------HA---------
--------QVFY----------Q--LN----------RNFHNAVVVITTH-P------SF
S-YQLSDKI------LFA-KKIEIIGFDKPNQDQYIKVAF---

>AqueNLRX1
SWILIE-GVSGIGKSTL-AYEMLKQWKD-------G-TAL---------------QNYSY
V-LLLR-FRNEN--VHQYWSP--S-K--------VT--QLIQE-CLN-E---------QY
HEQP---DIL-S--NS--GQDLLLILEGYDE---LPKKELECH------A----------
--------QVFY----------Q--LN----------RNFHNAVVVITTH-P------SF
S-YQLSDKI------LFA-KKIEIIGFDKPNQDQYIKVAFKND

>AqNLRD1
--VLIE-GIPGIGKSTL-AYEMCKRWAD-------G-I--------------ALQ-KYTL
I-LLLR-LRDND--VQCNWSP--D-K--------VQ--KLIGI-YLD----------KQS
WKSEAVQKIF-D--GD--GEGLLIILEGFDE---LPED-KPS-------NV---------
--------EVLR----------V--IM----------RDMSEATIIVTTR-T------ST
T-HELSHNI------RFK-KHIEIQGFNQDNRNRYVKTFF---

>AqueNLRD4
DAILIE-GAPGIGKSML-AFEICSRWVK-------G-KAL---------------MGYTL
L-LLFR-LREKF--VQ----D--CET--------VK--ELLGC-FLV----------GQS
WKEEVVRDII-D--NS--GEGVIIILEGFDE---LPHELTTPD------S----------
-----------VFL--------K--LS----------FELPSASFIFTSR-P------SA
K-HCLKQEI------LFE-RHVEVIGFTKPSIEKYVTEFFKGN

>AqueDN4
EAVLIE-GAPGIGKSML-AFEICSRWVK-------G-EAL---------------KKYPL
L-LLLR-LRDRV--IK----N--CES--------VR--ELLGC-FLK----------EQS
WKDAVVQDII-D--NG--GEGLIVILEGFDE---LPEHLTKQD------S----------
-----------VFF--------Q--FS----------TVLPCASLVFTSR-P------SA
K-HFLRLKV------EFE-LHVEVIGFTHENINEYIQKFCKGN
```

>AqueNLRD1
DAILIE-GAPGIGKSML-AFEMCSRWVK-------G-EAL---------------KKYPL
L-LLLR-LRDRV--IQ----N--CKS--------VK--DLLGC-FLK----------EQS
WKDAAVQDII-D--NG--GEGLIVILEGFDE---LPEHLTKRG------S----------
-----------LFF--------Q--FS----------NELPCATLVFTSR-P------SA
K-HYLKNEI------EFE-RHVEVIGFKKENIDEYIEKFCNNN

>AqueNLRD2
EAVLIE-GAPGIGKSML-AFEICSRWIK-------G-EAL---------------KKYPL
L-LLLR-LRDKF--IQ----N--CKT--------VK--DLLGC-FLK----------EQS
WKEEAVQHIF-D--KS--GEGLIVILEGFDE---LPEDLTQAS------S----------
---------V--FL--------Q--IS----------EELPFTSLIYTSR-P------SA
K-HSLKQEI------SFS-RHIEVLGFTGKSINEYIQLFFEND

>AqNLRD4
--VLIE-GAPGIGKSML-AFEICSRWVK-------G-E--------------ALK-KYPL
L-LLLR-LRDRV--IQ----N--CKS--------VR--ELLGC-FLT----------EQN
WKDATVQHIF-N--KS--GKGLIVILEGFDE---LPEDLTQ--------PG---------
--------SVFL----------N--IS----------EELPFASLIYTSR-P------SA
K-HYLRQEI------TFS-HHIEVIGFTSKSISQYIQIFF---

>AqueNLRD3
EAVLIE-GAPGIGKSML-AFEICSRWVK-------G-EAL---------------KKYPL
L-LLLR-LRDRV--IQ----N--CKS--------VR--ELLGC-FLT----------EQN
WKDATVQHIF-N--KS--GKGLIVILEGFDE---LPEDLTQ--------PG---------
--------SVFL----------N--IS----------EELPFASLIYTSR-P------SA
K-HYLRQEI------TFS-HHIEVIGFTSKSISQYIQIFFQDN

>LchaCN7
SMVTVQ-AAPGMGKTFTFAKMLPLRWLK-D-----P-EFW---------------PKFDL
V-FVVH-VGDPA--VYN------ATT--------LE--EFLL--GSY--------HLVDK
T-KRAVLDFV-E--TS--PGRVLVICDALDE---GHDHI-----------S---------
--------AEVKTI------L----KG----------ELLPGLHLLITSR-P------CQ
GFHELAE--------HAH-RQLELLGIPNDKLHEFVLKQFSGD

>LcomNLRX20
RNILAA-GPPGSGKSYLFTKIIPFLWSC-------D-LLW---------------KGRFD
LVAFFD-LCRED--VRN------AAD--------MK--ELLAT-FTEDT------MMDDG
ERSAAYFHSA-R--MM--AERLCLIFDGLSE---CSVAD---C-------S---------
--------EFIRGI------L----GR----------TRWSACHVIVLTR-H------LT
DATIVDKRF------QYH-RFLEVSGLTRCAMDGIITSRVSAD

>LcomNLRX21
RNILAA-GPPGSGKSYLFTKIIPFLWSC-------D-LLW---------------KGRFD
LVAFFD-LCRED--VRN------AAD--------MK--ELLAT-FTEDT------MMDDG
ERSAAYFHSA-R--MM--AERLCLIFDGLSE---CSVAD---C-------S---------

--------EFIRGI------L----GR----------TRWSACHVIVLTR-H------LT
DATIVDKRF------QYH-RFLEVSGLTRCAMDGIITSRVSAD

>LchaCN1
RRQIAT-GCAGAGKTTAFTLKAPYEWAKEG-----S-DFW---------------QQFRL
F-FFGS-LNDTK--WRD------SKS--------LN--DVFGL-GEF--------GLTKR
A-QTDVLTFI-R--NH--SEKVLLVADSFEE---APQLD---H-------S---------
--------SLLWMV------LSG--KE----------L--PQLNVMVSSR-P------CK
MASWLSRNF------PFH-QRLEVMGFTLEKILLYVKEYFSHD

>LchaCN8
RRLIAT-GCSGAGKTTAFTIKAPYEWAKEG-----S-DFW---------------QQFKL
F-FYGS-FNDTK--WRD------AES--------LA--DVFHL-NEF--------QLTEA
E-QKEVLAYI-R--EH--PDKVLLVADSLEE---APEPK---Q-------S---------
--------SMLLKV------LSG--KE----------RGLLNLNVVVSSR-P------CR
ITAVLSKDC------PFD-QRVEVAGFTPDKIKRYVEWFFSQD

>ScilNLRX10
---IAV-ASAGCGKTFAFTKVAPLKWAM-------G-KLC---------------QRKKL
L-IARE-LHHED--VKM------AKS--------LS--GLLGL-EGI--------GIEDS
HDRQVICEYV-R--AQ--PDALCLVLDGLDE---INLSE---C-------S---------
--------SFVQGV------I----QG----------EELPGVHLIVTSR-P------CP
DVFSLSAMP------HFQ-QHVELVGFQPDDVQMYVNKVLRSE

>ScilNLRX11
---IAV-ASAGCGKTFAFTKVAPLKWAM-------G-KLC---------------QRKKL
L-IARE-LHHED--VKM------AKS--------LS--GLLGL-EGI--------GIEDS
HDRQVICEYV-R--AQ--PDALCLVLDGLDE---INLSE---C-------S---------
--------SFVQGV------I----QG----------EELPGVHLIVTSR-P------CP
DVFSLSAMP------HFQ-QHVELVGFQPDDVQMYVNKVLRSE

>ScilNLRX12
---IAV-ASAGCGKTFAFTKVAPLKWAM-------G-KLC---------------QRKKL
L-IARE-LHHED--VKM------AKS--------LS--GLLGL-EGI--------GIEDS
HDRQVICEYV-R--AQ--PDALCLVLDGLDE---INLSE---C-------S---------
--------SFVQGV------I----QG----------EELPGVHLIVTSR-P------CP
DVFSLSAMP------HFQ-QHVELVGFQPDDVQMYVNKVLRSE

>ScilNLRX13
---IAV-ASAGCGKTFAFTKVAPLKWAM-------G-KLC---------------QRKKL
L-IARE-LHHED--VKM------AKS--------LS--GLLGL-EGI--------GIEDS
HDRQVICEYV-R--AQ--PDALCLVLDGLDE---INLSE---C-------S---------
--------SFVQGV------I----QG----------EELPGVHLIVTSR-P------CP
DVFSLSAMP------HFQ-QHVELVGFQPDDVQMYVNKVLRSE

>LcomNLRX22

---------AGCRETELFLTKSPRDWAI-------G-KLW---------------REFDL
L-VARE-LRSDS--VHK------ARN--------VS--DLFAL-EYY--------GVRSL
EEQQIVSKFV-Q--RN--PARVCLILVGLDE---IQMSE---C-------S---------
--------SFMQQV------I----NG----------ETLRGIRLLLTSK-P------SP
EVFKLSVSS------PFD-RHIEVAGFLPQNTRNYICKALSPN

>SciINLRX15
RRVLAV-ASAGCGKTVLFTIKIPHDWAS-------G-DLW---------------AAHFD
LLCVVQ-LNDVA--ARS------AHN--------TE--ELLQL-ATL--------GLTAA
E-QAELAQYV-H--AH--PERVCLILDGLDE---CHVYE---C-------S---------
--------LFVQQV------IYD--QC----------PCLAGMRVIITSR-P------CI
ATSVLTQRA------RVD-RRLEIIGFTHSSVYDFVRKYLTGE

>LchaNLRX2
TRVLAI-GTAGSGKSTAFTVKAPHDWSL-------G-QLW---------------PETVL
FQCL-K-LRDRS--VWK------AQS--------VS--ELFQL-NAL--------GLNDA
E-RTEVENFI-V--KN--PGRVLLACDGLDE---CVVQE---F-------G---------
--------LLWSV---------L--QG----------KALAGVRLLMTSR-P------CE
LLIDLSRET------AID-SHVRLFGFTEENVLVFVDRYVGGA

>LchaNLRX15
---------------SRFCFRVLLQWAC-------Y-KVF---------------RQFAI
V-FYIS-MRDHE--RSC------TTS--------VA--SLL----RLDV------MGFKS
PEQRSILAYL-N--KE--SHKVLLIVDSCTH------DISQQQ-------G---------
--------SAVQQL--------F--DG----------VLFPNASIIISAQ-P------SL
SLLQLTARC------ERH-YCFQ--------------------

>LchaNLRX1
-RCLLL-SPPGGGKTMT-CHQILNLWAG-------G-KSF---------------KHFKA
V-VYLT-GKEDQ--RVC------TSK--------VG--EFLAL-NDSEP------TDRKS
LTERLIQSHV-A--SH--GEELLIILDAADE---SSSDSLFKD-------G---------
--------GILATLFSPAKSLSS--GL----------GKLTDASVIVTSR-P------CP
ASDYLVQGG------DCT-AVLWLCGFTDYNLQQLLRKRLGEE

>LchaNLRX8
-KSLVI-GVPGAGKSRL-CLELCERWAK-------G-QMM---------------RNIRF
P-FLLP-ARELT--RAL------ATCNFSADNPNWL--GFLCM-SELAI------GGLSN
EEQNIAWRYM-R--SH--SEEVIIILDGLDE---LDTDIVDSQ-------N---------
--------SYLYQL--------V--HR----------HVLPSCSLLITSR-P------CE
KALELVRWT------D---KHYQIKGISRKQLRGLMEEWLGEE

>SciINLRX5
TKCLLI-GPPGAGKTMT-SLEILRRWGC-------C-DMF---------------TEYSL
A-LYVA-ARRLR--ED-------MTF--------W---DVLGL-G-TAA------YGLSA
ADMAEVRDYA-T--QH--SKQILVVMDGLDE---LRPSLIGEE-------S---------
--------AVLRVL--------A--GT----------GELSACHVVATSR-P------CA

QARRLLSNT------A---MHYQMCELSSQQQTAMLRRRIQQR

>LcomNLRX17
SRCLLI-GPPGAGKTMT-SLELLRRWSS-------C-TMF---------------TEYYL
A-LYVP-ARALC--EEV------QSV--------W---DLLRL-A-SPC------YGLTQ
GELEDVQQYI-L--DH--SQQLLLVLDGLDE---VPVRLISGT-------S---------
--------TFSM-L--------L--HG----------NPLTSCHIVATSR-P------CK
QALVLLRSV------E---IRYQICDLSQVQRDAMLKSRLNG-

>LcomNLRX28
SRCLLI-GPPGAGKTMT-SLELLRRWSS-------C-TMF---------------TEYYL
A-LYVP-ARALC--EEV------QSV--------W---DLLRL-A-SPC------YGLTQ
GELEDVQQYI-L--DH--SQQLLLVLDGLDE---VPVRLISGT-------S---------
--------TFSM-L--------L--HG----------NPLTSCHIVATSR-P------CK
QALVLLRSV------E---IRYQICDLSQVQRDAMLKSRLNG-

>LchaNLRX11
QRCLML-GQAGSGKTMA-CRKLANSWAQ-------G-KGW---------------RDYTL
V-IYVA-ARELT--RHS------GTID-------MQ--CLLGI-SRTPK------LS--D
KEKSYVWQHI-Q--EN--EERVIVIFDGIDE---GAGSLLK---------L---------
--------EAFQQI--------V--KR----------QSLARCTVLMTSR-P------CG
AAFQFVDKV------D---QFVELVGFPKESLNLMIAKKLTTD

>SciINLRX19
RQYLVY-GPPGVGKTVALSYKLPFEWAT-------E-KAL---------------KIYKF
V-IVVP-LRESH--LAA------ELDP-------AI--LLL---SRISR------LREDA
ELRHKIYKVL-E--RE--EKQLLIVFDGLDE---VQNPNQ----------K---------
--------AAFMRI--------L--HG----------DLLPMCTVLVASR-P------CR
LASEYATQA------D---QCVEVIGFNDDDVRLFVESRFIRS

>LcomNLRX19
KRYLVY-GPPGVGKTVALCYKLPYEWSV-------M-RAL---------------KTYMF
V-IVVP-LREVK--LAQ------ETDP-------AK--LLL---SRIGP------VARKP
HLLRGLYERL-C--DN--EYRLLIVFDGIDE---VRGASP----------D---------
--------SAFVRI--------M--YS----------QLLPKATVLVSGR-P------CK
LASEYGAVA------N---QCVEVIGFNDQDVDLFIRHRFTKD

>LcomNLRX25
KRYLVY-GPPGVGKTVALCYKLPYEWSV-------M-RAL---------------KTYMF
V-IVVP-LREVK--LAQ------ETDP-------AK--LLL---SRIGP------VARKP
HLLRGLYERL-C--DN--EYRLLIVFDGIDE---VRGASP----------D---------
--------SAFVRI--------M--YS----------QLLPKATVLVSGR-P------CK
LASEYGAVA------N---QCVEVIGFNDQDVDLFIRHRFTKD

>LcomNLRX26
KRYLVY-GPPGVGKTVALCYKLPYEWSV-------M-RAL---------------KTYMF

V-IVVP-LREVK--LAQ------ETDP-------AK--LLL---SRIGP------VARKP
HLLRGLYERL-C--DN--EYRLLIVFDGIDE---VRGASP----------D---------
--------SAFVRI--------M--YS----------QLLPKATVLVSGR-P------CK
LASEYGAVA------N---QCVEVIGFNDQDVDLFIRHRFTKD

>LcomNLRX27
KRYLVY-GPPGVGKTVALCYKLPYEWSV-------M-RAL---------------KTYMF
V-IVVP-LREVK--LAQ------ETDP-------AK--LLL---SRIGP------VARKP
HLLRGLYERL-C--DN--EYRLLIVFDGIDE---VRGASP----------D---------
--------SAFVRI--------M--YS----------QLLPKATVLVSGR-P------CK
LASEYGAVA------N---QCVEVIGFNDQDVDLFIRHRFTKD

>jgi_Nemve1_204364
-----E-------------------------------------------------DVKV
V-LLLQ-CRDL----VDVSDW---KD--------VI--RQAI------P------EHYSE
QDQESIIRYI-C--DN--EEHIAFVLDGYDE---LPSSSKG------------------
---------PLDMI--------IN-AK----------KVLKQCYVIVTSR----------
M--NQIPQEL---KNCFD-RTLEIKGWDLSNAEQFIKDY----

>SPU_014719
-LAFI--GEAGVGKSTL-FAKIALDWAL---------------------------QEINL
L-FLES-FREI----KESGFFG--DT--------VM--GHFP------D--------DSE
VNGEWIDEYI-R--KN--QRKVLILLDGLDE------AQIDIKKPNRND-----------
---------AIVSI--------V--RG----------ERFIDTPVVITTR----P-----
-------------------------FGADQIKS---------

>SPU_013619NLRD
--VVAR-GRTGSGKTTF-LLKLSSDWAK----------------------------DAVA
V-FLLQ-LRKL----DHTSNFG--AA--------VV--DQLL------P--------KKD
FTPQFIEEFA-E--KN--QKRVVVLLDGYDE---FKGKGLDQKNCG-------------
---------NIVKM--------L--RK----------EYLPFVQILITTR-P--G---RV
G--DFIKLEG---HFTNEYRHLQITGFSSQDIDAYVKKI----

>SPU_003539
--ILIS-APAGRGKTTA-VAKMAYDWVL---------------------------KHLPL
L-FVVK-FRNT----SQLTSIG--EA--------IK--SQLL------N-------DVDD
LTPEGLESFI-R--AN--QEICHIILDGLDEYAGISSSEQSSRS----------------
---------NIVSV--------I--RY----------EEFTECRVVVTTR----P---HL
E--NFFNQAE---LPRVY-TKMWIEGFSRESSRDYIDKF----

>jgi_Nemve1_208313
--VLIE-GESGAGKTTL-CQKLAFDWA-----------------------------EVEL
L-LFLRCKEM------KDGGIA--KA--------MQ--EQLL------S------SETSD
EERKQVFDYM-K--EN--QSKILMILDGYDE---LPVEASNVKE----------------
---------RLGAV--------I--TR----------KAFATSNVLVTIR--------SQ
E--ARPEQIS--------------------------------

>jgi_Nemve1_203213_fgenesh1_pg_scaffold_41000052
--VLLE-GESGVGKTTF-CQKLAYDWA-----------------------------QVEI
L-LSLK-SKGM-----SGGGLW--KA--------IS--EQLV------P------EETND
DEREELFHYI-K--EN--QEKVLIILDGYDE---LPSHASVKE-----------------
---------SINKV--------V--SK----------KALPFSYTLITSR----P-----
E--KRLSKYF------IGGNVLQVKGLR---------------

>adi_v1_01267NLRG
--VLIE-GEPGMGKTTY-CQKLVFDWAP----------------------------RIDI
F-LLLR-CRGI------KSTIW--DA--------IE--DQIL------P------DEINP
QDKKIFFQFL-R--EN--PSKVLLVLDGLDE---ADPQKLD-------------------
---------MYLKL--------V--QS----------KQLFGCYIVLTSR----H---EA
G--VKVRPYT--------DTLLEIVGFTVSDAKCYIRKYVHIE

>adi_v1_00412
--VLVE-GSPGIGKSTF-CLKLAHDWAP----------------------------SFKL
V-FLLK-CRDM------KGDIV--ED--------IF--EQLL------P------EDLKE
KTKEVLVNFL-GDLNN--QKQILIILDGLDE-----------------------------
------------------------------------------------------------
-------------------------------------------

>pfu_aug1_0_7988_1_38671_t1
--IFLY-GDAGSGKTCY-AHRMASLWAN----------------------------EIDY
V-FFIS-MKYL---SGDDETVE--DI--------VC--NDFLF------------SQYDE
NARSTARKAL-R--SP--QTRCLVIFDGLDE---WPKN----------------------
-------------K--------LP-ST----------GNLRNCQVLFTSR-P--SKIATM
N--LRY---------EET-------------------------

>pfu_aug1_0_1515_1_29740_t1
--VLLI-GLPGRGKTTL-CHWLAQNWCR---------------------------SKFDF
L-FLIK-LCEI---SKTMSSLQ--EI--------IC--KQLFK--------------NNQ
KCHDIVHEIL-S--SP--DYRCLIIFDGLDE-----------------------------
----------LKSG--------IKLPI----------TNFPSCTTLITTR-P--VRRY--
-------------------------------------------

>pfu_aug1_0_1515_1_29739_t1
--IFMI-GHPARGKTTY-CQWLSHNWC-----------------------------EFDF
L-FFVK-LRNI----ATFDSLA--NA--------IW--YDVFK--------------DDP
VTAHDVNEVL-M--NA--NERCLIILDGLDE---LKSEC---------------------
----------TCLD--------I--EM----------SSIRNCVTFFTTR-P--WRYADV
S--HLV---------DDNDKVLEIKGLDDEGIRNVTEKV----

>pfu_aug1_0_16419_1_54300_t1
--VYML-GDAAHGKTTY-CMWLINTWC-----------------------------QFDF
V-FYLQ-FRHV---DPSTTNVN--DM--------IC--KYILG--------------RDT

TLYEVVKNVL-F--CD--RYKCFILMDGMDE---QKSKK---------------------
-------------G--------MP-YI----------DNMSNCVLFIPMR-P--WRFADV
T--EFV---------NYNDRVVEILGLDDKSIEAVIQKV----

>pfu_aug1_0_188_1_14872_t1
--IFMH-GEAARGKSMY-CKQLLNYWC-----------------------------EFDF
T-FLVS-LRHV---HQSRTLLE--DM--------IC--YDTFK--------------NNK
ECHDTIRHVL-S--SD--QYRCLILVDGLDE---WRISKDKSSEL---------------
----------THNG--------LP-NI----------ETFSACTCFFTLR-P--SRLDVV
A--SAI---------YDSKKVVQIFGLNEKGQNLVVDKV----

>pfu_aug1_0_128_1_14815_t1
--IFMT-GDSGRGKTTF-CLWLLKNWCK----------------------------EFDF
C-FLLQ-LGDFQ--DVQINVASVKDM--------IC--QVVFE--------------SDT
DHHATIRHVL-S--SN--QYRSLIIIDGLDE---WNPNEETRKHL---------------
----------IYDR--------MP-KL----------DVSSTVSCFIAMR-P--WVLSTM
S--KTV---------RNSDRVIEVFGLNEAGIENVVKNV----

>gi_2642132_gb_U19251_1_NAIP
-VMCVE-GEAGSGKTVL-LKKIAFLWAN----------------------------RFQL
V-FYLS-LSST---RPDEGLA---SI--------IC--DQLLE-------------KEGS
VTEMCMRNII-Q--QL--KNQVLFLLDDYKEICSIPQ-----------------------
---------VIGKL--------I--QK----------NHLSRTCLLIAVR----T-----
---NRARDIRRYL-----ETILEIKAFPFYNTVCILRKL----

>AF376061_1_IPAF_hsNLRC4
-CIIE--GESGKGKSTL-LQRIAMLWGT----------------------------KFKF
V-FFLR-L-----SRAQGGLF---ET--------LC--DQLLD-------------IPGT
IRKQTFMAML-L--KL--RQRVLFLLDGYNE---FKPQNCP-------------------
---------EIEAL--------IK-EN----------HRFKN-MVIVTTT-T--------
---ECLRHIR--QFGALT---AEVGDMTEDSAQALIREV----

>Lotgi1_152683NLRD
--VILE-GESGSGKSTL-AAMMTYQWAS---------------------------EYYRF
M-ILVD--AELL----EG-------N--------VR--KSIYQQTIL---------ESSK
ITFDEFWETL-E--SY--DEDVVLVIDGFTG------GRS--------------------
---------ELCKL--------I-DGT----------HLLKSAVLVMVS--P--------
----------------------NIS------------------

>CGI_10018771NLRC
--ILME-GPSGSGKTIL-SHMVAYMWAK---------------------------NKYGI
L-LHVD-LKRI----NGD----------------FQ--EQV--HRNLLP-------DDFK
LNPSEFFSML-E--AN--ASEVVMIVDGYDG------KPNQK------------------
---------VLEDI--------L--SG----------SQLRQANVIVMMN----P-----
-------------------EIVSSPGFLPDSRMFS--------

>Aqu1_2238711455876_AqNLRX1
--ILIS-GRPGAGKTVL-LTKVAKDWAH----------------------------DVNL
L-LHIS-LREL--ARKSDFP-----Q--------LE--DAVS-----------LCL-LDK
EKVHELAKVIDE--VN--GKGICFAFDGLDE---YPKR----N--DPSD-----------
---------AIMKI--------I--RK----------EFLPLATVIVSSR----P---TA
S--SCVPVQS-------RDLHVEIIGFMPEQIKEYVTQY----

>Aqu1_211634_AqNLRX51
--VVID-GPPGIGKTTL-CRKLLNMWSH---------------------------QQYDL
V-LYCP-LRNS--KIAT------ATT--------LA--DLFE---------------YKS
SKISKVVDWFSN--GD--GEGLLIMFDGWDE---LSEQ----L--RQSS-----------
---------LAASI--------I--QR----------KELDQCSVVVTSR----S---YA
S--SSLLKMD-----TLS-RHVQVIGFSKEEISTVIIQT----

>Aqu1_208189_AqNLRX20
--VVID-GPPGIGKTTL-CRKLLNMWSH---------------------------QQYDL
V-LYCP-LRNS--KISA------ATT--------LA--DLFV---------------RKL
KRYKNVPEWFEE--RD--GEGLLIIFDGWDE---LSET----L--RQSS-----------
---------LVASI--------I--CK----------DELDQCSVIVTSR----S---YA
S--FSLFEVM----SNLS-KHVQVIGFTEEEISTVIIQT----

>Aqu1_204933DN_AqDN2
--VLIE-GEPGIGKSTL-AKELALRWAN----------------------------NYDI
V-FLIQ-LRFE--TYHK------ATS--------VE--DLLV-----------DLD-DQS
INMTDLHVEIRK--RK--GAGILWILDGFDE---LPSH----L--KNNSS----------
---------VLMKL--------I--KG----------DILPKSTVIVTSR----P---VA
S--DLLLHFL---HDHNS-KRISLRGFDSTKIEEYASKY----

>Aqu1_220887DN_AqDN10
--VLIE-GEPGIGKSTL-AKELALQWAN----------------------------NFKI
I-ILIP-LRLE--IYQK------AEN--------IE--DLLI-------------Y-VED
IDMVKITSSINR--AR--GAEVLWILDGFDE---LPHH----L--RNSSTS---------
---------IFIKL--------I--KG----------DILPKSTVIVTSR----H---AA
T--DRLLTFL----EDDS-KHIVLRGFGPNEILEYVSKY----

>jgi_Capca1_219741
--ILVE-GEAGIGKTTF-LQMLVSEWNH----------------------------SFDL
L-FKLH-AKDF--IGC--------TS--------IA--DVIK--SCLLA-------RDSE
ISTETLEAILQ-------KHNVIILVDAYDE------GHVENH-----------------
---------LLNDV--------I--EG----------RVLKDATVLLSSR----P----A
F--LKFRFFD---------SIVVVQGFDEEHQIEYVDR-----

>jgi_Capca1_205954NLR
---------PGIGKSIL-CQFFAYEWS-----------------------------KFDL
V-FYLK-AEDL--KHQ--------DS--------IA--EAIR--CHLLP-------EDFK
MSPNNLEELLQ-------TTTVLFLIDAYDE------ACVDNL-----------------

---------LLEKL--------I--EK----------KHLRKSSLLVTSR----------
------RHFL---------------------------------

>jgi_Capca1_203863
--LLIE-GNPGNGKSTI-CDWLAYKWS-----------------------------SFDL
V-IYLH-AKHL--KDQ--------TT--------IA--DAIK--SHLLP-------EDFD
ISSCRLTEVL-K------AKYVLFIVDGFDE------ACSENP-----------------
---------LLEQL--------I--ER----------KILRHTNILLTSR-P--G---FL
Q--EKLKYFD---------LIFEVEDYDEQQQ-----------

>jgi_Capca1_186473NLRCas
--ILVE-GDPAQGKSTL-CQALAYAWSK----------------------------SFDL
V-ILLH-AGDL--RGQ--------DS--------VA--EAIK--THLLP-------KDCD
ITSGQVNGLLLA------K-NVLLIIDAFDE------AANENE-----------------
---------ILHQL--------I--EG----------KLLKHKTLLLTSR-P--N---FL
K--NKLLHFH---------STATVEGYNEKEQLEHVRRY----

>jgi_Capca1_192408NLRC
--IMVE-GNPGVGKSTL-CQKLAFAWSH----------------------------SFDI
V-IYLT-AADF--KGF--------KD--------VA--SLIR--AHLLP-------KDFR
ISVNAISSALR-------DRCVLFLVDGYDE------SYSDNP-----------------
---------LLHDV--------I--QR----------KVSVQSTILLTSR-P--N---YL
R--EMLRYFD---------TRLHLDGFNAQQRQNYIEKF----

>jgi_Capca1_193620NLRC
--ILVE-GDAGMGKTTL-CQNLAFRWAH----------------------------SWTF
V-IYLT-AANL--QGY--------DD--------IH--TAVH--DHLLP----------G
VSNEEVKKALD---SS--QSSTLFIIDSFDE------GYMDNM-----------------
---------LLRDL--------I--QG----------RVYSNATLLLTSR-P--N---YL
Q--DLLKEFD---------SKLSTDGFSTEQKLLYIKKF----

>gi_28866920_gb_AF389420_1NOD4
-TVLL--GKAGMGKTTL-AHRLCQKWA-----------------------------CFQA
L-FLFE-FRQL--NLITRFL-----T--------PS--ELLF-------D---LYLSPES
DHDTVFQ-YL-E--KN--ADQVLLIFDGLDE-----ALQPM-GPDGPGPVLT--------
----------LFSH--------LC-NG----------TLLPGCRVMATSR-P--G---KL
P--ACLPAEA---------AMVHMLGFDGPRVEEYVNHF----

>jgi_Capca1_214069_NLRC
--ILLI-GRPAIGKTCF-LRKTCQDWAP---------------------------ENFDC
V-FLFQ-LREF---NQDDILK-KGIS--------LR--KLLM-------E---RHGPEKP
DEDIIWN-QL-K------PNKCLIIFDGLDE---FKGFEEL-PPKDPPKYSS-SSARMRL
PF-------LLWNL--------L--KG----------NIFKGCYVICSSR-P--R---AL
A--LPLLQDL------FQ-RNVDLHGFDEAQVVAYFQ------

>jgi_Nemve1_209080

--VLVV-GRPGIGKTMM-CTRLLRLWA-----------------------------SSDE
V-ALSIAFRNV---SSTS-----NIN--------IR--DLLT-LST---T---TQGLDDN
EISYILN-YI-T--AN--PSKLILYLDGVDE---FSSRSNI-SQSEDSERST--DGKMAP
HI--------LFRK--------IW-SR----------ELLDKATVITTTR----P---TA
V--ECLRNLP---KDCLN-RTVEILGFSFKQVKDYVTKF----

>adi_v1_15689
--VLVV-GRPGIGKTSF-STKFLRLWAD---------------------------KHFNF
A-FLLK-FRRF--NNENA-----NLS--------LR--DLMA-----------RSETVQS
LDDAVWD-FI-K--QE--PTKVLLIFDGLDE---YSRKEDI-VVHHDSPYKN--CVEEKM
PLSV------LYNK--------LA-EG----------KLLPGASILTTTR----P---TA
V--KCVLHLR------FQ-RTVEIRGFTSDDVKEYVKNF----

>jgi_Capca1_207210_NLRC
--VLLQ-GSSGAGKSTL-CASVAYAWSK---------------------------TPFRL
A-FYVD--LKMA----KGGLL---EC--------IQ--QQCL----L-------------
QEQQDLDALI-E--NN--QRDSLFLLDGYEN---ISNPDS--------------------
---------DIYDL--------I--HR----------RLYPKSTVLLTTE-S--T---FL
T--RPLAKLF--------DMRLVLSGISPESQGALIQHY----

>gi_836908_gb_U18259_1_CIITA
--IAVL-GKAGQGKSYW-AGAVSRAWA----------------------------PQYDF
V-FSVP-CHCL--NRPGDA----Y-G--------LQ--DLLF---SLGPQP-------LV
AADEVFSHIL-K---R--PDRVLLILDAFEE---LEAQ----DGFLHSTCG--PAPAEPC
SLRG-----LLAGL--------F--QK----------KLLRGCTLLLTAR----PR-GRL
V--QSLSKAD---------ALFELSGFSMEQAQAYVMRY----

>AF126484_1_HsNLRC1
--IFIL-GDAGVGKSML-LQRLQSLWAG---------------------------VKFFF
H-FR---CRMF--SCFKESD---RLC--------LQ--DLLF-------KH--YCYPERD
PEE-VFAFLL----RF--PHVALFTFDGLDE---LHSD----LDLSRVPDS--SCPWEPA
HPLV-----LLANL--------L--SG----------KLLKGASKLLTAR--------TG
I--EVPRQFL--------RKKVLLRGFSPSHLRAYARRM----

>HsNLRC2_NACHT_trunc
--VLV-VGEAGSGKSTL-LQRLHLLWAA---------------------------GQFVF
P-F---SCRQLQCMA-KPLSV--------------R--TLLFEH---------CCWPDVG
QE-DIFQLLL-DH-----PDRVLLTFDGFDEF-----K----FRFTDRERH--CSPTD--
--PT-SVQTLLFNL--------L--QG----------NLLKNARKVVTSR-P-------A
A--AFLRKYI---RTEFN-----LKGFSEQGIELYLRKR----

>gi_30348947_tpg_BK001112_1_NOD3
--VSITIGVAGMGKTTL-VRHFVRLWAG---------------------------KDFSL
V-LPLT-FRDL--NTH-------EKL--------CA--DRLI--------C--SVFPHVG
EPSLAVA----V------PARALLILDGLDE---CRTP----LDFSNTVAC--TDPKKEI
PVDH-----LITNI--------I--RG----------NLFPEVSIWITSR----P---SA

S--GQIPGGL------VD-RMTEIRGFNEEEIKVCLEQM----

>AB094095_1_NOD5
--VVLY-GTVGTGKSTL-VRKMVLDWCL---------------------------PAFEL
L-IPFS-C-----EDLSSLGP-APAS--------LC--QLVA---------------QRY
TPLKEVLPLM-A--AA--GSHLLFVLHGLEH---LNLD----FRLAGTGLC--SDPEEPQ
EPAA-----IIVNL--------L--RK----------YMLPQASILVTTR----P---SA
I--GRIPSKY------VG-RYGEICGFSDTNLQKL--------

>gi_19387135_gb_AF479748_1_NALP6
--VVLQ-GPAGIGKTMA-AKKILYDWA-----------------------------QVDF
A-FFMP-CGEL--LERPGTR-----S--------LA--DLIL-----------DQCPDRG
APVPQ---ML-A---Q--PQRLLFILDGADE---L-----PALGGPEAAPC--TDPFEAA
SGAR-----VLGGL--------L--SK----------ALLPTALLLVTTR----A---AA
P--GRLQGRL------CSPQCAEVRGFSDKDKKKYFYKF----

>gi_28436379_gb_AY154468_1_NALP13
--IVLV-GRAGVGKTTL-AMRAMLHWAQ---------------------------QRFSY
V-FYLS-CHKI--RYMK------ETT--------FA--ELIS-----------LDWPDFD
APIEE---FM-S---Q--PEKLLFIIDGFEE---IIISESRSESLDDGSPC--TDWYQEL
PVTK-----ILHSL--------L--KK----------ELVPLATLLITIK----T---WF
V--RDLKASL------VNPCFVQITGFTGDDLRVYFMRH----

>gi_19387133_gb_AF479747_1_NALP4
--VIIQ-GPQGIGKTTL-LMKLMMAWSR---------------------------DRFLY
T-FYFC-CREL--RELP------PTS--------LA--DLIS-----------REWPDPA
APITE---IV-S---Q--PERLLFVIDSFEE---LQGG----LNEPDSDLC--GDLMEKR
PVQV-----LLSSL--------L--RK----------KMLPEASLLIAIK----P---VC
P--KELRDQV-----TIS-EIYQPRGFNESDRLVYFCCF----

>gi_28436371_gb_AY154464_1_NALP9
--VVLE-GPDGIGKTTL-LRKVMLDWAK---------------------------DRFTF
V-FFLN-VCEM--NGIA------ETS--------LL--ELLS-----------RDWPESS
EKIED---IF-S---Q--PERILFIMDGFEQ---LKFN----LQL-KADLS--DDWRQRQ
PMPI-----ILSSL--------L--QK----------KMLPESSLLIALG----K---LA
M--QKHYFML------RHPKLIKLLGFSESEKKSYFSYF----

>gi_28436373_gb_AY154465_1_NALP10
--VVLQ-GSAGTGKTTL-ARKMVLDWAP---------------------------GRFDY
V-FYVS-CKEV-------VLL-LESK--------LE--QLLF-----------WCCGDNQ
APVTE---IL-R---Q--PERLLFILDGFDE---LQRPFEEKLKKRGLSPK--ES-----
---------LLHLL--------I--RR----------HTLPTCSLLITTR----P---LA
L--RNLEPLL------KQARHVHILGFSEEERARYFSSY----
